# Faster adult implicit probabilistic statistical learning following childhood adversity

**DOI:** 10.1101/2025.10.15.682535

**Authors:** Bence C. Farkas, Bianka Brezóczki, Teodóra Vékony, Pierre O. Jacquet, Dezso Nemeth

**Author notes:** Corresponding Author: Bence C. Farkas, UVSQ, Inserm, Centre de Recherche en Epidémiologie et Santé des Populations, Université Paris-Saclay, 78000 Versailles, France.

## Abstract

According to deficit models, early life adversity disrupts normal development, leading to long-term emotional, behavioural, and cognitive difficulties. However, some evidence suggests that certain psychological skills may be preserved or even enhanced by early adversity. We hypothesised that implicit learning and memory would be equally effective in individuals exposed to childhood adversity and those from more favourable backgrounds, and compared the effects of childhood versus adult adversity. To this aim, retrospective childhood harshness and unpredictability measurements and current perceived socio-economic status were collected in a sample of 325 participants at a Hungarian university taking part in an online experiment. They also completed a task allowing the assessment of multiple components of implicit statistical learning, including initial acquisition of regularities, consolidation of established regularities, resistance of established regularities against interference, and acquisition of novel regularities. Results showed that although statistical learning reached the same eventual level, its pace was quicker in individuals with relatively greater early life adversity exposure. Conversely, lower current socio-economic status was linked to reduced learning performance. These findings partially support the ‘hidden talents’ framework, suggesting that early adversity may promote certain adaptive cognitive skills.

## Introduction

Childhood adversity refers to negative environmental experiences, such as poverty, neglect, or maltreatment, that require significant adaptation by a typical child (Ellis, Sheridan, et al., 2022; Frankenhuis & Amir, 2022; McLaughlin et al., 2021; Sheridan & McLaughlin, 2014). An extensive body of research has demonstrated that growing up in such adverse environments can hinder children’s development and lead to worse outcomes in physical (Felitti et al., 1998; Hughes et al., 2017) and mental health (Kessler et al., 2005; McLaughlin et al., 2010), academic achievement (Chang et al., 2019; Yeo et al., 2024). They also negatively impact various cognitive abilities, such as executive functions (Johnson et al., 2021; Lund et al., 2022), which refer to the collection of top-down mental processes required for goal-directed control of behaviour, and include working memory, cognitive flexibility, and inhibition (Diamond, 2013; Miyake et al., 2000). These deficits are apparent in both the behavioural and neural level (McDermott et al., 2012) and emerge early (Hostinar et al., 2012). While such a deficit-oriented approach has been crucial in drawing attention to the pervasive negative effects of early life adversity and informing interventions and public policy, it has also inadvertently biased our understanding of early life stress by focusing only on negative outcomes. An emerging ‘hidden talents’ framework is aiming to rectify this bias and identify psychological abilities that might be preserved or even enhanced by childhood adversity (Ellis, Abrams, et al., 2022; Frankenhuis, Young, et al., 2020). The framework extends earlier evolutionary-developmental models and focuses on contextual adaptation. It argues that childhood stress does not merely lead to dysregulation but induces a set of adaptive developmental processes selected to optimize the organism’s biological fitness, and can include physiological (e.g., hormonal changes, earlier puberty initiation), behavioural (e.g., more risk-taking and aggressive behaviours), and psychological changes (e.g., hypervigilance to threat, increased set-shifting ability) (Del Giudice & Ellis, 2016; Ellis, Sheridan, et al., 2022; Frankenhuis & Del Giudice, 2012). Some models also assume that these responses are part of a broader ‘accelerated life history strategy’ phenotype, characterized by a faster body growth, an earlier reproduction, a poorer somatic maintenance and, ultimately, a shorter lifespan (Belsky et al., 2012; Chang et al., 2019; Del Giudice, 2020; Ellis et al., 2009). Empirical tests of this idea have indeed found intact (social reasoning in Frankenhuis, Vries, et al., 2020; cooperative behaviour in Lettinga et al., 2021) and even enhanced cognitive skills in individuals who have grown up under more adverse conditions (sensitivity to social influence in Jacquet et al., 2019; set shifting in Mittal et al., 2015; working memory in Young et al., 2018; attention shifting in Young et al., 2022). In line with this reasoning, we propose that contrary to the pervasive negative effects on executive functions and explicit learning and memory, adversity exposure might preserve and even enhance implicit cognitive processes and more specifically, implicit statistical learning. These capacities might help mitigate the challenges of harsh and unpredictable environments (Young et al., 2020). Unpredictability might be especially influential in shaping implicit learning abilities as a quick and accurate estimation of environmental regularities can help to efficiently predict threats and opportunities, which would have clear fitness consequences (Young et al., 2018). Accordingly, the abovementioned studies on sensitivity to social influence (Jacquet et al., 2019), set shifting (Mittal et al., 2015; Young et al., 2022) and working memory updating (Young et al., 2018) tended to find effects of unpredictability, but weaker or non-existent effects of harshness. Increased implicit learning can be a useful complement to the enhancement of these explicit systems, due to its quicker, and attentionally and energetically less demanding nature (Conway, 2020; Hardwick et al., 2019; Janacsek & Nemeth, 2013; Schwabe & Wolf, 2013).

This idea dovetails research showing that while acute stress diminishes the flexible, hippocampal, declarative memory system, it seems to enhance the striatal memory system responsible for habitual, rigid behaviour (Dolfen et al., 2019; Schwabe & Wolf, 2012; Tóth-Fáber et al., 2021). Studies using reinforcement learning models have also investigated the effects of acute stress on model-based and model-free learning systems, generally thought to be the computational bases of goal-directed, declarative, and habitual procedural learning, respectively (Daw et al., 2005). These studies have similarly shown that acute stress diminishes the contribution of model-based learning to behaviour, while leaving model-free learning intact (Heller et al., 2018; Otto et al., 2013; Park et al., 2017). Finally, a recent study by Sherman et al. (2024) also revealed some evidence for an enhancement of implicit statistical learning by acute stress.

There have been far fewer studies that looked at the effects of chronic childhood stress on such implicit statistical learning (SL) abilities. Implicit SL refers to the set of neurocognitive processes that enable the extraction of environmental regularities in an incidental and unsupervised fashion (Conway, 2020; Kaufman et al., 2010; Saffran et al., 1996; Ullman, 2004). This implicit knowledge can then contribute to the acquisition, storage, and use of various motor, cognitive, linguistic and social skills (Conway, 2020; Kaufman et al., 2010; Ullman, 2004) and in predictive processing (Éltető et al., 2022; Janacsek & Nemeth, 2012). Implicit SL also seems to be related to the model-free learning and habitual decision making processes described in the reinforcement learning literature (Doll et al., 2015; Graybiel & Grafton, 2015; Sherman, Turk-Browne, et al., 2024). We are aware of only a handful of studies that investigated implicit SL abilities in relation to childhood adversity, and which revealed mixed findings. Most of these studies compared individuals based on socio-economic status (SES). While SES is no substitute for adversity (Amso & Lynn, 2017), it does correlate with exposure to adverse childhood experiences (Suglia et al., 2022; Walsh et al., 2019), and is related to nearly all forms of morbidity and mortality in Western societies (Adler & Newman, 2002; Ellis et al., 2009; Hughes et al., 2017). In a combined behavioural and neuroimaging study, Leonard et al. (2015) found weaker working memory performance and reduced hippocampus and dorsolateral prefrontal cortex volumes in individuals with lower SES. On the other hand, caudate volumes and probabilistic classification performance were not different between lower and higher SES individuals. Dang et al. (2016) similarly compared lower and higher SES individuals in a categorization task thought to rely on procedural learning mechanisms. They found that, contrary to previous reports, suggesting impaired explicit cognitive function in low-income individuals under financial stress (Mani et al., 2013), lower SES individuals outperformed higher SES individuals when financial concerns were introduced. They reasoned that it is precisely the reduction of explicit processing by such financial stress that allowed the more efficient procedural system to take the lead in guiding behaviour. Finally, Sheridan et al. (2018) compared three groups of children on two associative learning tasks, a monetary incentive delay task assessing reward learning and a serial reaction time task assessing implicit motor skill learning. One group of children were a control group of never institutionalised children, who were compared with two groups of children who either received prolonged institutional care or were placed into foster care. The prolonged institutional care group performed significantly worse than both other groups on both the reward learning and implicit motor skill learning tasks. The foster care group’s performance was closer to the control group on both tasks.

There are a number of factors that might have contributed to these mixed findings. Firstly, the reliability and validity of many implicit learning and procedural memory tasks are questionable, raising serious doubts about whether the tasks employed so far measure the same construct, and with enough sensitivity to study interindividual differences (West et al., 2018, 2019; but see Conway et al., 2019). Studies also tended to employ post learning, offline measures of performance, which do not allow us to study whether adversity affects the dynamics of learning (Siegelman et al., 2017). The implicitness of established knowledge was also rarely verified, and there was seldom a separation of current from early life adversity, leaving important potential confounds uncontrolled for (Frankenhuis, Young, et al., 2020). Finally, many studies reduce early adversity to the morbidity/mortality risks conveyed by economic scarcity, thereby neglecting the role of other dimensions of the environment that might affect development in a specific way, such as the stochastic variation in these risks inside and outside the family. These studies also frequently rely on group comparisons between ‘low’ and ‘high’ SES subgroups of relatively small sample sizes, resulting in low statistical power.

In the present research, we aimed to overcome these limitations, and offer a comprehensive investigation of the effects of both early life adversity and current subjective SES on multiple phases of implicit statistical learning in a large online sample. Our measurement of early life adversity focused on capturing interindividual variability in levels of exposure to harshness (i.e., mortality-morbidity risk, or cues to it) and unpredictability (i.e., stochastic spatiotemporal variability in harshness, or cues to it) in the ‘typical’ range during childhood (Ellis et al., 2009), which we defined as the period from infancy to 12 years of age. We employ a task proven to be both valid (Buffington et al., 2021) and reliable (Farkas et al., 2023), and robustly demonstrated to result in purely implicit knowledge (Vékony et al., 2021). In addition, we assessed associations between early life adversity and current executive functions and multiple dimensions of psychopathology, as a validity check of our adversity operationalisation. We expected that childhood unpredictability will be associated with increased SL ability. Regarding which phase of learning would be enhanced, we were agnostic due to the lack of previous empirical or theoretical work. Based on the results of previous studies (Doom et al., 2016; Jacquet et al., 2019; Mittal et al., 2015; Young et al., 2018), we expected weaker or non-existent effects of childhood harshness. Regarding the effect of current subjective SES, we were also agnostic. Our results suggest that while current subjective SES has a negative effect on implicit learning, moderate levels of early life unpredictability is associated with somewhat improved implicit learning.

## Methods

### Participants

A total of 406 university students took part in an online experiment. Participants received course credit for their participation. Informed consent was obtained from all participants, and the study was approved by the Research Ethics Committee of Eötvös Loránd University, Budapest, Hungary (Decision 2024/214).

From this initial sample, we excluded participants who: 1) at any point quit or restarted the experiment (N = 21, 5.17%); 2) had missing data on demographic information or any of our early life adversity variables (N = 14, 3.45%); 3) had self-reported neurological impairment or head injury, or a self-reported diagnosis of epilepsy, Autism spectrum disorder, Attention deficit hyperactivity disorder, Obsessive-compulsive disorder, Schizophrenia, or Post-traumatic stress disorder (N = 26, 6.40%); 4) reported having consumed alcohol or other recreational drugs or psychoactive medications within 6 hours of commencing the experiment (N = 36, 8.87%); 5) had a delay of more than 120 minutes between the two phases of the experiment (see below) (N = 2, 0.49%). These combined criteria led to the exclusion of a total of 81 participants (19.95%, note that participants could meet more than one exclusion criteria). The final sample thus consisted of the remaining 325 participants (239 (73.54%) females; M_age_ = 22.92 years ± 5.11 SD).

### Alternating Serial Reaction Time (ASRT) task

The ASRT task was programmed in JavaScript using the jsPsych library v.6.1.0. The task involves presenting participants with a visual stimulus (a drawing of a dog’s head) in one of four horizontal locations on the screen. Participants were instructed to indicate the location of the target stimulus by pressing the corresponding key on the keyboard (S, F, J, or L keys). In case of a correct response, the target stimulus disappeared, and after a 120 ms interstimulus interval, the next stimulus appeared. In case of an incorrect response, the target stimulus remained in place until the first correct response. Unbeknownst to the participants, the stimuli followed a probabilistic eight-element sequence, with pattern and random elements alternating with each other (Figure 1A, e.g., 1 – r – 2 – r – 4 – r – 3 – r, where r indicates a random location, and the numbers represent the predetermined positions from left to right). Each participant was assigned to one of 24 possible sequences.

**Figure 1.**
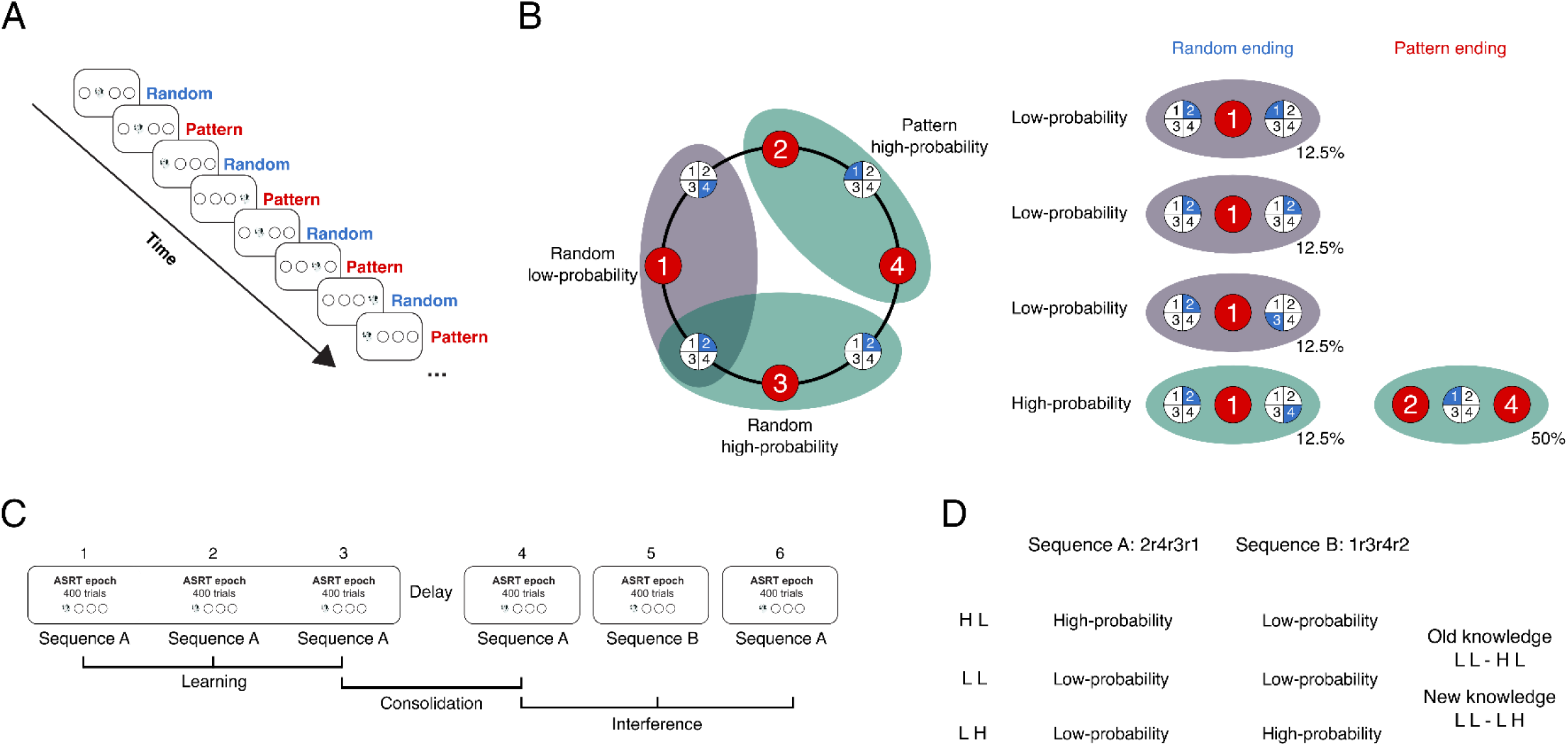
Study design and structure of the ASRT task. **A)** In the ASRT task, participants have to press keys corresponding to the location of the target stimulus (dog’s head). Every second trial forms the deterministic part of an 8-element sequence. Random elements are inserted between pattern elements to form the sequence (e.g., 2-r-4-r-3-1-r, where numbers indicate the location of pattern trials from left to right, and *r* represent random positions). **B)** Every trial can be categorized as the third element of three consecutive trials (a triplet). The probabilistic sequence structure results in some triplets occurring with a higher frequency (high-probability triplets, 62.5% of all trials) than others (low-probability triplets, 37.5% of all trials). Statistical learning is operationalised as the performance improvement in high-probability trials, compared to low-probability trials. **C)** Our design consisted of a learning phase, where participants were exposed to the same sequence (Sequence A) for a total of 1200 trials (for analyses divided into 3 Epochs of 400 trials each). This was followed by a self-paced delay of approximately 50 minutes. In the second phase, there was an initial Epoch with the originally practice sequence (Sequence A), followed by an Epoch with a new sequence (Sequence B), and finally an Epoch with the original sequence once again (Sequence A). Models assessing the *learning and acquisition of a probabilistic regularity* concern the first 3 Epochs, models assessing the *consolidation of a probabilistic regularity* concern the 3^rd^ and 4^th^ Epoch, and models assessing the *resistance to interference of a previously learned probabilistic regularity* concern Epochs 3 to 6. **D)** Due to the introduction of a novel sequence, the predictability of triplets changes during the task. As Sequence B is a reversed version of Sequence A, there are three categories of such changes: High-probability triplets in Sequence A become low-probability in Sequence B (‘H L’); Some low-probability triplets in Sequence A become high-probability in Sequence B (‘L H’); Some low-probability triplets in Sequence A remain low-probability in Sequence B (‘L L’). We can assess the resistance of *old implicit knowledge* by comparing performance between ‘L L’ and ‘H L’ triplets, with the expectation that performance should be better for ‘H L’ than ‘L L’ triplets even after exposure to the novel sequence during Epoch 5. We can also assess the acquisition of *new implicit knowledge* by comparing performance between ‘L L’ and ‘L H’ triplets, with the expectation that performance should be better for ‘L H’ than ‘L L’ triplets after exposure to the novel sequence during Epoch 5.

The underlying sequence gives rise to a probabilistic structure where certain runs of three consecutive stimuli (triplets) appear with a higher frequency (high-probability triplets) than others (low-probability triplets). A trial refers to a single element in the sequence that could be either a pattern or random element, and, crucially, also the last element in a high- or low-probability triplet. For example, in a sequence such as 1 – r – 2 – r – 4 – r – 3 – r, triplets such as 1-X-2, 2-X-4, 4-X-3, and 3-X-1 (where X represents the middle element of a triplet) occur more frequently than triplets such as 2-X-1 or 2-X- 3 (Figure 1B). Each trial can thus be categorised as high- or low-probability based on the previous two trials. Specifically, high-probability triplets occur in 62.5% of all trials, and low-probability triplets occur in the remaining 37.5% of all trials. An extensive literature has robustly demonstrated that participants show performance facilitation in both reaction times and accuracy to high-probability triplets, over low-probability triplets, indicating a sensitivity to the underlying probabilistic regularity of their environment (J. H. Howard et al., 2008; Kóbor et al., 2017; Nemeth, Janacsek, Balogh, et al., 2010; Song et al., 2007; Takács et al., 2018; Tóth-Fáber et al., 2023). Moreover, this knowledge is entirely implicit, with no evidence of the contribution of explicit processing as assessed by both self-reports and sequence generation procedures (Vékony et al., 2021). We also confirmed this by asking participants a series of questions at the end of the ASRT task. Specifically, they were asked if they noticed anything unusual or any regularities in the task, and if so, to elaborate on their response. Most participants indicated they did not notice any regularity (N = 201 (61.85%)). Among those who indicated some perceived regularity (N = 124 (38.15%)), 17 (13.71%) felt uncertain about its accuracy, 73 (58.9%) felt neither certain nor uncertain, and 34 (27.4%) felt certain. When asked to elaborate on the perceived regularity, none of the participants were able to describe the sequence accurately (example responses include “Maybe the ‘patterns’ repeated?”; “Always began on the right, the 3rd position was the least frequent”; “It consisted of three 5-element sequences”; “If I was incorrect in a response, it was repeated after some time”). Thus, the performance differences between triplets are indices of *implicit SL*. In addition to this, there tends to be an overall *visuomotor performance* improvement in RT and accuracy during the task, irrespective of triplet probability (J. H. Howard et al., 2008; Nemeth, Janacsek, Balogh, et al., 2010; Tóth-Fáber et al., 2023). The separation of these two processes is helpful to control for possible confounds such as recall or practice effects, and is one of the main strengths of the task (Nemeth, Janacsek, Londe, et al., 2010; Song et al., 2007).

The task was divided into blocks of 80 trials, with short self-paced rest periods between blocks. The first two blocks were practice blocks, with no underlying sequence. These two blocks were not analysed. To facilitate data analysis, we aggregated the remaining blocks into units of 5 blocks, henceforth referred to as epochs. The experiment consisted of 6 such epochs in total. The experiment was divided into a learning phase, a consolidation phase, and an interference phase, with the learning and consolidation phases separated by self-paced delay period that lasted approximately 50 minutes (M = 43.19 minutes ± 15.88 SD, Supplementary Figure S1). In the learning phase, participants completed 3 epochs of the task (15 blocks, 1200 trials) with a predetermined sequence (Sequence A, Figure 1C). This phase allows us to measure the *learning and acquisition* of a probabilistic regularity (by comparing high- and low-probability triplets), and the overall effect of practice with the task in visuomotor performance (by comparing overall performance between successive epochs). After the delay period, the consolidation phase followed. This phase consisted of participants being presented with the initially learned sequence again for one epoch. This was followed by an interference phase when in epoch 5, without warning, the underlying sequence was changed to the reversed version of the original sequence (Figure 1D, e.g., if Sequence A was 1 – r – 2 – r – 4 – r – 3 – r, then Sequence B was 3 – r – 4 – r – 2 – r – 1 – r). This change leads to certain triplets changing their status between low-probability and high-probability between sequences. Specifically, three different categories of triplets can be differentiated during the interference phase (Figure 1D). ‘H L’ triplets were high-probability in Sequence A, but low-probability in Sequence B. ‘L H’ triplets were low-probability in Sequence A, but high-probability in Sequence B. ‘L L’ triplets were low-probability in both Sequences. As the sequences were reversed versions of each other, they were entirely non-overlapping, making ‘H H’ triplets not possible. Finally, during epoch 6, participants were once again presented with Sequence A. This design allows us to measure three additional aspects of implicit SL (Figure 1C). First, by comparing performance between the final epoch of the learning phase (Epoch 3) and the first epoch of the interference phase with the original sequence (Epoch 4), we can measure the effect of *consolidation* during the delay period, both in terms of overall performance improvements in visuomotor performance, and the implicit SL of the probabilistic regularity (Szegedi-Hallgató et al., 2017). Second, by comparing performance between ‘H L’ and ‘L L’ triplets during the interference epochs, we can measure the extent to which previously established implicit knowledge and practice is *resistant to interference* by novel environmental regularities (Horváth et al., 2022). Third, by comparing performance between ‘L H’ and ‘L L’ triplets during the interference epochs, we can measure the extent of which a *novel environmental regularity can be learned* that conflicts with previously established implicit knowledge (Figure 1D) (Horváth et al., 2022).

### Adversity dimensions

Our measurement of early life adversity focused on capturing each participant’s exposure to harshness (i.e., mortality-morbidity risk, or cues to it) and unpredictability (i.e., stochastic spatiotemporal variability in harshness, or cues to it) during their childhood. These two dimensions of early adversity were proposed by Ellis et al. (2009) to be the most influential in guiding development and give rise to adaptive responses, that can include the modulation of both reproductive and health behaviours, personality traits, and neurocognitive processes (Belsky et al., 2012; Ellis et al., 2009; Farkas et al., 2022; Szepsenwol et al., 2017). Our operationalisation was guided by the assumptions of the dimensional models of adversity, which propose that it is the distinct developmental consequences of adversity factors, rather than their co-occurrence patterns that should define relevant dimensions, i.e., certain experiences do not belong to the same dimension because they co-occur, but rather because their effect on development are similar in their mechanisms, and they can serve as reliable cues of the same sources of environmental adversity (Berman et al., 2022; Ellis, Sheridan, et al., 2022; McLaughlin et al., 2019, 2021). For example, unstable parental work status and material deprivation are likely highly correlated, yet we believe the former to be an experience signaling environmental unpredictability, while the latter signaling environmental harshness, each expected to trigger different types of responses from the developing organism. As such, researchers generally recommend the use of composites of possibly only weakly correlated adversity indicators based on *a priori* theoretical and empirical demonstration of their effects on psychological and biological development, instead of reflective latent variable approaches based on mere co-occurrence patterns (Berman et al., 2022; Brumbach et al., 2009; McLaughlin et al., 2021, 2023; Mell et al., 2018). In our assessment of harshness, we include items related to both subjective and objective socio-economic status. While adversity and SES likely impact child development through different mechanisms (Amso & Lynn, 2017), the use of SES as a proxy for harshness is justified by its association with nearly all forms of morbidity and mortality in Western societies (Adler & Newman, 2002; Ellis et al., 2009; Hughes et al., 2017), including in Hungary (Csilla et al., 2010; Skultéti & Pikó, 2006; Szőcs et al., 2019; Tombor et al., 2010). In our assessment of unpredictability, we focused on experiences signaling unpredictable environments in different contexts on multiple timescales. A weakness of this approach is that it does not separate contexts (Munakata et al., 2023), timescales (Farkas et al., 2024), or unpredictability formalisations (Farkas & Jacquet, 2024; Walasek et al., 2024), which remain for future work to disentangle. We mainly rely on subjective assessment of adversity by participants, instead of objective indicators, due to strong evidence that perceptions of adverse events are often much more important determinants of psychological or physiological responses, than objective events themselves (Bollini et al., 2004; Francis et al., 2023; Martinez et al., 2022; Rivenbark et al., 2020) and issues with ‘objective’ definitions of adversity in terms of ‘expectable environments’ (Frankenhuis & Amir, 2022).

### Unpredictability

We measured childhood unpredictability with a modified version of the Questionnaire of Unpredictability in Childhood (QUIC) - a self-report measure designed to assess lack of predictability in multiple aspects of the childhood environment (social, emotional, and physical) (Glynn et al., 2019). Items are binary, with ‘yes’ (= 1) or ‘no’ (= 0) response options, with some items reverse coded. Example items include ‘At least one of my parents was unpredictable’ and ‘At least one of my parents regularly checked that I did my homework’. QUIC total sum scores have demonstrated strong internal consistency, excellent test-retest reliability and have been shown to predict other unpredictability indicators and internalising symptoms, suggesting good construct validity (Glynn et al., 2019). We made three important changes to the original questionnaire. First, the original questionnaire uses a timeframe of ‘prior to age 12’ for some items and ‘prior to age 18’ for others. We modified this, such that in our implementation all questions were phrased so as to refer to the period ‘prior to age 12’. Second, as we felt that the Safety and Security subscale measures harshness, instead of unpredictability, we included the 3 items of that subscale in our harshness composite instead (note that this was done before any data collection has taken place). Finally, we were sceptical about the validity of a question about family holiday traditions as an unpredictability indicator and decided to not administer it. Total sum scores for our final unpredictability measure thus ranged from 0 to 34, with a higher score indicating greater exposure to unpredictability before the age of 12. Observed scores had a median of 5, with a median absolute deviation of 4.45, and range = [0,27] (Supplementary Figure S1). This measure was z-scored before being entered into the models.

### Harshness

We measured childhood harshness with a set of custom items, items adapted from Mittal et al. (2015), and the abovementioned Safety and Security subscale of the QUIC. All of the items were binary, with ‘yes’ (= 1) or ‘no’ (= 0) response options, with some items reverse coded. Specifically, our measurement consisted of 3 items adapted from Mittal et al. (2015) assessing relative socio-economic status (‘When I was younger than 12 … My family usually had enough money for things when I was growing up.’; ‘When I was younger than 12 … I grew up in a relatively wealthy neighborhood.’; When I was younger than 12 … I felt relatively wealthy compared to other kids in my school.), the 3 items of the Safety and Security subscale of the QUIC assessing intrafamilial deprivation and threat (‘When I was younger than 12 … There was a period of time when I often worried that I was not going to have enough food to eat.’; ‘When I was younger than 12 … There was a period of time where I did not feel safe in my house.’; When I was younger than 12 … I have suffered from sexual or physical abuse.’) and 6 custom items asking whether the participant experienced the death of: their father; their mother; a sibling; another member of their close family; another family member or friend; or another person who was part of their daily environment, before the age of 12. What ties these experiences together is that they are indicative of an environment relatively lower in resources and security, capturing two subtypes of harshness defined by recent models, deprivation and threat, respectively (Ellis, Sheridan, et al., 2022). The Safety and Security subscale items (more absolute SES) and the Mittal et al. (2015) items (more relative SES) were moderately correlated (r = .269, 95% CI = [.165, .367], p < .001). Total sum scores for our final harshness measure ranged from 0 to 12, with a higher score indicating greater exposure to unpredictability before the age of 12. Observed scores had a median of 2, with a median absolute deviation of 1.48, and range = [0,8] (Supplementary Figure S1). This measure was z-scored before being entered into the models.

### Current subjective SES

To assess subjective current socio-economic status (cSES), participants completed a translated version of the MacArthur Scale of Subjective Social Status (Adler et al., 2000), a single-item measure of an individual’s perceiver rank relative to others in their group, with 10 response options ranging from 1 (worst off) to 10 (best off). Thus, higher values indicate lower perceived subjective social status. The median response was 5, with a median absolute deviation of 1.48, and range = [2,9] (Supplementary Figure S1).

### Parental education

To measure a more objective component of early life deprivation, we asked participants to report the highest educational status obtained by both of their parents. Participants could choose between the following options, that represented the typical ordinal classification within the Hungarian education system: 1 = “Legfeljebb 8 általános” (at most 8^th^ grade in primary school, <= 8 years of education); 2 = “Szakiskola vagy szakmunkásképző” (Vocational school, 12 years of education); 3 = “Szakközépiskola” (Technical school, 12 years of education); 4 = “Gimnázium” (Grammar school, 12 years of education); 5 = “Főiskolai vagy egyetemi alapfokozat” (Bachelor’s degree, 15 years of education); 6 = “Egyetemi mesterfokozat” (Master’s degree, 17 years of education); 7 = “Egyetemi posztgraduális fokozat” (Postgraduate degree, more than 17 years of education) (European Commission: European Education and Culture Executive Agency, 2023).

Maternal education had the following distribution: 9 (2.8%) at most 8^th^ grade in primary school; 31 (9.5%) Vocational school; 55 (16.9%) Technical school; 47 (14.5%) Grammar school; 115 (35.4%) Bachelor’s degree; 61 (18.8%) Master’s degree; 7 (2.2%) Postgraduate degree.

Paternal education had the following distribution: 7 (2.1%) at most 8^th^ grade in primary school; 60 (18.5%) Vocational school; 60 (18.5%) Technical school; 45 (13.8%) Grammar school; 81 (24.9%) Bachelor’s degree; 56 (17.2%) Master’s degree; 16 (4.9%) Postgraduate degree.

A composite Parental education score was created by averaging the numerical values corresponding to the options across the two parents. This composite was treated as a continuous variable (median = 4, median absolute deviation = 1.48, range = [1,7]) (Supplementary Figure S1).

### Additional measures

We collected additional data regarding participants’ executive functions and mental health, with the aim of using this data to validate our early life adversity operationalisation. Based on extensive previous research, we expected that both childhood harshness, childhood unpredictability and subjective cSES would be moderately positively associated with overall mental health and moderately negatively with executive functions. We assessed these relationships with bivariate Pearson’s linear correlations, uncorrected for multiple comparisons, both because they each tested distinct hypotheses and thus do not form a single family of tests, and because we were more interested in reporting the direction and magnitude of associations, rather than statistical inference about their difference from zero.

### Executive functions

Detailed description of executive functions measures can be found in Supplementary Text S1. We measured phonological short-term memory capacity using a computerised version of the digit span task (Isaacs & Vargha-Khadem, 1989); complex working memory with the 1-BACK task (Kirchner, 1958); cognitive inhibition with the Go / No-go task (Bezdjian et al., 2009); and cognitive flexibility with the Berg Card Sorting Task (Berg, 1948; Fox et al., 2013).

### Psychopathology

Detailed description of mental health measures can be found in Supplementary Text S1. Our mental health questionnaires included the Adult ADHD Self-Report Scale (ASRS) (Kessler et al., 2005); the Obsessive-Compulsive Inventory – Revised (OCI-R) (Foa et al., 2002); the Autism-Spectrum Quotient (AQ) (Baron-Cohen et al., 2001); the Eating Attitude Test (EAT) (Garner et al., 1982); the Hypomania checklist (HCL) (Angst et al., 2005); and the Brief Multidimensional Schizotypy Scale (MSS-B) Gross et al. (2018).

### Preprocessing and statistical analysis

Each trial of the ASRT task was categorized based on the two preceding trials into the different triplet categories, as being the last element of either a high- or a low-probability triplet (Figure 1B). Trills (e.g., 1-2-1) and repetitions (e.g., 2-2-2) were removed, as participants can show pre-existing response tendencies, such as automatic facilitation, to these types of trials (D. V. Howard et al., 2004; Soetens et al., 2004). The first two trials of each block were also removed, as no triplet membership can be identified for them. Inaccurate responses were additionally removed from the analyses of RTs. These criteria led to the removal of 31.8% of trials from epochs 1, 2, and 3 for the learning analyses of RTs (24.7% for the accuracy models); 22.5% of trials from epochs 3 and 4 for the consolidation analyses of RTs (14.7% for the accuracy models); and 22.2% of trials from epochs 4, 5, and 6 for the interference analyses of RTs (14.7% for the accuracy models). These may seem high, but they are standard for analyses of the task, and most exclusions come from trills and repetitions (see e.g., D. V. Howard et al., 2004; Pesthy et al., 2023; Soetens et al., 2004; Song et al., 2007, 2009).

In reporting our generalised linear mixed model (GLMM) analyses, we follow best practice guidelines proposed by Meteyard and Davies (2020). All pre-processing and statistical analysis was performed in *R* 4.2.1. GLMMs were fit with the *mixed* function from the *afex* package (Singmann et al., 2023). No a priori power analysis was done for the study. Our primary independent variables of interest were Epoch, Triplet Type, Harshness, Unpredictability and subjective cSES. Triplet Type was a factor variable, effects coded to reflect the high- and low-probability triplet categories, with High-probability triplets being the reference category. Epoch was a factor variable, coded to reflect Epochs (1 to 6), with the last relevant Epoch being the reference category in all models. Harshness and Unpredictability were continuous variables, which were z-scored. Covariates of Age, Delay duration and subjective cSES were continuous and were mean-centered. Our models contained fixed effects for the main effect of Triplet type, Epoch, subjective cSES, Harshness and Unpredictability, Age and Delay duration (for the consolidation models only) and the higher order interactions of Triplet type and Epoch with the other variables. The models further specified participant-specific intercepts and correlated slopes of the within-subject variable of Epoch, as random effects. Thus, our models corresponded to the following general form in *lmer* syntax:

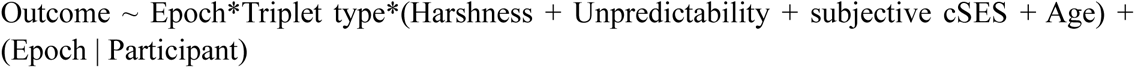

Delay duration was included as a control variable, as it is likely to be associated with performance, but unlikely to be associated with our adversity variables. We confirmed that this is indeed true, as it correlates with both overall RT (r = .184, 95% CI = [.076, .287], p < .001) and overall accuracy (r = .101, 95% CI = [-.008, .208], p = .069) in Epoch 4, but not with any of the adversity variables (with Harshness: r = -.035, 95% CI = [-.143, .074], p = .529; with Unpredictability: r = -.020, 95% CI = [-.128, .089], p = .726; with Current SES: r = .048, 95% CI = [-.061, .156], p = .387). This means that it shares variance with the outcome, but not our predictor variables, so including it in our models will lower residual variance in the outcome and lead to greater statistical power (Carlson & Wu, 2012; Wysocki et al., 2022). Age was found to not only share variance with our outcome variables (with overall RT: r = .175, 95% CI = [.067, .278], p = .002; with overall accuracy: r = .092, 95% CI = [-.017, .199], p = .098), but with adversity as well (with Harshness: r = .152, 95% CI = [.044, .256], p = .006; with Unpredictability: r = .089, 95% CI = [-.020, .196], p = .109; with subjective cSES: r = -.027, 95% CI = [-.136, .082], p = .624), meaning that its inclusion as a statistical control is also warranted as it might be a confound (Carlson & Wu, 2012; Wysocki et al., 2022). Assumptions regarding linearity, homoscedasticity and normality of residuals were evaluated by scatterplots and QQ plots of residuals. Assumptions of linearity and homoscedasticity were met in each case, but some of the models presented with non-normal residuals. We note however, that fixed effects estimates of LMMs have been shown to be remarkably robust against violation of this assumption (Schielzeth et al., 2020). Therefore, we deemed our models adequate.

We report sample sizes in terms of total number of data points and of sampling units, random effects estimates, Nakagawa’s marginal and conditional R^2^ (Nakagawa & Schielzeth, 2013), and the adjusted ICC in summary tables. Post-hoc contrasts were conducted with the *emmeans R* package, after regridding (Lenth, 2022). Significant interaction effects with continuous variables were interpreted by estimating performance at low (mean-1SD) and high (mean+1SD) levels of the relevant continuous variable. For inference about fixed effects, we used Type III tests (for RT models) or Likelihood ratio tests (for accuracy models), relying on comparing nested models with the effect of interest either included or removed. For RT models, degrees of freedom were obtained using the Satterthwaite approximation (Satterthwaite, 1941), for the accuracy models, they are asymptotic, making the test statistic a z statistic, instead of t statistic (Singmann & Kellen, 2019). Figures were created with *ggplot2* (Wickham, 2016). An alpha level of .05 was applied to all analyses. P values of all post hoc tests were adjusted with the Šidák method in case of multiple comparisons across Epochs (Šidák, 1967). All p values are two-tailed. The study was not pre-registered. Effect sizes are interpreted according to criteria laid out by Funder & Ozer (2019) and Gignac & Szodorai (2016) for interindividual differences psychology research, with r = .10, r = .20, and r = .30, corresponding with small, medium and large effects, respectively. All data and code necessary to reproduce the results of this study are available on the OSF platform, at the following link (NOTE: This link is currently anonymized for peer review): https://osf.io/gmr35/?view_only=c4da4956159f46128007e9c6cef3fba9. The code for the ASRT task can also be found at the following GitHub Repository: Link currently not included to maintain anonymity.

## Results

### Do our measures of early life and current adversity predict psychopathology and executive functions?

Before performing our analysis of the ASRT data, we wished to establish the validity of our adversity measures. To do this, we first inspected the distributions and summary statistics of the adversity variables (Supplementary Figure S1). This indicated that subjective cSES and Parental education are normally distributed around the respective mean of both scales. Childhood harshness and unpredictability are positively skewed, with very few individuals reporting having experienced high early life adversity, but with decent variability in the low and middle range of the spectra. This follows the pattern expected based on previous estimates of adverse childhood experience prevalence rates (Karatekin & Hill, 2019; Madigan et al., 2023). Overall, this suggests that we are well positioned to investigate variability of adverse childhood experiences in the low to middle range, but are likely unable to say much about the highest end of the spectrum. Then, we looked at the pattern of bivariate correlations between our childhood Harshness, childhood Unpredictability, Parental education, and subjective cSES measures, and multiple executive functions (EF) and subclinical psychopathology measures. Our EF measures included the Digit Span task, the 1-Back task, the Go / No-go task, and the Berg Card Sorting Task. Our psychopathology questionnaires included the Adult ADHD Self-Report Scale, the Obsessive-Compulsive Inventory – Revised, the Autism-Spectrum Quotient questionnaire, the Eating Attitude Test, the Hypomania checklist, and the Brief Multidimensional Schizotypy Scale (for more detail see Supplementary Text S1). Based on the literature, we expected that early life adversity should be associated with increased psychopathology and decreased EF performance. Our prediction was supported for psychopathology, as both Harshness and Unpredictability were modestly and positively correlated with OCD (r = .15, r = .21), ADHD (r = .15, r = .25), Eating disorder (r = .14, r = .24) and Schizotypy (r = .17, r = .32) traits and Unpredictability was additionally correlated with the Autism Quotient (r = .18) (Supplementary Figure S2). However, neither was correlated with any of the EF components tested (all correlations between r = -.10 and r = .05). Subjective cSES was not related to either psychopathology or EF (all correlations between r = -.07 and r = .11). This pattern of results suggests that in our sample, early life harshness and unpredictability tended to predict later life subclinical symptoms of multiple psychopathologies, supporting the validity of our operationalisation, but they did not seem to be associated with impaired EF abilities, potentially due to the relative lack of individuals from highly adverse backgrounds.

### Do early life and current adversity impact the initial implicit acquisition of environmental regularities?

We first evaluated implicit SL and visuomotor performance, and their relationship with childhood harshness and unpredictability and subjective cSES in the learning phase of the experiment (Figure 2). Trial-level RT (log transformed) and accuracy (0: incorrect vs 1: correct) were used as the outcome variables in GLMMs, containing fixed effects of Epoch (Factor: 1, 2, 3), Triplet type (Factor: High, Low), subjective cSES, Harshness, Unpredictability and Age, and their higher order interactions, with participant-specific correlated intercepts and slopes for the within-subjects factor of Epoch. Throughout the presentation of the results, main effects of Epoch are interpreted as indicating visuomotor performance, independent of statistical regularities, while main effects and interactions of Triplet type are interpreted as indicating implicit SL and its trajectory during the task. Henceforth, the term learning score always denotes performance differences between High and Low triplets, obtained from contrasts, with positive sign always corresponding with better learning. Statistical tests of the trajectories of visuomotor performance and implicit SL are omitted from the main text for brevity and are instead detailed in Supplementary Text S2. Figures always plot raw data, whereas post hoc tests are done using model estimated marginal means (Lenth, 2022). Note that adversity measures are incorporated into the model as continuous variables, and Low and High adversity subgroups are created only for post hoc testing and visualisation. To aid readability, we only detail the fixed effects corresponding to our effects of main interest here, the full list of fixed and random effects parameters can be found in Supplementary Table S1 and S2, for RT and accuracy, respectively.

**Figure 2.**
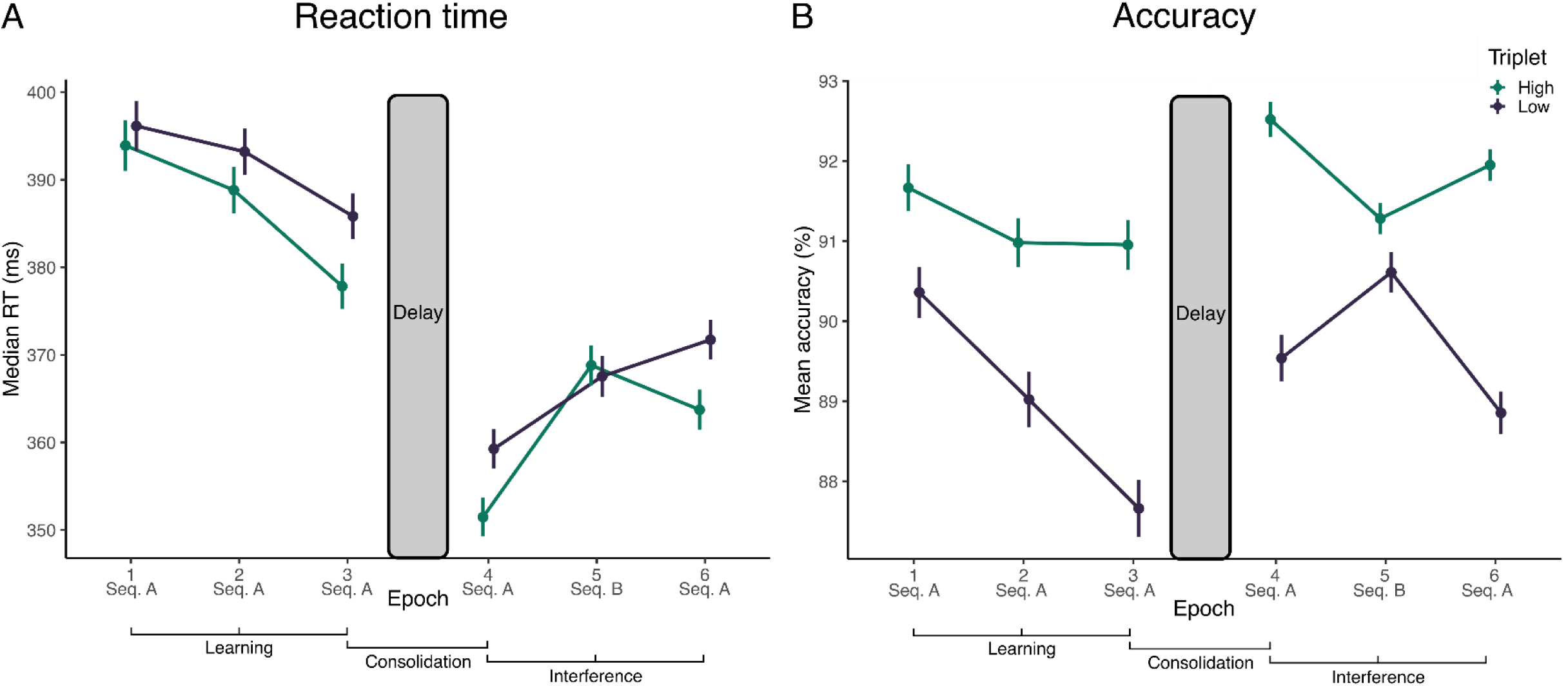
Trajectories of visuomotor performance and implicit statistical learning during the whole experiment. **A)** Performance in terms of epoch-wise median RT. X-axis indicates Epochs, y-axis indicates median RTs (in ms). Participants became overall faster during learning and showed further improvement following the delay. The introduction of the novel sequence during Epoch 5 disturbed performance. Participants also showed evidence of implicit SL, as they increasingly became faster for high-probability triplets, than for low-probability triplets during learning. This implicit knowledge was also perturbed by the introduction of the novel sequence. **B)** Performance in terms of epoch-wise mean accuracy. X-axis indicates Epochs, y-axis indicates median accuracies (in %). Participants became overall somewhat less accurate during learning but showed an improvement following the delay. The introduction of the novel sequence during Epoch 5 disturbed performance. Participants also showed evidence of implicit SL, as they increasingly became more accurate for high-probability triplets, than for low-probability triplets during learning. This implicit knowledge was also perturbed by the introduction of the novel sequence (note that Low triplets after the delay in this figure correspond to the sum of ‘H L’ and ‘L L’ triplets, whereas High triplets after the delay correspond to ‘H L’ triplets; compare with Figure 4). In all subfigures, points indicate means, error bars indicate 1 SEM.

In the RT model, main effects of Epoch (F_(2,337.39)_ = 55.45, p < .001), Triplet type (F_(1,289785.18)_ = 295.40, p < .001) and the Epoch x Triplet type interaction (F_(2,289788.17)_ = 20.17, p < .001) were all statistically significant, indicating both increasing visuomotor learning and implicit SL with time at the group level (Supplementary Text S2, Table S1 and Figure 2A). The model also resulted in a statistically significant Epoch x subjective cSES interaction (F_(2,337.02)_ = 3.16, p = .044). This stemmed from a somewhat different trajectory of overall performance in individuals with high versus low subjective cSES. Low cSES individuals showed a progressive decrease in RTs (Epoch 1: 392 ms, 95% CI = [384, 400]; Epoch 2: 388 ms, 95% CI = [381, 396]; Epoch 3: 383 ms, 95% CI = [375, 390]; Epoch 1-2 contrast p = .056, Epoch 1-3 contrast p < .001, Epoch 2-3 contrast p < .001), whereas high cSES individuals showed a similar decrease only during the final epoch (Epoch 1: 391 ms, 95% CI = [381, 401]; Epoch 2: 390 ms, 95% CI = [380, 400]; Epoch 3: 378 ms, 95% CI = [369, 387]; Epoch 1-2 contrast p = .980, Epoch 1-3 contrast p < .001, Epoch 2-3 contrast p < .001).

There was a statistically significant Epoch x Triplet type x Unpredictability interaction. This stemmed from a different trajectory of implicit SL in individuals exposed to high versus low unpredictability (F_(2,289784.06)_ = 3.61, p = .027) (Figure 3). Low unpredictability individuals showed an increase in SL between epochs 2 and 3 (learning scores in Epoch 1: 2.84 ms, 95% CI = [1.30, 4.39]; Epoch 2: 3.97 ms, 95% CI = [2.43, 5.52]; Epoch 3: 8.35 ms, 95% CI = [6.82, 9.88]; Epoch 1-2 contrast p = .671, Epoch 1-3 contrast p < .001, Epoch 2-3 contrast p < .001), whereas high Unpredictability individuals exhibited asymptotic SL performance already during the second epoch and did not improve further (learning scores in Epoch 1: 2.92 ms, 95% CI = [1.39, 4.44]; Epoch 2: 6.80 ms, 95% CI = [5.26, 8.34]; Epoch 3: 6.82 ms, 95% CI = [5.32, 8.32]; Epoch 1-2 contrast p = .001, Epoch 1-3 contrast p < .001, Epoch 2-3 contrast p > .999).

**Figure 3.**
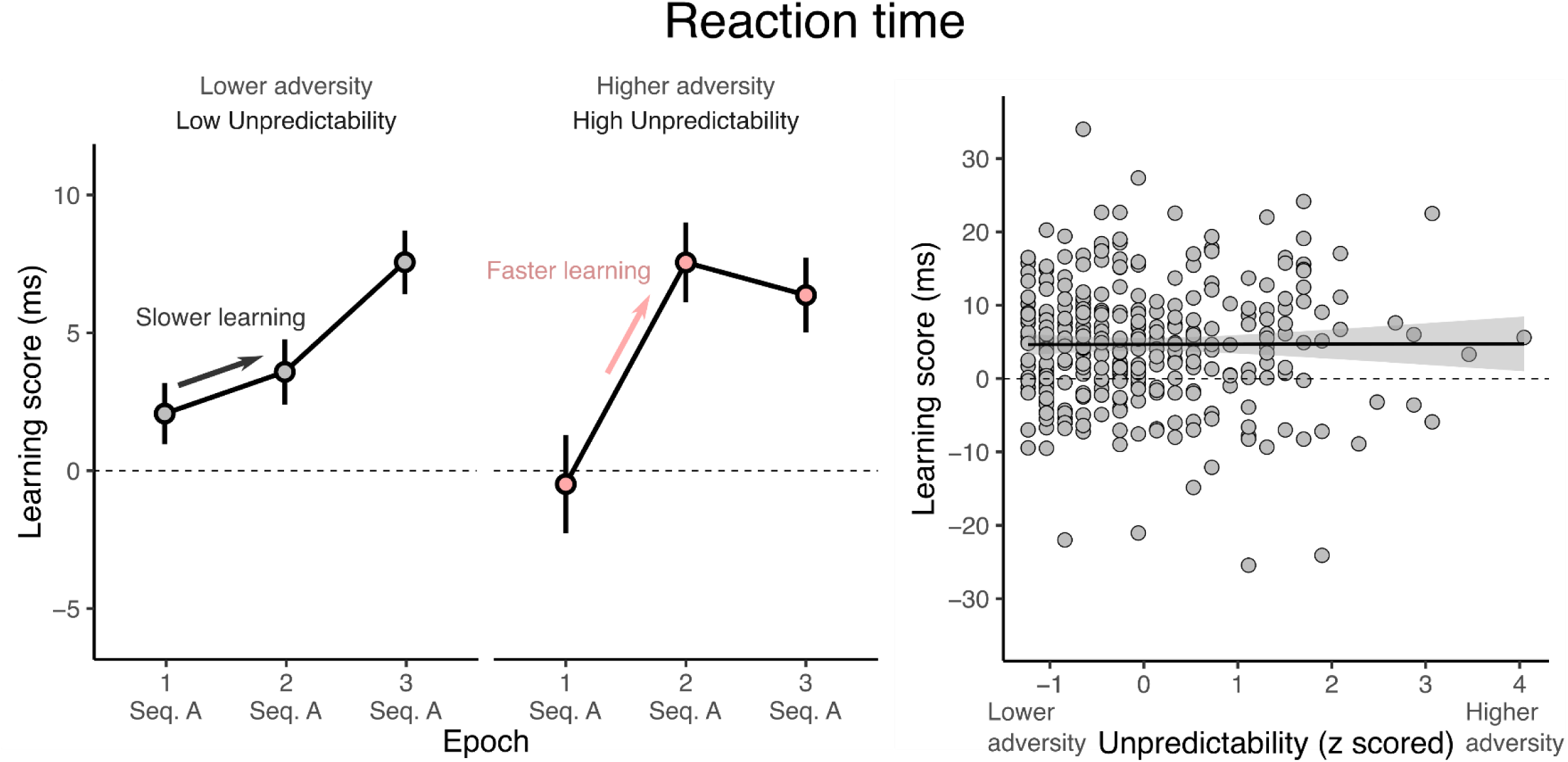
Effect of childhood unpredictability on initial learning. Epoch-wise (left) and whole task (right) average learning scores (RT difference between Low and High probability triplets) as a function of childhood unpredictability. Trajectories are separately shown for Low (lower third) and High Unpredictability (higher third) subgroups. Individuals with higher unpredictability achieved peak learning performance earlier but reached the same level. Note that splitting is only done for visualisation purposes and statistical models contain Unpredictability as a continuous variable. On the left figure, points indicate means, error bars indicate 1 SEM. On the right figure, points are subject specific means, line is line of best linear fit, with shading indicating 95% CI. Subfigures that correspond to statistically significant effects in the LMM are highlighted by pink.

No main effect or interaction involving Harshness was statistically significant. Overall, results suggest a facilitatory effect of current adversity on visuomotor performance, and of early life unpredictability on implicit SL, as assessed by RT.

In the accuracy model, main effects of Epoch (*χ^2^* (2) = 73.92, p < .001) and Triplet type (*χ^2^* (1) = 337.38, p < .001), as well as the Epoch x Triplet type interaction (*χ^2^* (2) = 38.10, p < .001) were statistically significant, and indicated increasing implicit SL, and a slight decrease in accuracy with time at the group level (Supplementary Text S2, Table S2 and Figure 2B). No main effect or interaction involving subjective cSES, Harshness or Unpredictability was statistically significant, suggesting that there is no support for early life or current adversity associations with initial learning, as assessed by accuracy.

### Do early life and current adversity impact the consolidation of extracted implicit probabilistic knowledge?

We proceeded to evaluate implicit SL and visuomotor performance, and their relationship with childhood harshness and unpredictability and subjective cSES during consolidation (Figure 2). After Epoch 3, participants took a self-paced break lasting on average 50 minutes. Thus, we evaluated the consolidation of visuomotor performance and implicit SL by comparing performance between Epochs 3 and 4. Trial-level RT (log transformed) and accuracy (0: incorrect vs 1: correct) were used as outcome variables in GLMMs, containing fixed effects of Epoch (Factor: 3, 4), Triplet type (Factor: High, Low), subjective cSES, Harshness, Unpredictability, Delay duration and Age, and their higher order interactions, with participant-specific correlated intercepts and slopes for the within-subjects factor of Epoch. Full list of fixed and random effects parameters can be found in Supplementary Tables S3 and S4, for RT and accuracy, respectively.

In the RT model, there was a statistically significant main effect of Epoch (F_(1,331.46)_ = 499.31, p < .001) and Triplet type (F_(1,193144.75)_ = 472.63, p < .001), but no significant effect of the Epoch x Triplet type interaction, corresponding with an increase in RT after the delay period, and the presence of implicit SL both before and after the delay, but no SL improvement after the break at the group level (Supplementary Text S2, Table S3 and Figure 2A). No main effect or interaction involving subjective cSES, Harshness or Unpredictability was statistically significant, providing no support for either early life or current adversity effects on the consolidation of visuomotor performance or implicit SL, as assessed by RT.

In the accuracy model, there was a statistically significant main effect of Epoch (*χ^2^* (1) = 32.33, p < .001) and Triplet type (*χ^2^*(1) = 493.07, p < .001), but no significant effect of the Epoch x Triplet type interaction, corresponding with an decrease in accuracy after the delay period, and the presence of implicit SL both before and after the delay, but no SL improvement after the break at the group level (Supplementary Text S2, Table S4 and Figure 2B). No main effect or interaction involving subjective cSES, Harshness or Unpredictability was statistically significant, providing no support for early life or current adversity effects on the consolidation of visuomotor performance or implicit SL, as assessed by accuracy.

### Do early life and current adversity impact the resistance of implicit probabilistic knowledge to interference?

Finally, we evaluated implicit SL and visuomotor performance, and their relationship with childhood harshness and unpredictability and subjective cSES during interference (Figure 4). After epoch 4, participants were presented with an epoch that, unknowingly to them, had a different underlying sequence structure. Specifically, the sequence was a reversed version of the sequence they had previously been exposed to. After this epoch, the final epoch of the task was governed by the original sequence again. For the analysis of this phase, we split triplets based on their status in the two sequences: ‘H L’ triplets were high-probability in the original sequence, but low-probability in the new sequence, ‘L H’ triplets were low-probability in the original sequence, but high-probability in the new sequence, and ‘L L’ triplets were low-probability in both (Figure 1D). This design allows us to investigate both the resistance of previously established implicit knowledge to interference (by comparing performance between ‘L L’ and ‘H L’ triplets), and the acquisition of new implicit knowledge (by comparing performance between ‘L L’ and ‘L H’ triplets).

**Figure 4.**
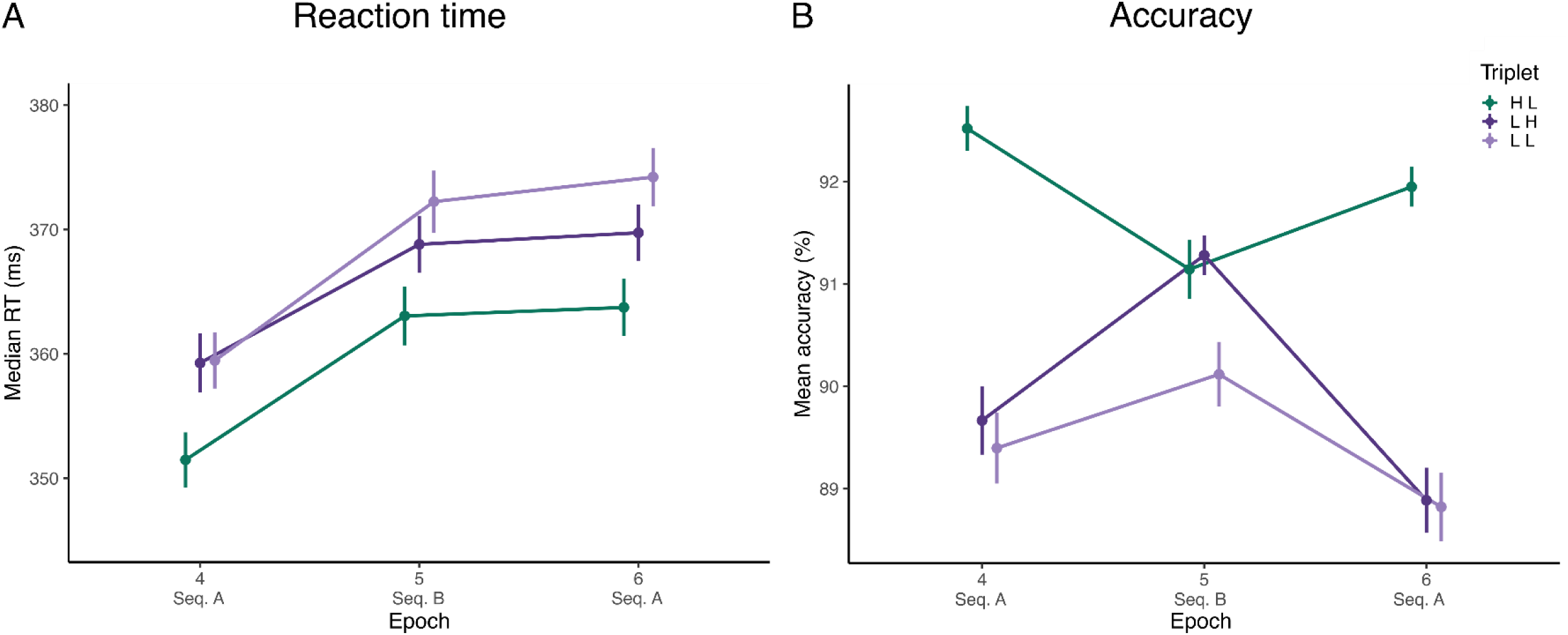
Trajectories of visuomotor performance and implicit statistical learning during interference. **A)** Performance in terms of epoch-wise median RT. X-axis indicates Epochs, y-axis indicates median RTs (in ms). Triplets are now separated according to how their predictability changed between Sequence A and B. Participants showed a successful acquisition of new implicit knowledge as they started to differentiate between ‘L H’ and ‘L L’ triplets during exposure to Sequence B during Epoch 5 and retained this performance difference even during Epoch 6, where the original sequence was reintroduced. Participants also showed a resistance of old implicit knowledge to interference as they retained the performance difference between ‘H L’ and ‘L L’ triplets even after exposure to Sequence B. **B)** Performance in terms of epoch-wise mean accuracy. Triplets are now separated according to how their predictability changed between Sequence A and B. X-axis indicates Epochs, y-axis indicates median accuracies (in %). Participants showed a successful acquisition of new implicit knowledge as they started to differentiate between ‘L H’ and ‘L L’ triplets during exposure to Sequence B during Epoch 5 but contrary to RTs, this difference in accuracy was not retained during Epoch 6, where the original sequence was reintroduced. Participants also showed a resistance of old implicit knowledge to interference as they retained the performance difference between ‘H L’ and ‘L L’ triplets even after exposure to Sequence B. In all subfigures, points indicate means, error bars indicate 1 SEM.

### Impact on old knowledge

Trial-level RT (log transformed) or accuracy (0: incorrect vs 1: correct) were used as the outcome variables in GLMMs, containing fixed effects of Epoch (Factor: 4, 5, 6), Triplet type (Factor: H L, L L), subjective cSES, Harshness, Unpredictability, and Age and their higher order interactions, with participant-specific correlated intercepts and slopes for the within-subjects factor of Epoch. Full list of fixed and random effects parameters can be found in Supplementary Table S5 and S6, for RT and accuracy, respectively.

In the RT model, there was a statistically significant main effect of Epoch (F_(2,365.11)_ = 99.41, p < .001) and Triplet type (F_(1,195586.83)_ = 484.72, p < .001), but no significant effect of the Epoch x Triplet type interaction, which indicated slower responses after exposure to the new sequence, and a retention of implicit knowledge of the old sequence, despite exposure to the new sequence at the group level (Supplementary Text S2, Table S5 and Figure 4A). No main effect or interaction involving subjective cSES, Harshness or Unpredictability was statistically significant, thus there was no support for the either dimension of childhood adversity impacting the resistance of implicit SL or visuomotor performance to interference during the interference phase of the experiment, as assessed by RT.

In the accuracy model, there was a statistically significant main effect of Epoch (*χ^2^* (2) = 9.34, p = .009) and Triplet type (*χ^2^* (1) = 217.58, p < .001), and significant the Epoch x Triplet type interaction (*χ^2^* (2) = 22.27, p < .001), which indicated less accurate responses after exposure to the new sequence, and a disruption of implicit knowledge of the old sequence, after exposure to the new sequence at the group level (Supplementary Text S2, Table S6 and Figure 4B). There was a statistically significant Epoch x subjective cSES interaction, which stemmed from a different trajectory of overall performance in individuals with high versus low cSES. Low cSES individuals showed a decrease in accuracy in the final epoch (Epoch 4: 91.9%, 95% CI = [91.3, 92.6]; Epoch 5: 91.5%, 95% CI = [90.8, 92.2]; Epoch 6: 90.8%, 95% CI = [90.0, 91.4]; Epoch 4-5 contrast p = .594, Epoch 4-6 contrast p = .001, Epoch 5-6 contrast p = .086), whereas high cSES individuals showed better overall accuracy and no differences between epochs (Epoch 4: 92.6%, 95% CI = [91.8, 93.3]; Epoch 5: 91.7%, 95% CI = [90.8, 92.6]; Epoch 6: 92.6%, 95% CI = [91.8, 93.3]; Epoch 4-5 contrast p = .168, Epoch 4-6 contrast p > .999, Epoch 5-6 contrast p = .133).

No main effect or interaction involving either Harshness or Unpredictability was statistically significant, thus there was no evidence for either dimension of childhood adversity impacting visuomotor performance or the resistance of implicit SL to interference during the interference phase of the experiment, as assessed by accuracy.

### Impact on new knowledge

Trial-level RT (log transformed) or accuracy (0: incorrect vs 1: correct) were used as the outcome variables in GLMMs, containing fixed effects of Epoch (Factor: 4, 5, 6), Triplet type (Factor: L H, L L), subjective cSES, Harshness, Unpredictability, and Age and their higher order interactions, with participant-specific correlated intercepts and slopes for the within-subjects factor of Epoch. Full list of fixed and random effects parameters can be found in Supplementary Table S7 and S8, for RT and accuracy, respectively.

In the RT model, there was a statistically significant main effect of Epoch (F_(2,354.19)_ = 87.77, p < .001) and Triplet type (F_(1,137396.15)_ = 39.15, p < .001), but no significant effect of the Epoch x Triplet type interaction, which indicated slower responses after exposure to the new sequence, and acquisition of the new regularity at the group level (Supplementary Text S2, Table S7 and Figure 4A). There was a statistically significant subjective cSES x Triplet type interaction (F_(1,137417.54)_ = 8.43, p = .027). This stemmed from a lack of significant ‘L H – L L’ difference in low cSES individuals (Figure 5C, learning score: 0.82 ms, L H: 363 ms, 95% CI = [357, 369]; L L: 364 ms, 95% CI = [358, 370], p = .204), while this effect was observed in high cSES individuals (learning score: 4.34 ms, L H: 361 ms, 95% CI = [353, 369]; L L: 365 ms, 95% CI = [357, 373], p < .001). This effect indicates a detrimental effect of current adversity on the ability to acquire novel environmental contingencies when they conflict with initially established ones.

**Figure 5.**
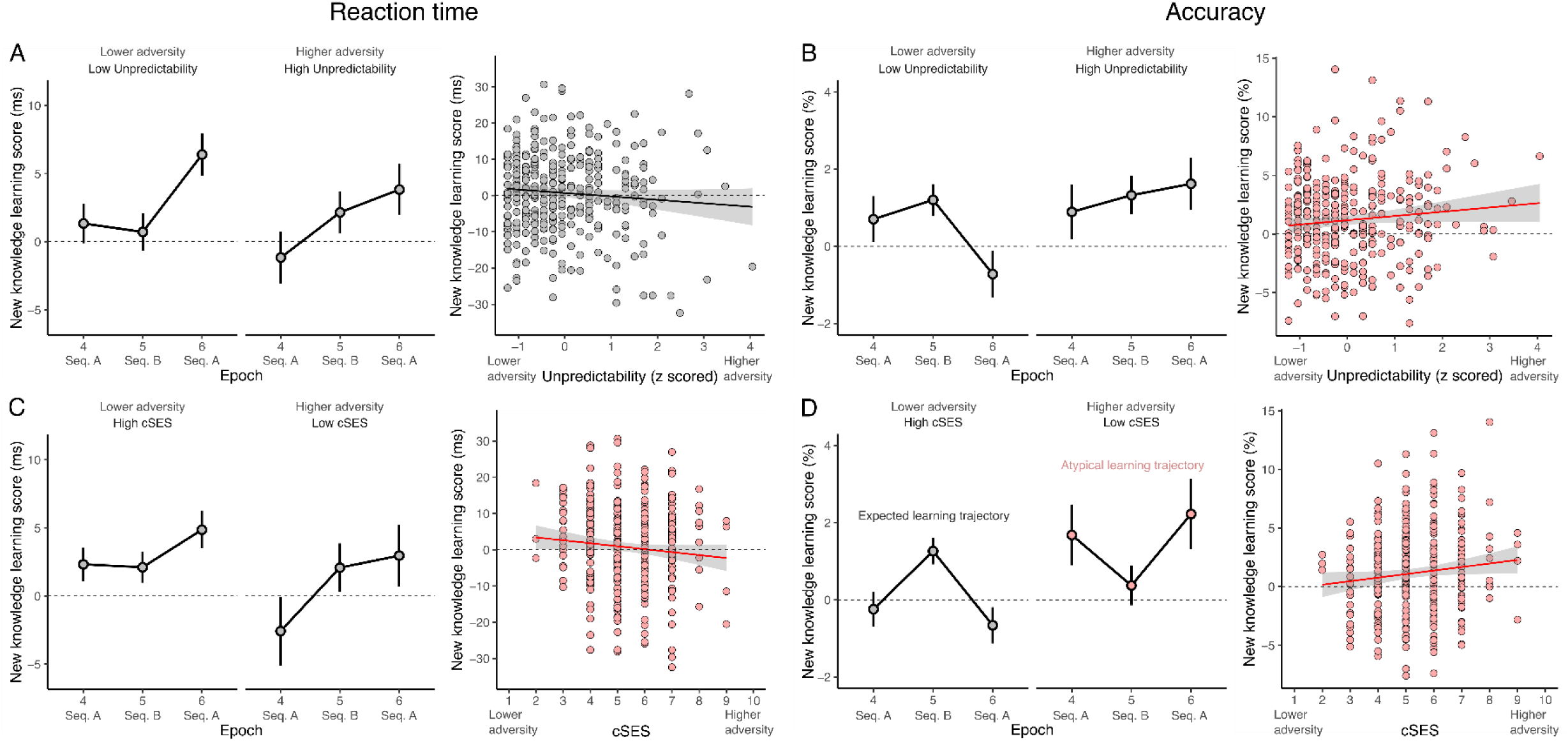
Effect of childhood unpredictability and current SES on the acquisition of new knowledge during the interference phase. **A)** Epoch-wise (left) and interference phase (right) average new knowledge learning scores (RT difference between ‘L L’ and ‘L H’ triplets) as a function of childhood unpredictability. Childhood unpredictability was unrelated to both the trajectory of learning and the overall level reached. **B)** Epoch-wise (left) and interference phase (right) average new knowledge learning scores (accuracy difference between ‘L L’ and ‘L H’ triplets) as a function of childhood unpredictability. Individuals with higher childhood unpredictability exhibited stronger learning overall than individuals with lower childhood unpredictability. **C)** Epoch-wise (left) and interference phase (right) average new knowledge learning scores (RT difference between ‘L L’ and ‘L H’ triplets) as a function of current SES. Individuals with lower current SES (thus higher adversity) exhibited weaker learning overall than individuals with higher current SES (thus lower adversity). **D)** Epoch-wise (left) and interference phase (right) average new knowledge learning scores (accuracy difference between ‘L L’ and ‘L H’ triplets) as a function of current SES. Individuals with lower current SES (thus higher adversity) exhibited increased learning by the final epoch, but also an atypical learning pattern in which their learning scores decreased during the epoch when the new sequence is introduced, in contrast with the expected trajectory observed in higher current SES (thus lower adversity) individuals. On all left figures, points indicate means, error bars indicate 1 SEM. On all right figures, points are subject specific means, line is line of best linear fit, with shading indicating 95% CI. Subfigures that correspond to statistically significant effects in the LMM are highlighted by pink.

With no main effect or interaction involving either Harshness or Unpredictability being statistically significant, there was no support for the hypothesis that either dimension of childhood adversity impacted visuomotor performance or the implicit acquisition of new statistical regularities during the interference phase of the experiment, as assessed by RT.

In the accuracy model, there was a statistically significant main effect of Epoch (*χ^2^* (2) = 43.67, p < .001) and a significant Epoch x Triplet type interaction (*χ^2^* (2) = 14.09, p < .001), which indicated less accurate responses after exposure to the new sequence, and acquisition of the new sequence at the group level (Supplementary Text S2, Table S8 and Figure 4B). There were statistically significant Triplet Type x subjective cSES (*χ^2^* (1) = 11.60, p < .001) and Triplet Type x subjective cSES x Epoch interactions (*χ^2^* (2) = 12.03, p = .002). These effects stemmed from the atypical learning pattern in individuals with lower cSES (Figure 5D, learning score in Epoch 4: 1.38%, 95% CI = [0.35, 2.42]; in Epoch 5: 0.85%, 95% CI = [0.06, 1.62]; in Epoch 6: 1.53%, 95% CI = [0.43, 2.63]; Epoch 4-5 contrast p = .798, Epoch 4-6 contrast p = .996, Epoch 5-6 contrast p = .681), which was contrasted by the expected learning trajectory of successfully acquiring the new sequence during epoch 5 in individuals with higher cSES (learning score in Epoch 4: -1.40%, 95% CI = [-2.69, -0.09]; in Epoch 5: 1.36%, 95% CI = [0.37, 2.35]; in Epoch 6: -1.95%, 95% CI = [-3.29, -0.61]; Epoch 4-5 contrast p = .003, Epoch 4-6 contrast p = .911, Epoch 5-6 contrast p < .001).

There was also a statistically significant Triplet type x Unpredictability interaction (*χ^2^*(2) = 12.03, p = .002), which stemmed from a lack of significant ‘L H – L L’ difference in low unpredictability individuals (Figure 5B, learning score: 0.07%, L H: 91.0%, 95% CI = [90.3, 91.5]; L L: 90.9%, 95% CI = [90.2, 91.5], p = .785), while this effect was observed in high unpredictability individuals (learning score: 0.9%, L H: 91.1%, 95% CI = [90.5, 91.6]; L L: 90.2%, 95% CI = [89.5, 90.8], p < .001).

No main effect or interaction involving Harshness was statistically significant. Overall, these results indicate a facilitatory effect of early unpredictability and a detrimental effect of current adversity on the ability to acquire novel environmental contingencies when they conflict with initially established ones, as assessed by accuracy.

### Sensitivity analysis

In a set of sensitivity analyses, we added Parental education as a control variable, to confirm that our results with respect to subjective adversity hold when we incorporate a more objective measurement. These models thus had the general form:

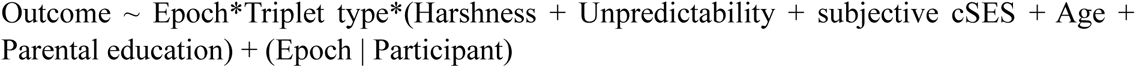

All effects reported in the main models remained robust in all of these sensitivity models (Supplementary Tables S9 to S16).

## Discussion

We examined the associations between variability in exposure to multiple dimensions of early life adversity and current socio-economic status on multiple phases of implicit SL of individuals, in an online sample. We hypothesized that adversity would leave implicit SL ability intact, in stark contrast with its detrimental effects on explicit cognitive processes, such as executive functions (Johnson et al., 2021; Lund et al., 2022). Our results supported our hypothesis, while also revealing a dissociation between the effects of childhood and current forms of adversity. Current adversity, operationalised as subjective current socio-economic status (cSES), had detrimental effects on both implicit SL of novel regularities and general visuo-motor performance in the interference phase, despite being weakly associated with better initial visuomotor performance. Childhood unpredictability, on the other hand, left both visuo- motor performance and implicit SL ability mostly intact, and it was even associated with better implicit SL ability, as shown by the quicker reach of asymptotic performance during the learning phase and the relatively more successful acquisition of novel regularities in the interference phase. Childhood harshness had no associations with task performance. Interestingly, results were not consistent between RT and accuracy, with childhood unpredictability’s SL enhancing effect being apparent in RTs during the learning phase, but in accuracy during the interference phase. Dissociation between RT and accuracy have been reported before in the implicit SL literature (Janacsek et al., 2012; Kiss et al., 2022), and could suggest different underlying neurocognitive mechanisms (Hedge et al., 2018; Mulder & van Maanen, 2013; Prinzmetal et al., 2005; van Ede et al., 2012), but this possibility remains to be fully explored.

These results are in line with the emerging ‘hidden talents’ framework, which highlights the potential for adversity and chronic stress to influence cognition in adaptive ways, leading to complex patterns of diminished, intact and enhanced abilities in a single individual (Ellis, Abrams, et al., 2022; Frankenhuis, Vries, et al., 2020; Frankenhuis & Nettle, 2020). This complexity is evident in our results as well. While childhood unpredictability was associated with faster implicit SL during acquisition of the initial regularity and larger implicit SL during the learning of a novel, conflicting one, childhood harshness was not associated with either visuo-motor performance of implicit SL. On the other hand, subjective current SES proved to have rather complex effects, facilitating quicker RT improvements during acquisition, but worse learning of a novel regularity and a rather atypical learning trajectory. There are studies showing similar effects, for example, better performance by low SES individuals in a categorization task during financial stress (Dang et al., 2016), and intact probabilistic learning and caudate nucleus volumes in low SES individuals, contrasted by impaired working memory and smaller hippocampus volumes (Leonard et al., 2015). Childhood unpredictability has also been found to be related to enhanced task switching and working memory, especially in uncertain contexts (Mittal et al., 2015; Young et al., 2018, 2022). The relationship between task switching, working memory and executive functions in general and implicit SL is not entirely clear (Janacsek & Nemeth, 2013, 2022; Park et al., 2020; Pedraza et al., 2024). However, an enhanced cognitive flexibility could enable individuals to build and decide between competing internal models of the environment more efficiently. This then could underlie the observed facilitatory effects on initial learning, and especially rewiring. Overall, our results reveal the importance of separating current and childhood adversity, as well as different phases of learning, when studying cognition from a ‘hidden talents’ perspective. Our results could equally be interpreted in light of evolutionary-developmental theories of stress adaptation, which suggest that early life stress should have highly non-linear effects on development, with moderate levels of stress leading to less stress-reactive profiles (Boyce & Ellis, 2005; Del Giudice, 2014; Ellis et al., 2005; Ellis & Del Giudice, 2019). Given our focus on levels of adversity within the ‘typical’ range, our results are consistent with a theory in which moderate levels of unpredictability are associated with increased implicit learning of regularities, whereas lower than typical, and higher than typical levels of it are associated with diminished implicit learning.

What evolutionary advantage might increased implicit SL bring to individuals in moderately high unpredictability contexts? We propose that this cognitive enhancement might be understood as a complementary mechanism to working memory and task switching for mitigating the threats that stochastic variations of the environments pose (Mittal et al., 2015; Young et al., 2018, 2022). Whereas working memory and task switching enable the tracking of rapidly fluctuating environmental contingencies in a *supervised* and *conscious* manner, implicit SL might achieve the same by picking up on subtle environmental regularities in an *unsupervised* and *non-conscious* manner and shaping behaviour, perception, and other aspects of cognition implicitly (Conway, 2020; Janacsek & Nemeth, 2022). Ultimately, both explicit and implicit regularity detection help individuals to predict important variables, such as threats, the emotional states of others, and social dynamics (Plate et al., 2022; Woodard et al., 2022). Faster SL suggests that participants with greater childhood unpredictability exposure are more efficient precisely in finding structure in the information stream (the environment). This extraction of probabilities can then form the basis for building predictive models in the brain (Éltető et al., 2022; Janacsek & Nemeth, 2012). One reason why increased implicit tracking of contingencies might be preferred in some situations over explicit tracking is its relatively less cognitively demanding nature (Buabang et al., 2024; Hardwick et al., 2019; Schwabe & Wolf, 2013). Given energetic constraints, the brain might shift between explicit and implicit systems in a resource-efficient manner (Lieder & Griffiths, 2020).

As for how the brain detects cues during childhood to guide energy allocation decisions (presumably including whether to invest in enhanced implicit statistical learning systems), currently there are two, not mutually exclusive hypotheses (Young et al., 2020). Under the ‘ancestral cue’ approach, proximate neurocognitive mechanisms are ‘wired’ to detect specific experiences, that have reliably served as cues to environmental states (Ellis et al., 2009). For example, for developing children in ancestral environments, parental instability might have been a reliable indicator of environmental instability they could expect in the future. Under the ‘statistical learning’ approach, proximate mechanisms learn internal models of environmental states in a general way by integrating many different sources of information, continuously through development (Frankenhuis et al., 2019). We are not able to distinguish these potential explanations, and it is likely that both are at play (Li et al., 2023). Nevertheless, our measurement of adversity seems more aligned with the ‘statistical learning’ perspective, as while some items in our questionnaire could reasonably be ‘ancestral cues’ (e.g., frequent residential change, mortality), most are unlikely to have been frequently recurring and reliable cues to mortality-morbidity through human evolution. In addition, a possibility remains that the effects reported herein are produced not by developmental adaptation to early life environments, but only by current stress levels (Schwabe & Wolf, 2013). We believe this explanation is unlikely to account for the results for two reasons. Firstly, while we did not measure current subjective stress, both subjective and objective (parental education) current SES were measured, and both are known to be associated with stress (Chandola & Marmot, 2011) and subjective well-being (Tan et al., 2020). Thus, the inclusion of them as covariates in our main and sensitivity models should have adjusted for current stress levels to some degree. In addition, the association between these measures and early life adversity were quite low, suggesting that, at least in our sample, they capture quite different components of lifetime environmental stress.

Regarding potential neurobiological mechanisms, there is evidence that acute stress causes a shift towards the striatal, model-free, procedural memory system, from the hippocampal, model-based, declarative memory system (Otto et al., 2013; Schwabe & Wolf, 2012, 2013). Some recent studies also suggest facilitatory effects of acute stress on implicit SL (Sherman, Huang, et al., 2024), including a study specifically using the ASRT task (Tóth-Fáber et al., 2021). A possibility is that chronic stress, such as imposed by unpredictable variations of the environment during development, is associated with a more general and long-term shift in the balance between these memory systems, a hypothesis also put forward by others (Ellis, Abrams, et al., 2022). While this is possible, an enhancement of both implicit SL (as suggested by our study) and explicit working memory and task switching abilities (as reported in the literature, Mittal et al., 2015; Young et al., 2018, 2022) argue against a general shift from top-down towards bottom-up cognition. More evidence from longitudinal human studies and neural data will be necessary to shed light on this issue.

Our study has a number of important limitations that need to be mentioned and addressed by future work. Firstly, the effect sizes of the adversity variables were relatively small, and their practical importance is unclear, given the abstract and non-ecological nature of the task. This is especially pertinent given recent evidence that there can be drastic differences in adversity-exposed individuals’ performance between abstract, non-ecological and concrete, ecological contexts (Young et al., 2022; but see Duquennois, 2022; Muskens et al., 2024). However, we note that small effect sizes can be associated with quite large practical effects, especially when the psychological process they concern is cumulative, like the hypothesized developmental effects we were interested in here (Abelson, 1985; Funder & Ozer, 2019). Given the association between implicit SL ability and many cognitive (Graybiel, 2008; Ullman, 2004), social (D. Baldwin et al., 2008; Ruffman et al., 2012), and motor skills (Hallgató et al., 2013; Verburgh et al., 2016), even small increases in implicit SL ability can scaffold a host of further advantages throughout the lifetime, if they emerge early. Similarly, we can easily imagine how even slightly faster estimation of environmental contingencies (e.g., threat cues) could translate to meaningful fitness increases over an entire lifetime in ancestral environments. Moreover, the study we cite above (Young et al., 2022) showed that performance of more adversity exposed individuals increased with more ecological stimuli, suggesting that if anything, the effects we see with relatively abstract stimuli are underestimates (but see Duquennois, 2022; Muskens et al., 2024). Nevertheless, future studies should consider using more ecologically valid tasks. Secondly, it is surprising that in our sample, there seemed to be only weak associations between early life adversity dimensions and EF measures, especially given previous evidence of increased cognitive flexibility (Mittal et al., 2015; Young et al., 2022). This could stem from our undersampling of individuals exposed to the highest levels of harshness and unpredictability, insufficient variability in early life adversity in our sample, or shortcomings of the WCST perseverative error score as a measure of cognitive flexibility (Barceló, 2001; Nyhus & Barceló, 2009). The robust associations between both of our adversity composites and multiple dimensions of psychopathology and parental education are nevertheless reassuring, regarding the validity of our operationalisation. The lack of correlation between early life adversity dimensions and subjective current SES is similarly surprising, and could be due to increasing social mobility in Hungarian society (though this is not as high as one would initially expect, see Bukowski et al., 2022; and Róbert & Bukodi, 2004), or reflect the relatively simplistic nature of our subjective cSES measure compared to the more extensive childhood adversity assessment. Given the need to accurately assess both early life and current adversity for ‘hidden talents’ research (Ellis, Abrams, et al., 2022; Frankenhuis, Young, et al., 2020), this present study should be interpreted with the required degree of caution, and future work should use a more extensive assessment of current adversity, that is matched to the childhood measures to the extent it is possible. Finally, our use of retrospective and subjective self-report measures of adversity could be taken as a limitation, given the relatively low correspondence between retrospectively and prospectively reported adverse childhood experiences (J. R. Baldwin et al., 2019; Reuben et al., 2016). While we highly agree that it is important for the ‘hidden talents’ research programme to validate its findings with objective measures and prospective, longitudinal samples, we note that concordance between subjective and objective measures is not always low (Naicker et al., 2021; Patten et al., 2015), and there is plenty of evidence indicating that subjective adversity indicators are often better predictors of functional outcomes than objective ones (Danese & Widom, 2020; Francis et al., 2023).

In conclusion, we found that the effects of current and early life adversity dimensions on implicit SL are quite distinct, with current forms of adversity (i.e., SES) being associated with generally detrimental, and early life adversity (i.e., moderate levels of unpredictability) with generally beneficial effects. This dissociation further highlights the importance of distinguishing multiple dimensions and exposure windows when studying the effects of chronic stressors on cognition, and the facilitatory effects stand in stark contrast with the widely reported negative impacts on other areas of cognition, stressing the value of the emerging ‘hidden talents’ framework.

## Acknowledgements

The authors would like to thank all members of the Cognitive Variability Lab of the École Normale Supérieure for their helpful comments.

## Data availability

The data that support the findings of this study are openly available in https://osf.io/gmr35/?view_only=c4da4956159f46128007e9c6cef3fba9. (This is currently the anonymized link to the private project, which will be replaced by the public project’s DOI link upon acceptance).

## CRediT author statement

Bence C. Farkas: Conceptualization, Methodology, Software, Validation, Formal analysis, Writing – Original Draft, Writing – Review & Editing, Visualization

Bianka Brezóczki: Conceptualization, Software, Validation, Investigation, Data Curation, Writing – Review & Editing

Teodóra Vékony: Conceptualization, Software, Validation, Investigation, Resources, Data Curation, Writing – Review & Editing

Pierre O. Jacquet: Methodology, Writing – Review & Editing, Supervision

Dezso Nemeth: Conceptualization, Validation, Resources, Writing – Review & Editing, Supervision, Project administration, Funding acquisition

