## Supplementary Materials for "Faster adult implicit probabilistic statistical learning following childhood adversity"

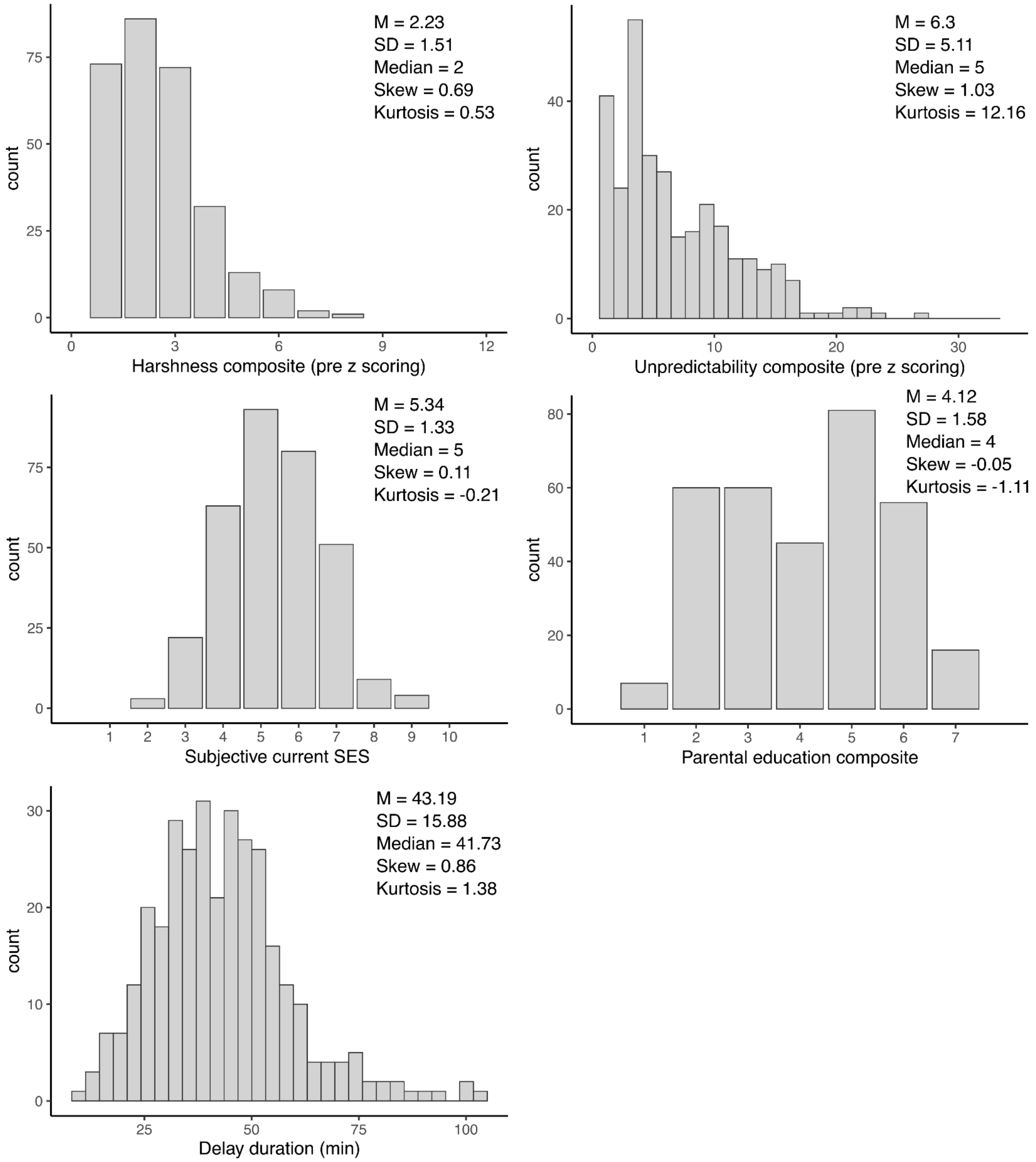


**Supplementary Figure S1. Histograms and descriptive statistics of the distributions of primary variables.**

**Supplementary Text S1**

**Psychopathology and Executive function measures**

**Adult ADHD Self-Report Scale (ASRS)**

Adult attention-deficit/hyperactivity disorder (ADHD) symptoms were measured by the Adult ADHD Self-Report Scale (ASRS), a short screening tool intended to capture adult ADHD symptoms in the general population (Kessler et al., 2005). The ASRS includes 18 questions about frequency of recent DSM-IV Criterion A symptoms of adult ADHD, with items such as *"How often do you make careless mistakes when you have to work on a boring or difficult project ?* and *"How often do you feel restless or fidgety ?"*. Each question asks how often a given symptom occurred over the past 6 months on a 0–4 scale with responses of never (0), rarely (1), sometimes (2), often (3), and very often (4). To operationalise ADHD symptoms, a sum score was used.

**Obsessive-Compulsive Inventory – Revised (OCI-R)**

Obsessive-Compulsive Disorder (OCD) symptoms were measured by the Obsessive-Compulsive Inventory – Revised (OCI-R) (Foa et al., 2002), an 18-item self-report scale. The OCI-R includes items such as *"I check things more often than necessary."* and *"I frequently get nasty thoughts and have difficulty in getting rid of them."*. Each item asks how much a given experience distressed or bothered the individual in the past month, on a 0-4 scale with responses of not at all (0), a little (1), moderately (2), a lot (3), and extremely (4). To operationalise OCD symptoms a sum score was used.

**Autism-Spectrum Quotient (AQ)**

Adult Autism spectrum disorder (ASD) symptoms were measured by the Autism-Spectrum Quotient (AQ), a self-report instrument of autistic traits in the general population (Baron-Cohen et al., 2001). It comprises 50 questions, such as "I prefer to do things with others rather than on my own." and "I am fascinated by numbers.". Each question asks to what degree the participant thinks a given autistic trait or behaviour describes them, on a 4-point Likert scale, with responses of "definitely agree", "slightly agree". "slightly disagree', and "definitely disagree". Each item scores 1 point if the individual reports the autistic trait mildly or strongly, with approximately half of the items being reverse coded.

**Eating attitudes (EAT)**

The Eating Attitude Test (EAT) is a self-report scale designed to assess disordered eating attitudes commonly linked to anorexia and bulimia (Garner et al., 1982). In our study, we utilized the 26-item shortened version, with participants responding on a 6-point Likert scale. Example items include: "I am terrified about being overweight."; "I am aware of the calorie content of foods that I eat."; "I feel that food controls my life.".

**Hypomania checklist (HCL)**

Hypomania symptoms were measured by the Hypomania checklist (HCL), a self-report scale intended to identify hypomanic symptoms and differentiate Major Depressive Disorder from Bipolar Disorder (Angst et al., 2005). It is a self-report questionnaire with 32 items. This tool evaluates two factors: active/elated mood and irritable/risk-taking tendencies. Example items include: “My thoughts dart from topic to topic.”; “I am more impatient and get irritated more easily.”; “I talk more.”.

**Brief Multidimensional Schizotypy Scale (MSS-B)**

To measure schizotypal traits, we used the Brief Multidimensional Schizotypy Scale (MSS-B) of Gross et al. (2018). This questionnaire assesses positive, negative, and disorganized schizotypy symptoms through 38 items. Participants respond with "yes" or "no," and higher scores reflect greater severity of schizotypal traits. Example items include: “Sometimes I felt as if strangers were reading my thoughts.”; “My thoughts and behavior are almost always disorganized.”; “I believe there are secret signs in the world; you just need to know how to find them.”.

**Digit Span (DSPAN) task**

We measured phonological short-term memory capacity using a computerised version of the digit span task (Isaacs & Vargha-Khadem, 1989). In this task, participants were presented with and then typed a list of digits enunciated by an experimenter. The task comprised seven levels of difficulty ranging from three to nine items. Each level comprised four different lists. If participants correctly recalled all items for at least three out of the four lists, they were permitted to move up a level. If participants could not recall at least three lists from the four lists correctly, the task ended. Memory span was operationalised as the level (three to nine) at which the participant was still able to recall three from the four lists correctly. A higher memory span score is indicative of a better phonological short-term memory capacity.

**1-BACK task**

Another measure of working memory was the 1-BACK task (Kirchner, 1958). In the task, letters are presented on the screen consecutively. The users' task is to press the "J" key on the keyboard for the target elements, and the "F" for the non-target elements. The target stimulus differs between the different levels of the task. The task begins with written instructions. Before the two blocks, a 10-trial practice is implemented. During the practice, the users receive feedback about their answer ("Correct", "Wrong", "You did not respond"). After the practice, the two 50-trial long blocks begin. Between blocks, a self-paced rest period is inserted. After the end of the second block, the users receive feedback about their overall success rate and reaction time. Working memory capacity was operationalised by d prime (d’):

$$d' = Z(\text{hit rate}) - Z(\text{false alarm rate})$$

A higher d’ indicates greater working memory capacity.

**Go / No-go task (GNG)**

We measured cognitive inhibition with a computerised version of the Go / No-go task (Bezdjian et al., 2009). In this task, participants were instructed to respond to certain stimuli (“go” stimuli) by clicking on a button as fast as possible and to refrain from clicking on other stimuli (“no-go” stimuli). Participants were presented with a 2×2 array with four blue stars (one in the centre of each square of the array). Every 1500 ms, a stimulus (the letter P or R) would appear for 500 ms in the place of one of the blue stars. For the first half of the task, the letter P would be the “go” stimulus and the letter R would be the “no-go”. This rule would be then inverted in the second half of the task. Participants completed 320 trials in total, with an 80:20 ratio of “go” and “no-go” trials. Cognitive inhibition capacity was operationalised by d prime (d’), using the same formula as for 1-BACK. A higher d’ indicates better cognitive inhibition.

**Card Sorting Task (WCST)**

We measured cognitive flexibility using a computerised version of the Berg Card Sorting Task (Berg, 1948; Fox et al., 2013). In this test, a set of four cards were presented to the participant. Each card had three characteristics: colour, shape and number of items. Participants were told to match new cards to the cards on the top of the screen according to one of the three characteristics, but they were not told which one. Instead, they received feedback about whether each choice they made was right or wrong, and could use this to find the correct rule. Cognitive flexibility was operationalised as the fraction of perseverative errors, which was then subtracted from 100 so that higher numbers reflected better cognitive flexibility to aid interpretation.

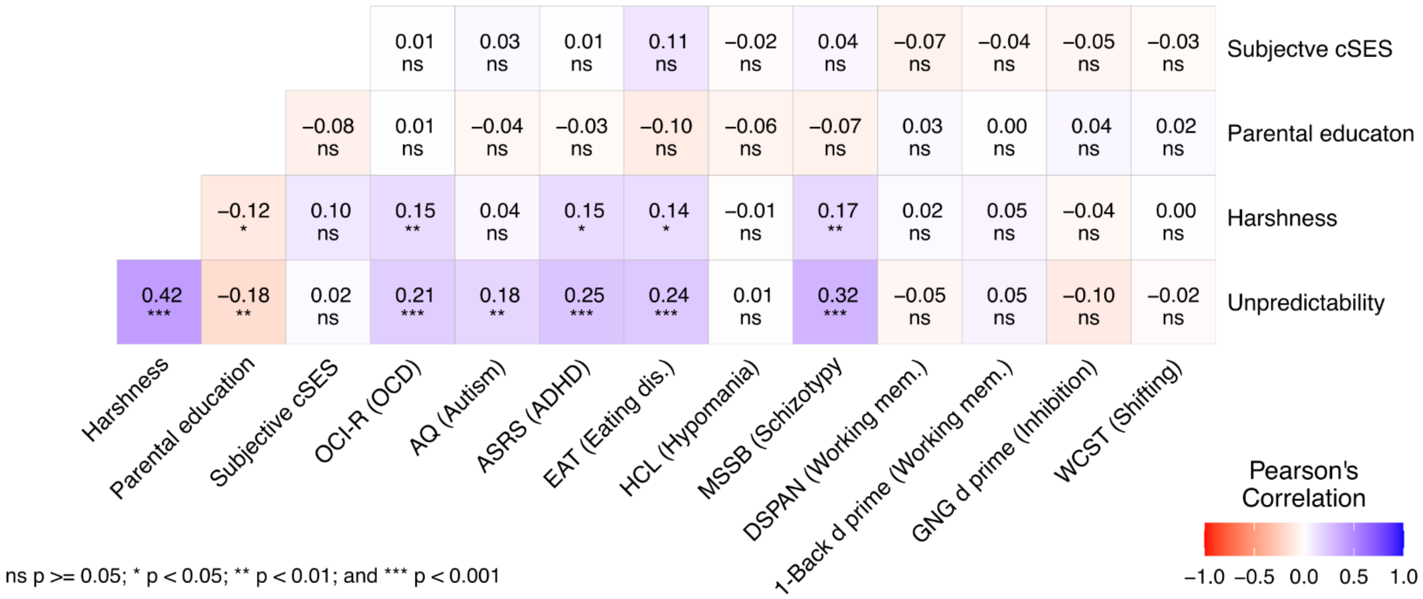


**Supplementary Figure S2. Bivariate correlations between our adversity measures and adult psychopathology and executive functions.** Coefficients are Pearson's r, with their direction and strength also indicated by colour. Each variable is coded such that higher values correspond to higher number of symptoms, better executive functions, greater adversity, worse subjective cSES and higher parental education.

#### Supplementary Text S2

#### Trajectories of visuomotor performance and implicit SL

##### Learning

The RT model resulted in a statistically significant main effect of Epoch ($F_{(2,337.39)} = 55.45$, $p < .001$). Participants became faster in subsequent epochs (Epoch 1: 391 ms, 95% CI = [386, 396]; Epoch 2: 389 ms, 95% CI = [384, 394]; Epoch 3: 381 ms, 95% CI = [376, 385]; Epoch 1-2 contrast p = .037, Epoch 1-3 contrast p < .001, Epoch 2-3 contrast p < .001). This effect indicates that on average subjects showed general visuomotor performance improvement, i.e., they became faster overall as the task went on, irrespective of triplet probabilities. There was a statistically significant main effect of Triplet type ($F_{(1,289785.18)} = 295.40$, $p < .001$), with faster reaction times for high-probability triplets than for low-probability triplets, indicating implicit SL (High: 384 ms, 95% CI = [380, 389]; Low: 390 ms, 95% CI = [385, 394]). There was also a statistically significant Epoch x Triplet type interaction ($F_{(2,289788.17)} = 20.17$, $p < .001$), which stemmed from increasing implicit SL with time (learning score in Epoch 1: 2.88 ms, 95% CI = [1.85, 3.91]; in Epoch 2: 5.40 ms, 95% CI = [6.43, 4.36]; in Epoch 3: 7.58 ms, 95% CI = [6.56, 8.60]; Epoch 1-2 contrast p = .002, Epoch 1-3 contrast p < .001, Epoch 2-3 contrast p = .010).

There was a statistically significant main effect of Age ($F_{(1,320.55)} = 9.83$, $p = .002$), with overall slower responses in relatively older participants (b = 1.004, 95% CI = [1.001 – 1.006]). There was a statistically significant Epoch x Age interaction ($F_{(2,335.51)} = 4.31$, $p = .014$), with a larger change from Epoch 1 to 2 in older participants (RT in Epoch 1: 399 ms, 95% CI = [391, 406]; in Epoch 2: 395 ms, 95% CI = [388, 403]; in Epoch 3: 390 ms, 95% CI = [383, 396]; Epoch 1-2 contrast p = .062, Epoch 1-3 contrast p < .001, Epoch 2-3 contrast p < .001), compared to younger participants (RT in Epoch 1: 384 ms, 95% CI = [377, 391]; in Epoch 2: 383 ms, 95% CI = [376, 389]; in Epoch 3: 372 ms, 95% CI = [366, 378]; Epoch 1-2 contrast p = .550, Epoch 1-3 contrast p < .001, Epoch 2-3 contrast p < .001). There was also a Triplet type x Age interaction ($F_{(1,289787.73)} = 4.61$, $p = .032$). This stemmed relatively higher learning scores in older individuals (learning score: 6.08 ms, High: 392 ms, 95% CI = [385, 398]; Low: 398 ms, 95% CI = [391, 405], p < .001), compared to younger ones (learning score: 4.57 ms, High: 377 ms, 95% CI = [371, 384]; Low: 382 ms, 95% CI = [375, 388], p < .001).

The accuracy model resulted in a statistically significant main effect of Epoch ($\chi^2$ (2) = 73.92, p < .001). Participants became less accurate in subsequent epochs (Epoch 1: 93.1%, 95% CI = [92.6, 93.5]; Epoch 2: 91.7 %, 95% CI = [91.3, 92.1]; Epoch 3: 91.3 %, 95% CI = [90.9, 91.7]; Epoch 1-2 contrast p < .001, Epoch 1-3 contrast p < .001, Epoch 2-3 contrast p = .030). There was a statistically significant main effect of Triplet type ($\chi^2$ (1) = 337.38, p < .001), with higher accuracy for high-probability triplets than for low-probability triplets, indicating implicit SL (High: 93.0%, 95% CI = [92.6, 93.73]; Low: 91.1%, 95% CI = [90.6, 91.5]). There was also a statistically significant Epoch x Triplet type interaction ($\chi^2$ (2) = 38.10, p < .001), which stemmed from increasing implicit SL with time (learning score in Epoch 1: 1.08%, 95% CI = [0.74, 1.41]; in Epoch 2: 1.82%, 95% CI = [1.44, 2.19]; in Epoch 3: 3.04%, 95% CI = [2.63, 3.44]; Epoch 1-2 contrast p = .010, Epoch 1-3 contrast p < .001, Epoch 2-3 contrast p < .001).

##### Consolidation

The RT model resulted in a statistically significant main effect of Epoch ($F_{(1,331.46)} = 499.31$, $p < .001$). Participants were overall faster after the delay period (Epoch 3: 377 ms, 95% CI =

[373, 381]; Epoch 4: 354 ms, 95% CI = [350, 357]). There was a statistically significant main effect of Triplet type ($F_{(1,193144.75)}$ = 472.63, p < .001), with faster reaction times for high-probability triplets than for low-probability triplets (High: 361 ms, 95% CI = [357, 365]; Low: 369 ms, 95% CI = [365, 373]). The lack of a significant Epoch x Triplet type interaction indicates that there was no improvement after the delay in implicit SL in terms of RT.

There was again a statistically significant effect of Age ($F_{(1,319.72)}$ = 18.13, p < .001), with overall slower responses in relatively older participants (b = 1.005, 95% CI = [1.003 – 1.007]), as well as a Triplet type x Age interaction ($F_{(1,193148.88)}$ = 4.17, p = .041), with relatively higher learning scores in older individuals (learning score: 8.18 ms, High: 370 ms, 95% CI = [364, 376]; Low: 378 ms, 95% CI = [372, 384], p < .001), compared to younger ones (learning score: 6.48 ms, High: 353 ms, 95% CI = [347, 359]; Low: 359 ms, 95% CI = [354, 365], p < .001). There was also a statistically significant main effect of Delay duration ($F_{(1,319.81)}$ = 10.76, p < .001), with slower responses in participants with longer delays (b = 1.001, 95% CI = [1.000 – 1.002]).

The accuracy model resulted in a statistically significant main effect of Epoch ($\chi^2$ (1) = 32.33, p < .001). Participants were overall less accurate after the delay period (Epoch 3: 91.2%, 95% CI = [90.8, 91.6]; Epoch 4: 92.3 %, 95% CI = [91.9, 92.6]). There was a statistically significant main effect of Triplet type ($\chi^2$ (1) = 493.07, p < .001), with higher accuracy for high-probability triplets than for low-probability triplets (High: 93.1%, 95% CI = [92.8, 93.4]; Low: 90.2%, 95% CI = [89.7, 90.6]). The lack of a significant Epoch x Triplet type interaction indicates that there was no improvement after the delay in implicit SL in terms of accuracy.

**Interference**

*Old knowledge*

The RT model resulted in a statistically significant main effect of Epoch ($F_{(2,365.11)}$ = 99.41, p < .001). Participants became overall slower after exposure to the new sequence during Epoch 5 (Epoch 4: 353 ms, 95% CI = [349, 357]; Epoch 5: 363 ms, 95% CI = [359, 367]; Epoch 6: 365 ms, 95% CI = [361, 369]; Epoch 4-5 contrast p < .001, Epoch 4-6 contrast p < .001, Epoch 5-6 contrast p = .029). There was a statistically significant main effect of Triplet type ($F_{(1,195586.83)}$ = 484.72, p < .001), with faster reaction times for ‘H L’ triplets than for ‘L L’ triplets (H L: 356 ms, 95% CI = [352, 360]; L L: 364 ms, 95% CI = [360, 368]). This indicates that overall, participants retained the implicit knowledge of the old sequence, despite exposure to the new sequence. Moreover, the lack of a statistically significant Epoch x Triplet type interaction indicates that this resistance to interference was observed in every Epoch to roughly the same extent.

There was again a statistically significant effect of Age ($F_{(1,321.35)}$ = 16.98, p < .001), with overall slower responses in relatively older participants (b = 1.005, 95% CI = [1.002 – 1.007]), as well as a Triplet type x Age interaction ($F_{(1,195578.64)}$ = 5.39, p = .020), with relatively higher learning scores in older individuals (learning score: 9.21 ms, H L: 364 ms, 95% CI = [358, 370]; Low: 373 ms, 95% CI = [368, 379], p < .001), compared to younger ones (learning score: 7.13 ms, H L: 348 ms, 95% CI = [343, 354]; Low: 356 ms, 95% CI = [350, 361], p < .001).

The accuracy model resulted in a statistically significant main effect of Epoch ($\chi^2$ (2) = 9.34, p = .009), with participants becoming overall less accurate after exposure to the new sequence during epoch 5 (Epoch 4: 92.2%, 95% CI = [91.8, 92.6]; Epoch 5: 91.6%, 95% CI = [91.2, 92.1]; Epoch 6: 91.6%, 95% CI = [91.1, 92.0]; Epoch 4-5 contrast p = .028, Epoch 4-6 contrast p = .004, Epoch 5-6 contrast p = .987). There was a statistically significant main effect of Triplet type ($\chi^2$ (1) = 217.58, p < .001), with higher accuracy for ‘H L’ triplets than for ‘L L’ triplets (H L: 92.9%, 95% CI = [92.6, 93.2]; L L: 90.6%, 95% CI = [90.1, 91.0]). This indicates that overall, participants retained the implicit knowledge of the old sequence, despite exposure to the new sequence. There was a statistically significant Epoch x Triplet type interaction ($\chi^2$ (2) = 22.27, p < .001), which stemmed from a reduced difference in performance between ‘H L’ and ‘L L’ triplets during exposure to the new sequence in Epoch 5 (learning score in Epoch 4: 2.94%, 95% CI = [2.45, 3.44]; in Epoch 5: 1.17%, 95% CI = [0.55, 1.78]; in Epoch 6: 2.92%, 95% CI = [2.41, 3.43]; Epoch 4-5 contrast p < .001, Epoch 4-6 contrast p > .999, Epoch 5-6 contrast p < .001). This indicates that the introduction of a novel sequence interfered with the previously established implicit knowledge, but also that performance returned to similar levels again when the previously established sequence was reintroduced.

*New knowledge*

The RT model once again captured the effect of exposure to the new sequence and resulted in a statistically significant main effect of Epoch ($F_{(2,354.19)}$ = 87.77, p < .001), with participants becoming overall slower after exposure to the new sequence during epoch 5 (Epoch 4: 356 ms, 95% CI = [352, 360]; Epoch 5: 366 ms, 95% CI = [362, 370]; Epoch 6: 368 ms, 95% CI = [364, 372]; Epoch 4-5 contrast p < .001, Epoch 4-6 contrast p < .001, Epoch 5-6 contrast p = .014). There was a statistically significant main effect of Triplet type ($F_{(1,137396.15)}$ = 39.15, p < .001), with faster reaction times for ‘L H’ triplets than for ‘L L’ triplets (L H: 362 ms, 95% CI = [358, 366]; L L: 364 ms, 95% CI = [360, 368]). This indicates that overall, participants acquired the implicit knowledge of the new sequence, however the extremely small differences suggest that establishing this knowledge proved rather difficult.

There was again a statistically significant effect of Age ($F_{(1,320.71)}$ = 17.49, p < .001), with overall slower responses in relatively older participants (b = 1.005, 95% CI = [1.002 – 1.007]).

The accuracy model resulted in a statistically significant main effect of Epoch ($\chi^2$ (2) = 43.67, p < .001), with participants becoming overall less accurate after exposure to the new sequence during epoch 5 (Epoch 4: 90.8%, 95% CI = [90.2, 91.3]; Epoch 5: 91.6%, 95% CI = [91.2, 92.0]; Epoch 6: 89.9%, 95% CI = [89.4, 90.4]; Epoch 4-5 contrast p = .003, Epoch 4-6 contrast p = .009, Epoch 5-6 contrast p < .001). There was a statistically significant Epoch x Triplet type interaction ($\chi^2$ (2) = 14.09, p < .001), which captured the fact that the ‘L H’ – ‘L L’ difference emerged only during epoch 5 (learning score in Epoch 4: 0.23%, 95% CI = [-0.41, 0.87]; in Epoch 5: 1.06%, 95% CI = [0.58, 1.55]; in Epoch 6: 0.02%, 95% CI = [-0.64, 0.69]; Epoch 4-5 contrast p = .119, Epoch 4-6 contrast p = .961, Epoch 5-6 contrast p = .041).

There was a statistically significant Triplet type x Age interaction ($\chi^2$ (1) = 4.21, p = .040), with relatively higher learning scores in younger individuals (learning score: 0.80 %, L H: 91.0, 95% CI = [90.4, 91.5]; L L: 90.1, 95% CI = [89.5, 90.7], p < .001), compared to older ones (learning score: 0.10 %, L H: 91.0, 95% CI = [90.5, 91.6]; L L: 90.9, 95% CI = [90.3, 91.5], p = .684).

#### Table S1. Learning RT.

| Type III test of effects | *F* | *df* | *p* |
|---|---|---|---|
| **Epoch** | 55.45 | 2, 337.39 | **< .001** |
| **Triplet type** | 295.40 | 1, 289785.18 | **< .001** |
| Harshness | 0.32 | 1, 320.60 | .572 |
| Unpredictability | 1.01 | 1, 320.61 | .316 |
| Subj cSES | 0.04 | 1, 320.58 | .842 |
| **Age** | 9.83 | 1, 320.55 | **.002** |
| **Epoch x Triplet type** | 20.17 | 2, 289788.17 | **< .001** |
| Epoch x Harshness | 0.15 | 2, 337.76 | .859 |
| Epoch x Unpredictability | 0.46 | 2, 338.17 | .633 |
| **Epoch x Subj cSES** | 3.16 | 2, 337.02 | **.044** |
| **Epoch x Age** | 4.31 | 2, 335.51 | **.014** |
| Triplet type x Harshness | 0.00 | 1, 289781.43 | .997 |
| Triplet type x Unpredictability | 0.62 | 1, 289770.51 | .433 |
| Triplet type x Subj cSES | 0.00 | 1, 289777.80 | .994 |
| **Triplet type x Age** | 4.61 | 1, 289787.73 | **.032** |
| Epoch x Triplet type x Harshness | 1.41 | 2, 289783.82 | .245 |
| **Epoch x Triplet type x Unpredictability** | 3.61 | 2, 289784.06 | **.027** |
| Epoch x Triplet type x Subj cSES | 1.02 | 2, 289789.05 | .361 |
| Epoch x Triplet type x Age | 0.02 | 2, 289783.23 | .976 |

| Fixed effects | *exp(b)* | *95% CI* | *t* | *df* | *p* |
|---|---|---|---|---|---|
| **(Intercept)** | 386.819 | 382.070 – 391.626 | 948.997 | 320.596 | **< .001** |
| **Epoch [1]** | 1.011 | 1.008 – 1.015 | 6.570 | 331.371 | **< .001** |
| **Epoch [2]** | 1.006 | 1.003 – 1.008 | 4.659 | 344.214 | **< .001** |
| **Triplet type [High]** | 0.993 | 0.992 – 0.994 | -17.187 | 289785.183 | **< .001** |
| Harshness | 1.004 | 0.990 – 1.017 | 0.566 | 320.604 | .572 |
| Unpredictability | 0.993 | 0.980 – 1.007 | -1.004 | 320.611 | .316 |
| Subj cSES | 1.001 | 0.992 – 1.010 | 0.199 | 320.584 | .842 |
| **Age** | 1.004 | 1.001 – 1.006 | 3.135 | 320.549 | **.002** |
| **Epoch [1] × Triplet type [High]** | 1.003 | 1.002 – 1.004 | 5.739 | 289771.758 | **< .001** |
| Epoch [2] × Triplet type [High] | 1.000 | 0.999 – 1.001 | -0.479 | 289806.080 | .632 |
| Epoch [1] × Harshness | 1.000 | 0.997 – 1.004 | 0.211 | 331.767 | .833 |
| Epoch [2] × Harshness | 1.001 | 0.998 – 1.003 | 0.372 | 344.899 | .710 |
| Epoch [1] × Unpredictability | 1.002 | 0.998 – 1.005 | 0.939 | 331.663 | .348 |
| Epoch [2] × Unpredictability | 1.000 | 0.997 – 1.002 | -0.238 | 345.933 | .812 |
| Epoch [1] × Subj cSES | 1.000 | 0.997 – 1.002 | -0.116 | 331.082 | .908 |
| **Epoch [2] × Subj cSES** | 0.998 | 0.996 – 1.000 | -2.224 | 343.870 | **.027** |
| Epoch [1] × Age | 1.000 | 0.999 – 1.000 | -0.549 | 330.750 | .584 |

| | | | | | |
|---|---|---|---|---|---|
| **Epoch [2] × Age** | 0.999 | 0.999 – 1.000 | -2.376 | 341.504 | **.018** |
| Triplet type [High] × Harshness | 1.000 | 0.999 – 1.001 | 0.004 | 289781.434 | .997 |
| Triplet type [High] × Unpredictability | 1.000 | 0.999 – 1.001 | -0.785 | 289770.511 | .433 |
| Triplet type [High] × Subj cSES | 1.000 | 0.999 – 1.001 | 0.007 | 289777.796 | .994 |
| **Triplet type [High] × Age** | 1.000 | 1.000 – 1.000 | -2.147 | 289787.734 | **.032** |
| Epoch [1] × Triplet type [High] × Harshness | 1.001 | 1.000 – 1.002 | 1.655 | 289763.276 | .098 |
| Epoch [2] × Triplet type [High] × Harshness | 1.000 | 0.998 – 1.001 | -0.585 | 289803.832 | .559 |
| Epoch [1] × Triplet type [High] × Unpredictability | 1.000 | 0.999 – 1.001 | 0.447 | 289760.319 | .655 |
| **Epoch [2] × Triplet type [High] × Unpredictability** | 0.998 | 0.997 – 1.000 | -2.519 | 289809.713 | **.012** |
| Epoch [1] × Triplet type [High] × cSES | 1.000 | 0.999 – 1.001 | -0.352 | 289771.519 | .725 |
| Epoch [2] × Triplet type [High] × cSES | 1.001 | 1.000 – 1.001 | 1.376 | 289810.524 | .169 |
| Epoch [1] × Triplet type [High] × Age | 1.000 | 1.000 – 1.000 | -0.213 | 289759.386 | .831 |
| Epoch [2] × Triplet type [High] × Age | 1.000 | 1.000 – 1.000 | 0.161 | 289804.191 | .872 |
| **Random Effects** | | | | | |
| $\sigma^2$ | 0.035 | | | | |
| $\tau_{00\ \text{Participant}}$ | 0.012 | | | | |
| $\tau_{11\ \text{Participant.Epoch1}}$ | 0.001 | | | | |
| $\tau_{11\ \text{Participant.Epoch2}}$ | < 0.001 | | | | |
| $\rho_{01\ \text{Participant.Epoch1}}$ | 0.167 | | | | |
| $\rho_{01\ \text{Participant.Epoch2}}$ | 0.072 | | | | |
| ICC | 0.265 | | | | |
| $N_{\text{Participant}}$ | 325 | | | | |
| Observations | 290682 | | | | |
| Marginal $R^2$ / Conditional $R^2$ | 0.013 / 0.274 | | | | |

**Supplementary Table S1. Results of the linear mixed model on log transformed RT, learning.** Top table shows the Type 3 tests of fixed effects. Bottom table shows regression coefficients of fixed effects and summary information about the random effects. Coefficients are exponentiated to obtain the multiplicative factor for each 1 unit increase in the given independent variable. E.g., the coefficient of Age being 1.004 means that each year corresponds to a 0.4% increase in RTs. The marginal R-squared considers only the variance of the fixed effects, while the conditional R-squared takes both the fixed and random effects into account (based on Nakagawa et al., 2017). Degrees of freedom are based on Satterthwaite's approximation. Statistically significant terms are highlighted in bold. Terms in brackets indicate the level of factor that is contrasted against the reference level, which is Low triplets for the Triplet type factor and Epoch 3 for the Epoch factor.
Model equation in *lmer* syntax: log(RT) ~ Epoch*Triplet type*(Harshness + Unpredictability + Subj cSES + Age) + (Epoch | Participant)

### Table S2. Learning Accuracy.

| LRT test of effects | $\chi^2$ | *df* | *p* |
|---|---|---|---|
| **Epoch** | 73.92 | 2 | **< .001** |
| **Triplet type** | 337.38 | 1 | **< .001** |
| Harshness | 0.55 | 1 | .458 |
| Unpredictability | 0.03 | 1 | .860 |
| Subj cSES | 0.90 | 1 | .342 |
| Age | 2.24 | 1 | .135 |
| **Epoch x Triplet type** | 38.10 | 2 | **< .001** |
| Epoch x Harshness | 2.20 | 2 | .334 |
| Epoch x Unpredictability | 0.21 | 2 | .899 |
| Epoch x Subj cSES | 1.63 | 2 | .442 |
| Epoch x Age | 0.74 | 2 | .692 |
| Triplet type x Harshness | 0.01 | 1 | .907 |
| Triplet type x Unpredictability | 2.41 | 1 | .121 |
| Triplet type x Subj cSES | 3.32 | 1 | .069 |
| Triplet type x Age | 0.17 | 1 | .676 |
| Epoch x Triplet type x Harshness | 1.80 | 2 | .407 |
| Epoch x Triplet type x Unpredictability | 2.12 | 2 | .347 |
| Epoch x Triplet type x Subj cSES | 0.00 | 2 | .999 |
| Epoch x Triplet type x Age | 1.78 | 2 | .410 |

| Fixed effects | *Log-odds* | *95% CI* | *z* | *df* | *p* |
|---|---|---|---|---|---|
| **(Intercept)** | 2.459 | 2.406 – 2.512 | 90.801 | Inf | **< .001** |
| **Epoch [1]** | 0.148 | 0.115 – 0.182 | 8.695 | Inf | **< .001** |
| **Epoch [2]** | -0.043 | -0.068 – -0.019 | -3.472 | Inf | **.001** |
| **Triplet type [High]** | 0.135 | 0.120 – 0.149 | 18.697 | Inf | **< .001** |
| Harshness | -0.022 | -0.079 – 0.036 | -0.745 | Inf | .456 |
| Unpredictability | 0.005 | -0.052 – 0.062 | 0.180 | Inf | .857 |
| Subj cSES | -0.019 | -0.058 – 0.020 | -0.955 | Inf | .340 |
| Age | 0.008 | -0.002 – 0.018 | 1.497 | Inf | .134 |
| **Epoch [1] × Triplet type [High]** | -0.047 | -0.068 – -0.027 | -4.543 | Inf | **< .001** |
| Epoch [2] × Triplet type [High] | -0.012 | -0.032 – 0.008 | -1.170 | Inf | .242 |
| Epoch [1] × Harshness | -0.006 | -0.041 – 0.030 | -0.312 | Inf | .755 |
| Epoch [2] × Harshness | 0.018 | -0.007 – 0.044 | 1.416 | Inf | .157 |
| Epoch [1] × Unpredictability | 0.007 | -0.028 – 0.042 | 0.404 | Inf | .686 |
| Epoch [2] × Unpredictability | -0.005 | -0.031 – 0.020 | -0.406 | Inf | .684 |
| Epoch [1] × Subj cSES | 0.002 | -0.022 – 0.027 | 0.198 | Inf | .843 |
| Epoch [2] × Subj cSES | -0.011 | -0.028 – 0.007 | -1.194 | Inf | .233 |
| Epoch [1] × Age | -0.003 | -0.009 – 0.004 | -0.841 | Inf | .401 |
| Epoch [2] × Age | 0.001 | -0.003 – 0.006 | 0.587 | Inf | .557 |

| | | | | | |
|---|---|---|---|---|---|
| Triplet type [High] × Harshness | -0.001 | -0.016 – 0.014 | -0.117 | Inf | .907 |
| Triplet type [High] × Unpredictability | 0.012 | -0.003 – 0.027 | 1.559 | Inf | .119 |
| Triplet type [High] × Subj cSES | -0.010 | -0.020 – 0.001 | -1.829 | Inf | .067 |
| Triplet type [High] × Age | 0.001 | -0.002 – 0.003 | 0.419 | Inf | .675 |
| Epoch [1] × Triplet type [High] × Harshness | -0.013 | -0.035 – 0.009 | -1.179 | Inf | .238 |
| Epoch [2] × Triplet type [High] × Harshness | 0.001 | -0.020 – 0.022 | 0.079 | Inf | .937 |
| Epoch [1] × Triplet type [High] × Unpredictability | 0.008 | -0.015 – 0.030 | 0.673 | Inf | .501 |
| Epoch [2] × Triplet type [High] × Unpredictability | 0.008 | -0.013 – 0.029 | 0.733 | Inf | .464 |
| Epoch [1] × Triplet type [High] × Subj cSES | -0.000 | -0.016 – 0.015 | -0.053 | Inf | .958 |
| Epoch [2] × Triplet type [High] × Subj cSES | 0.000 | -0.015 – 0.015 | 0.029 | Inf | .977 |
| Epoch [1] × Triplet type [High] × Age | 0.001 | -0.003 – 0.005 | 0.704 | Inf | .481 |
| Epoch [2] × Triplet type [High] × Age | -0.003 | -0.007 – 0.001 | -1.339 | Inf | .180 |

| **Random Effects** | |
|---|---|
| $\sigma^2$ | 3.290 |
| $\tau_{00\ \text{Participant}}$ | 0.207 |
| $\tau_{11\ \text{Participant.Epoch1}}$ | 0.049 |
| $\tau_{11\ \text{Participant.Epoch2}}$ | 0.012 |
| $\rho_{01\ \text{Participant.Epoch1}}$ | 0.394 |
| $\rho_{01\ \text{Participant.Epoch2}}$ | -0.234 |
| ICC | 0.068 |
| $N_{\text{Participant}}$ | 325 |
| Observations | 316413 |
| Marginal $R^2$ / Conditional $R^2$ | 0.008 / 0.075 |

**Supplementary Table S2. Results of the binomial mixed model on accuracy, learning.** Top table shows the LRT tests of fixed effects. Bottom table shows regression coefficients of fixed effects and summary information about the random effects. Coefficients are log odds, thus

positive values indicate that the given independent variable is associated with an increase likelihood of correct responses, and negative values the opposite. P values for coefficients are from z tests, making the degrees of freedom infinite. The marginal R-squared considers only the variance of the fixed effects, while the conditional R-squared takes both the fixed and random effects into account (based on Nakagawa et al., 2017). Statistically significant terms are highlighted in bold. Terms in brackets indicate the level of factor that is contrasted against the reference level, which is Low triplets for the Triplet type factor and Epoch 3 for the Epoch factor.
Model equation in *lmer* syntax: Correct ~ Epoch*Triplet type*(Harshness + Unpredictability + Subj cSES + Age) + (Epoch | Participant)

#### Table S3. Consolidation RT.

| Type III test of effects | *F* | *df* | *p* |
|---|---|---|---|
| **Epoch** | 499.31 | 1, 331.46 | **< .001** |
| **Triplet type** | 472.63 | 1, 193144.75 | **< .001** |
| Harshness | 0.55 | 1, 319.81 | .461 |
| Unpredictability | 2.02 | 1, 319.81 | .156 |
| Subj cSES | 0.12 | 1, 319.80 | .726 |
| **Age** | 18.13 | 1, 319.72 | **< .001** |
| **Delay duration** | 10.76 | 1, 319.81 | **.001** |
| Epoch x Triplet type | 0.11 | 1, 193148.81 | .693 |
| Epoch x Harshness | 1.10 | 1, 331.20 | .296 |
| Epoch x Unpredictability | 0.00 | 1, 331.44 | .949 |
| Epoch x Subj cSES | 3.10 | 1, 331.04 | .079 |
| Epoch x Age | 0.24 | 1, 329.67 | .626 |
| Epoch x Delay duration | 1.67 | 1, 331.16 | .198 |
| Triplet type x Harshness | 0.03 | 1, 193139.23 | .874 |
| Triplet type x Unpredictability | 0.24 | 1, 193141.08 | .627 |
| Triplet type x Subj cSES | 0.45 | 1, 193147.56 | .500 |
| **Triplet type x Age** | 4.17 | 1, 193147.88 | **.041** |
| Triplet type x Delay duration | 3.01 | 1, 193140.96 | .083 |
| Epoch x Triplet type x Harshness | 0.22 | 1, 193148.29 | .638 |
| Epoch x Triplet type x Unpredictability | 0.51 | 1, 193151.72 | .476 |
| Epoch x Triplet type x Subj cSES | 0.06 | 1, 193146.63 | .813 |
| Epoch x Triplet type x Age | 1.57 | 1, 193131.92 | .211 |
| Epoch x Triplet type x Delay duration | 0.24 | 1, 193148.46 | .626 |

| Fixed effects | *exp(b)* | *95% CI* | *t* | *df* | *p* |
|---|---|---|---|---|---|
| **(Intercept)** | 364.892 | 360.734 – 369.098 | 1012.818 | 319.825 | **< .001** |
| **Epoch [3]** | 1.032 | 1.029 – 1.035 | 22.345 | 331.460 | **< .001** |
| **Triplet type [High]** | 0.990 | 0.989 – 0.991 | -21.740 | 193144.746 | **< .001** |
| Harshness | 1.005 | 0.992 – 1.017 | 0.738 | 319.810 | .461 |
| Unpredictability | 0.991 | 0.979 – 1.003 | -1.421 | 319.823 | .156 |
| Subj cSES | 1.002 | 0.993 – 1.010 | 0.351 | 319.801 | .726 |
| **Age** | 1.005 | 1.003 – 1.007 | 4.258 | 319.722 | **< .001** |
| **Delay duration** | 1.001 | 1.000 – 1.002 | 3.280 | 319.807 | **.001** |
| Epoch [3] × Triplet type [High] | 1.000 | 0.999 – 1.001 | 0.395 | 193148.807 | .693 |
| Epoch [3] × Harshness | 0.998 | 0.995 – 1.001 | -1.047 | 331.203 | .296 |
| Epoch [3] × Unpredictability | 1.000 | 0.997 – 1.003 | 0.064 | 331.443 | .949 |
| Epoch [3] × Subj cSES | 1.002 | 1.000 – 1.004 | 1.761 | 331.042 | .079 |
| Epoch [3] × Age | 1.000 | 1.000 – 1.001 | 0.488 | 329.673 | .626 |
| Epoch [3] × Delay duration | 1.000 | 1.000 – 1.000 | -1.291 | 331.164 | .198 |
| Triplet type [High] × Harshness | 1.000 | 0.999 – 1.001 | -0.159 | 193139.229 | .874 |
| Triplet type [High] × Unpredictability | 1.000 | 0.999 – 1.001 | 0.487 | 193141.075 | .627 |

| | | | | | |
|---|---|---|---|---|---|
| Triplet type [High] × Subj cSES | 1.000 | 0.999 – 1.000 | -0.674 | 193147.562 | .500 |
| **Triplet type [High] × Age** | 1.000 | 1.000 – 1.000 | -2.041 | 193147.875 | **.041** |
| Triplet type [High] × Delay duration | 1.000 | 1.000 – 1.000 | 1.734 | 193140.964 | .083 |
| Epoch [3] × Triplet type [High] × Harshness | 1.000 | 0.999 – 1.001 | -0.470 | 193148.291 | .638 |
| Epoch [3] × Triplet type [High] × Unpredictability | 1.000 | 0.999 – 1.001 | 0.713 | 193151.725 | .476 |
| Epoch [3] × Triplet type [High] × Subj cSES | 1.000 | 0.999 – 1.001 | -0.236 | 193146.627 | .813 |
| Epoch [3] × Triplet type [High] × Age | 1.000 | 1.000 – 1.000 | 1.251 | 193131.920 | .211 |
| Epoch [3] × Triplet type [High] × Delay duration | 1.000 | 1.000 – 1.000 | -0.488 | 193148.463 | .626 |

| **Random Effects** | |
|---|---|
| $\sigma^2$ | 0.030 |
| $\tau_{00\ \text{Participant}}$ | 0.010 |
| $\tau_{11\ \text{Participant.Epoch3}}$ | 0.001 |
| $\rho_{01\ \text{Participant.Epoch3}}$ | 0.112 |
| ICC | 0.262 |
| $N_{\text{Participant}}$ | 325 |
| Observations | 193772 |
| Marginal $R^2$ / Conditional $R^2$ | 0.049 / 0.298 |

**Supplementary Table S3. Results of the linear mixed model on log transformed RT, consolidation.** Top table shows the Type 3 tests of fixed effects. Bottom table shows regression coefficients of fixed effects and summary information about the random effects. Coefficients are exponentiated to obtain the multiplicative factor for each 1 unit increase in the given independent variable. E.g., the coefficient of Age being 1.005 means that each year corresponds to a 0.5% increase in RTs. The marginal R-squared considers only the variance of the fixed effects, while the conditional R-squared takes both the fixed and random effects into account (based on Nakagawa et al., 2017). Degrees of freedom are based on Satterthwaite's approximation. Statistically significant terms are highlighted in bold. Terms in brackets indicate the level of factor that is contrasted against the reference level, which is Low triplets for the Triplet type factor and Epoch 4 for the Epoch factor.

Model equation in *lmer* syntax: log(RT) ~ Epoch*Triplet type*(Harshness + Unpredictability + Subj cSES + Age + Delay duration) + (Epoch | Participant)

#### Table S4. Consolidation Accuracy.

| LRT test of effects | $\chi^2$ | *df* | *p* |
|---|---|---|---|
| **Epoch** | 32.33 | 1 | **< .001** |
| **Triplet type** | 493.07 | 1 | **< .001** |
| Harshness | 0.53 | 1 | .467 |
| Unpredictability | 0.09 | 1 | .767 |
| Subj cSES | 0.19 | 1 | .667 |
| Age | 3.71 | 1 | .054 |
| Delay duration | 2.80 | 1 | .094 |
| Epoch x Triplet type | 0.05 | 1 | .822 |
| Epoch x Harshness | 1.36 | 1 | .243 |
| Epoch x Unpredictability | 0.96 | 1 | .328 |
| Epoch x Subj cSES | 0.09 | 1 | .761 |
| Epoch x Age | 0.22 | 1 | .643 |
| Epoch x Delay duration | 0.19 | 1 | .666 |
| Triplet type x Harshness | 0.32 | 1 | .570 |
| Triplet type x Unpredictability | 1.57 | 1 | .210 |
| Triplet type x Subj cSES | 1.29 | 1 | .255 |
| Triplet type x Age | 0.88 | 1 | .349 |
| Triplet type x Delay duration | 0.14 | 1 | .705 |
| Epoch x Triplet type x Harshness | 0.34 | 1 | .558 |
| Epoch x Triplet type x Unpredictability | 1.96 | 1 | .161 |
| Epoch x Triplet type x Subj cSES | 0.37 | 1 | .541 |
| Epoch x Triplet type x Age | 0.03 | 1 | .866 |
| Epoch x Triplet type x Delay duration | 1.32 | 1 | .251 |

| Fixed effects | *Log-odds* | *95% CI* | *z* | *df* | *p* |
|---|---|---|---|---|---|
| **(Intercept)** | 2.414 | 2.365 – 2.462 | 97.681 | Inf | **<0.001** |
| **Epoch [3]** | -0.069 | -0.092 – -0.046 | -5.844 | Inf | **<0.001** |
| **Triplet type [High]** | 0.196 | 0.179 – 0.213 | 22.683 | Inf | **<0.001** |
| Harshness | -0.020 | -0.072 – 0.033 | -0.733 | Inf | 0.464 |
| Unpredictability | -0.008 | -0.059 – 0.044 | -0.292 | Inf | 0.770 |
| Subj cSES | -0.008 | -0.044 – 0.028 | -0.429 | Inf | 0.668 |
| Age | 0.009 | -0.000 – 0.019 | 1.930 | Inf | 0.054 |
| Delay duration | 0.003 | -0.000 – 0.006 | 1.679 | Inf | 0.093 |
| Epoch [3] × Triplet type [High] | -0.002 | -0.019 – 0.015 | -0.224 | Inf | 0.823 |
| Epoch [3] × Harshness | -0.015 | -0.039 – 0.010 | -1.173 | Inf | 0.241 |
| Epoch [3] × Unpredictability | 0.012 | -0.012 – 0.036 | 0.983 | Inf | 0.326 |
| Epoch [3] × Subj cSES | -0.003 | -0.019 – 0.014 | -0.305 | Inf | 0.760 |
| Epoch [3] × Age | 0.001 | -0.003 – 0.006 | 0.465 | Inf | 0.642 |
| Epoch [3] × Delay duration | -0.000 | -0.002 – 0.001 | -0.433 | Inf | 0.665 |
| Triplet type [High] × Harshness | 0.005 | -0.013 – 0.023 | 0.569 | Inf | 0.569 |
| Triplet type [High] × Unpredictability | 0.011 | -0.006 – 0.029 | 1.259 | Inf | 0.208 |

| | | | | | |
|---|---|---|---|---|---|
| Triplet type [High] × Subj cSES | -0.007 | -0.020 – 0.005 | -1.142 | Inf | 0.253 |
| Triplet type [High] × Age | 0.002 | -0.002 – 0.005 | 0.942 | Inf | 0.346 |
| Triplet type [High] × Delay duration | -0.000 | -0.001 – 0.001 | -0.381 | Inf | 0.703 |
| Epoch [3] × Triplet type [High] × Harshness | 0.005 | -0.013 – 0.023 | 0.590 | Inf | 0.556 |
| Epoch [3] × Triplet type [High] × Unpredictability | -0.013 | -0.031 – 0.005 | -1.405 | Inf | 0.160 |
| Epoch [3] × Triplet type [High] × Subj cSES | -0.004 | -0.016 – 0.009 | -0.614 | Inf | 0.539 |
| Epoch [3] × Triplet type [High] × Age | 0.000 | -0.003 – 0.004 | 0.170 | Inf | 0.865 |
| Epoch [3] × Triplet type [High] × Delay duration | 0.001 | -0.000 – 0.002 | 1.152 | Inf | 0.249 |

| **Random Effects** | |
|---|---|
| $\sigma^2$ | 3.290 |
| $\tau_{00\ \text{Participant}}$ | 0.161 |
| $\tau_{11\ \text{Participant.Epoch3}}$ | 0.017 |
| $\rho_{01\ \text{Participant.Epoch3}}$ | 0.058 |
| ICC | 0.051 |
| $N_{\text{Participant}}$ | 325 |
| Observations | 211110 |
| Marginal $R^2$ / Conditional $R^2$ | 0.012 / 0.062 |

**Supplementary Table S4. Results of the binomial mixed model on accuracy, consolidation.** Top table shows the LRT tests of fixed effects. Bottom table shows regression coefficients of fixed effects and summary information about the random effects. Coefficients are log odds, thus positive values indicate that the given independent variable is associated with an increase likelihood of correct responses, and negative values the opposite. P values for coefficients are from z tests, making the degrees of freedom infinite. The marginal R-squared considers only the variance of the fixed effects, while the conditional R-squared takes both the fixed and random effects into account (based on Nakagawa et al., 2017). Statistically significant terms are highlighted in bold. Terms in brackets indicate the level of factor that is contrasted against the reference level, which is Low triplets for the Triplet type factor and Epoch 4 for the Epoch factor.

Model equation in *lmer* syntax: Correct ~ Epoch*Triplet type*(Harshness + Unpredictability + Subj cSES + Age + Delay duration) + (Epoch | Participant)

### Table S5. Rewiring Old knowledge RT.

| Type III test of effects | *F* | *df* | *p* |
|---|---|---|---|
| **Epoch** | 99.41 | 2, 365.11 | **< .001** |
| **Triplet type** | 484.72 | 1, 195586.83 | **< .001** |
| Harshness | 1.11 | 1, 321.42 | .293 |
| Unpredictability | 1.67 | 1, 321.44 | .197 |
| Subj cSES | 0.03 | 1, 321.39 | .869 |
| **Age** | 16.98 | 1, 321.35 | **< .001** |
| Epoch x Triplet type | 2.01 | 2, 195590.43 | .134 |
| Epoch x Harshness | 0.03 | 2, 366.25 | .974 |
| Epoch x Unpredictability | 1.47 | 2, 366.84 | .232 |
| Epoch x Subj cSES | 0.70 | 2, 364.65 | .495 |
| Epoch x Age | 0.37 | 2, 363.44 | .691 |
| Triplet type x Harshness | 0.62 | 1, 195584.13 | .432 |
| Triplet type x Unpredictability | 0.19 | 1, 195592.27 | .664 |
| Triplet type x Subj cSES | 0.66 | 1, 195560.77 | .417 |
| **Triplet type x Age** | 5.39 | 1, 195578.64 | **.020** |
| Epoch x Triplet type x Harshness | 0.73 | 2, 195577.93 | .480 |
| Epoch x Triplet type x Unpredictability | 0.42 | 2, 195592.19 | .656 |
| Epoch x Triplet type x Subj cSES | 1.65 | 2, 195591.75 | .193 |
| Epoch x Triplet type x Age | 1.43 | 2, 195587.45 | .240 |

| Fixed effects | *exp(b)* | *95% CI* | *t* | *df* | *p* |
|---|---|---|---|---|---|
| **(Intercept)** | 360.326 | 356.328 – 364.368 | 1038.248 | 321.395 | **< .001** |
| **Epoch [4]** | 0.980 | 0.977 – 0.983 | -14.095 | 375.299 | **< .001** |
| **Epoch [5]** | 1.008 | 1.005 – 1.010 | 6.476 | 348.336 | **< .001** |
| **Triplet type [H L]** | 0.989 | 0.988 – 0.990 | -22.016 | 195586.830 | **< .001** |
| Harshness | 1.007 | 0.994 – 1.019 | 1.054 | 321.420 | .293 |
| Unpredictability | 0.992 | 0.980 – 1.004 | -1.291 | 321.435 | .197 |
| Subj cSES | 0.999 | 0.991 – 1.008 | -0.165 | 321.388 | .869 |
| **Age** | 1.005 | 1.002 – 1.007 | 4.121 | 321.354 | **< .001** |
| Epoch [4] × Triplet type [H L] | 1.000 | 0.999 – 1.002 | 0.322 | 195558.741 | .748 |
| Epoch [5] × Triplet type [H L] | 1.001 | 1.000 – 1.003 | 1.439 | 195362.426 | .150 |
| Epoch [4] × Harshness | 1.000 | 0.997 – 1.003 | -0.048 | 375.125 | .962 |
| Epoch [5] × Harshness | 1.000 | 0.998 – 1.003 | 0.223 | 349.805 | .824 |
| Epoch [4] × Unpredictability | 0.999 | 0.996 – 1.002 | -0.389 | 375.956 | .698 |
| Epoch [5] × Unpredictability | 1.002 | 1.000 – 1.005 | 1.671 | 350.401 | .096 |
| Epoch [4] × Subj cSES | 1.000 | 0.998 – 1.002 | -0.310 | 374.781 | .757 |
| Epoch [5] × Subj cSES | 0.999 | 0.997 – 1.001 | -0.899 | 347.281 | .369 |
| Epoch [4] × Age | 1.000 | 0.999 – 1.000 | -0.228 | 373.742 | .819 |
| Epoch [5] × Age | 1.000 | 0.999 – 1.000 | -0.648 | 346.283 | .518 |

| | | | | | |
|---|---|---|---|---|---|
| Triplet type [H L] × Harshness | 1.000 | 0.999 – 1.002 | 0.786 | 195584.135 | .432 |
| Triplet type [H L] × Unpredictability | 1.000 | 0.999 – 1.001 | -0.434 | 195592.266 | .664 |
| Triplet type [H L] × Subj cSES | 1.000 | 1.000 – 1.001 | 0.811 | 195560.768 | .417 |
| **Triplet type [H L] × Age** | 1.000 | 1.000 – 1.000 | -2.321 | 195578.639 | **.020** |
| Epoch [4] × Triplet type [H L] × Harshness | 0.999 | 0.998 – 1.001 | -1.149 | 195536.236 | .250 |
| Epoch [5] × Triplet type [H L] × Harshness | 1.001 | 0.999 – 1.003 | 0.970 | 195487.320 | .332 |
| Epoch [4] × Triplet type [H L] × Unpredictability | 1.000 | 0.999 – 1.002 | 0.652 | 195555.427 | .514 |
| Epoch [5] × Triplet type [H L] × Unpredictability | 0.999 | 0.998 – 1.001 | -0.902 | 195477.528 | .367 |
| Epoch [4] × Triplet type [H L] × Subj cSES | 1.000 | 0.999 – 1.002 | 0.965 | 195579.345 | .335 |
| Epoch [4] × Triplet type [H L] × Subj cSES | 0.999 | 0.998 – 1.000 | -1.813 | 195221.603 | .070 |
| Epoch [4] × Triplet type [H L] × Age | 1.000 | 1.000 – 1.000 | -0.583 | 195551.976 | .560 |
| Epoch [5] × Triplet type [H L] × Age | 1.000 | 1.000 – 1.001 | 1.634 | 195323.957 | .102 |

| **Random Effects** | |
|---|---|
| $\sigma^2$ | 0.028 |
| $\tau_{00\ \text{Participant}}$ | 0.010 |
| $\tau_{11\ \text{Participant.Epoch4}}$ | < 0.001 |
| $\tau_{11\ \text{Participant.Epoch5}}$ | < 0.001 |
| $\rho_{01\ \text{Participant.Epoch4}}$ | 0.029 |
| $\rho_{01\ \text{Participant.Epoch5}}$ | -0.060 |
| ICC | 0.266 |
| $N_{\text{Participant}}$ | 325 |
| Observations | 196226 |
| Marginal $R^2$ / Conditional $R^2$ | 0.024 / 0.283 |

**Supplementary Table S5. Results of the linear mixed model on log transformed RT, rewiring old knowledge.** Top table shows the Type 3 tests of fixed effects. Bottom table shows regression coefficients of fixed effects and summary information about the random effects.

Coefficients are exponentiated to obtain the multiplicative factor for each 1 unit increase in the given independent variable. E.g., the coefficient of Age being 1.005 means that each year corresponds to a 0.5% increase in RTs. The marginal R-squared considers only the variance of the fixed effects, while the conditional R-squared takes both the fixed and random effects into account (based on Nakagawa et al., 2017). Degrees of freedom are based on Satterthwaite's approximation. Statistically significant terms are highlighted in bold. Terms in brackets indicate the level of factor that is contrasted against the reference level, which is L L triplets for the Triplet type factor and Epoch 6 for the Epoch factor.
Model equation in *lmer* syntax: log(RT) ~ Epoch*Triplet type*(Harshness + Unpredictability + Subj cSES + Age) + (Epoch | Participant)

### Table S6. Rewiring Old knowledge Accuracy.

| LRT test of effects | $\chi^2$ | *df* | *p* |
|---|---|---|---|
| **Epoch** | 9.34 | 2 | **.009** |
| **Triplet type** | 217.58 | 1 | **< .001** |
| Harshness | 0.04 | 1 | .835 |
| Unpredictability | 0.55 | 1 | .460 |
| Subj cSES | 2.88 | 1 | .090 |
| Age | 3.04 | 1 | .081 |
| **Epoch x Triplet type** | 22.27 | 2 | **< .001** |
| Epoch x Harshness | 0.47 | 2 | .791 |
| Epoch x Unpredictability | 0.97 | 2 | .616 |
| **Epoch x Subj cSES** | 7.11 | 2 | **.029** |
| Epoch x Age | 0.18 | 2 | .914 |
| Triplet type x Harshness | 0.09 | 1 | .767 |
| Triplet type x Unpredictability | 3.45 | 1 | .063 |
| Triplet type x Subj cSES | 0.86 | 1 | .354 |
| Triplet type x Age | 0.45 | 1 | .503 |
| Epoch x Triplet type x Harshness | 0.08 | 2 | .959 |
| Epoch x Triplet type x Unpredictability | 1.27 | 2 | .529 |
| Epoch x Triplet type x Subj cSES | 1.56 | 2 | .458 |
| Epoch x Triplet type x Age | 0.74 | 2 | .692 |

| Fixed effects | *Log-odds* | *95% CI* | *z* | *df* | *p* |
|---|---|---|---|---|---|
| **(Intercept)** | 2.426 | 2.379 – 2.472 | 101.850 | Inf | **< .001** |
| **Epoch [4]** | 0.053 | 0.019 – 0.086 | 3.068 | Inf | **.002** |
| Epoch [5] | -0.031 | -0.066 – 0.004 | -1.722 | Inf | .085 |
| **Triplet type [H L]** | 0.154 | 0.134 – 0.174 | 14.968 | Inf | **< .001** |
| Harshness | 0.005 | -0.045 – 0.056 | 0.212 | Inf | .832 |
| Unpredictability | -0.019 | -0.068 – 0.031 | -0.742 | Inf | .458 |
| Subj cSES | -0.030 | -0.064 – 0.004 | -1.703 | Inf | .089 |
| Age | 0.008 | -0.001 – 0.017 | 1.748 | Inf | .080 |
| **Epoch [4] × Triplet type [H L]** | 0.044 | 0.017 – 0.072 | 3.218 | Inf | **.001** |
| **Epoch [5] × Triplet type [H L]** | -0.075 | -0.107 – -0.044 | -4.720 | Inf | **< .001** |
| Epoch [4] × Harshness | -0.010 | -0.045 – 0.025 | -0.562 | Inf | .574 |
| Epoch [5] × Harshness | -0.000 | -0.037 – 0.036 | -0.013 | Inf | .989 |
| Epoch [4] × Unpredictability | -0.005 | -0.039 – 0.030 | -0.275 | Inf | .784 |
| Epoch [5] × Unpredictability | 0.017 | -0.019 – 0.053 | 0.939 | Inf | .348 |
| Epoch [4] × Subj cSES | 0.008 | -0.016 – 0.032 | 0.626 | Inf | .531 |
| Epoch [5] × Subj cSES | 0.023 | -0.002 – 0.048 | 1.800 | Inf | .072 |
| Epoch [4] × Age | 0.001 | -0.005 – 0.008 | 0.327 | Inf | .743 |
| Epoch [5] × Age | -0.001 | -0.008 – 0.005 | -0.410 | Inf | .682 |

| | | | | | |
|---|---|---|---|---|---|
| Triplet type [H L] × Harshness | -0.003 | -0.025 – 0.018 | -0.296 | Inf | .767 |
| Triplet type [H L] × Unpredictability | 0.020 | -0.001 – 0.041 | 1.866 | Inf | .062 |
| Triplet type [H L] × Subj cSES | 0.007 | -0.008 – 0.022 | 0.931 | Inf | .352 |
| Triplet type [H L] × Age | -0.001 | -0.005 – 0.003 | -0.673 | Inf | .501 |
| Epoch [4] × Triplet type [H L] × Harshness | -0.001 | -0.029 – 0.028 | -0.034 | Inf | .973 |
| Epoch [5] × Triplet type [H L] × Harshness | 0.004 | -0.029 – 0.038 | 0.253 | Inf | .800 |
| Epoch [4] × Triplet type [H L] × Unpredictability | 0.009 | -0.019 – 0.037 | 0.617 | Inf | .537 |
| Epoch [5] × Triplet type [H L] × Unpredictability | -0.019 | -0.052 – 0.014 | -1.135 | Inf | .256 |
| Epoch [4] × Triplet type [H L] × Subj cSES | 0.010 | -0.009 – 0.030 | 1.037 | Inf | .300 |
| Epoch [4] × Triplet type [H L] × Subj cSES | -0.014 | -0.037 – 0.009 | -1.188 | Inf | .235 |
| Epoch [4] × Triplet type [H L] × Age | 0.000 | -0.005 – 0.005 | 0.039 | Inf | .969 |
| Epoch [5] × Triplet type [H L] × Age | 0.002 | -0.004 – 0.008 | 0.676 | Inf | .499 |
| **Random Effects** | | | | | |
| $\sigma^2$ | 3.290 | | | | |
| $\tau_{00}$ Participant | 0.137 | | | | |
| $\tau_{11}$ Participant.Epoch4 | 0.026 | | | | |
| $\tau_{11}$ Participant.Epoch5 | 0.012 | | | | |
| $\rho_{01}$ Participant.Epoch4 | 0.084 | | | | |
| $\rho_{01}$ Participant.Epoch5 | -0.145 | | | | |
| ICC | 0.047 | | | | |
| N Participant | 325 | | | | |
| Observations | 212870 | | | | |
| Marginal $R^2$ / Conditional $R^2$ | 0.008 / 0.055 | | | | |

**Supplementary Table S6. Results of the binomial mixed model on accuracy, rewiring old knowledge.** Top table shows the LRT tests of fixed effects. Bottom table shows regression coefficients of fixed effects and summary information about the random effects. Coefficients are log odds, thus positive values indicate that the given independent variable is associated with an increase likelihood of correct responses, and negative values the opposite. P values for coefficients are from z tests, making the degrees of freedom infinite. The marginal R-squared considers only the variance of the fixed effects, while the conditional R-squared takes both the fixed and random effects into account (based on Nakagawa et al., 2017). Statistically significant terms are highlighted in bold. Terms in brackets indicate the level of factor that is contrasted against the reference level, which is L L triplets for the Triplet type factor and Epoch 6 for the Epoch factor.

Model equation in *lmer* syntax: Correct ~ Epoch*Triplet type*(Harshness + Unpredictability + Subj cSES + Age) + (Epoch | Participant)

#### Table S7. Rewiring New knowledge RT.

| Type III test of effects | *F* | *df* | *p* |
|---|---|---|---|
| **Epoch** | 87.77 | 2, 354.19 | **< .001** |
| **Triplet type** | 39.15 | 1, 137396.15 | **< .001** |
| Harshness | 0.90 | 1, 320.77 | .343 |
| Unpredictability | 1.55 | 1, 320.84 | .213 |
| Subj cSES | 0.00 | 1, 320.74 | .954 |
| **Age** | 17.49 | 1, 320.71 | **< .001** |
| Epoch x Triplet type | 2.19 | 2, 137324.51 | .111 |
| Epoch x Harshness | 0.05 | 2, 355.10 | .952 |
| Epoch x Unpredictability | 2.70 | 2, 357.50 | .069 |
| Epoch x Subj cSES | 0.12 | 2, 353.19 | .889 |
| Epoch x Age | 1.33 | 2, 352.61 | .267 |
| Triplet type x Harshness | 0.17 | 1, 137370.68 | .676 |
| Triplet type x Unpredictability | 0.00 | 1, 137414.88 | .945 |
| **Triplet type x Subj cSES** | 8.43 | 1, 137417.54 | **.004** |
| Triplet type x Age | 3.30 | 1, 137425.03 | .069 |
| Epoch x Triplet type x Harshness | 0.04 | 2, 137304.38 | .957 |
| Epoch x Triplet type x Unpredictability | 0.11 | 2, 137324.02 | .893 |
| Epoch x Triplet type x Subj cSES | 0.51 | 2, 137333.07 | .603 |
| Epoch x Triplet type x Age | 1.06 | 2, 137340.33 | .348 |

| Fixed effects | *exp(b)* | *95% CI* | *t* | *df* | *p* |
|---|---|---|---|---|---|
| **(Intercept)** | 363.193 | 359.194 – 367.237 | 1047.457 | 320.756 | **< .001** |
| **Epoch [4]** | 0.981 | 0.978 – 0.984 | -13.234 | 326.685 | **< .001** |
| **Epoch [5]** | 1.007 | 1.005 – 1.009 | 6.143 | 399.694 | **< .001** |
| **Triplet type [L H]** | 0.996 | 0.995 – 0.998 | -6.257 | 137396.150 | **< .001** |
| Harshness | 1.006 | 0.994 – 1.018 | 0.950 | 320.774 | .343 |
| Unpredictability | 0.993 | 0.981 – 1.004 | -1.247 | 320.837 | .213 |
| Subj cSES | 1.000 | 0.992 – 1.008 | 0.058 | 320.739 | .954 |
| **Age** | 1.005 | 1.002 – 1.007 | 4.182 | 320.712 | **< .001** |
| Epoch [4] × Triplet type [L H] | 1.002 | 1.000 – 1.003 | 1.823 | 137368.296 | .068 |
| Epoch [5] × Triplet type [L H] | 1.000 | 0.999 – 1.002 | 0.133 | 137288.011 | .894 |
| Epoch [4] × Harshness | 1.000 | 0.997 – 1.004 | 0.299 | 325.782 | .765 |
| Epoch [5] × Harshness | 1.000 | 0.997 – 1.002 | -0.212 | 402.019 | .832 |
| Epoch [4] × Unpredictability | 0.999 | 0.996 – 1.002 | -0.575 | 328.819 | .565 |
| **Epoch [5] × Unpredictability** | 1.003 | 1.000 – 1.005 | 2.283 | 405.140 | **.023** |
| Epoch [4] × Subj cSES | 1.000 | 0.998 – 1.002 | -0.378 | 325.912 | .706 |
| Epoch [5] × Subj cSES | 1.000 | 0.998 – 1.001 | -0.117 | 397.093 | .907 |
| Epoch [4] × Age | 1.000 | 1.000 – 1.001 | 0.435 | 323.653 | .664 |
| Epoch [5] × Age | 1.000 | 0.999 – 1.000 | -1.607 | 398.959 | .109 |

| | | | | | |
|---|---|---|---|---|---|
| Triplet type [L H] × Harshness | 1.000 | 0.999 – 1.001 | -0.418 | 137370.675 | .676 |
| Triplet type [L H] × Unpredictability | 1.000 | 0.999 – 1.001 | 0.068 | 137414.875 | .945 |
| **Triplet type [L H] × Subj cSES** | 1.001 | 1.000 – 1.002 | 2.903 | 137417.538 | **.004** |
| Triplet type [L H] × Age | 1.000 | 1.000 – 1.000 | -1.817 | 137425.029 | .069 |
| Epoch [4] × Triplet type [L H] × Harshness | 1.000 | 0.998 – 1.002 | -0.269 | 137343.893 | .788 |
| Epoch [5] × Triplet type [L H] × Harshness | 1.000 | 0.999 – 1.002 | 0.233 | 137277.668 | .816 |
| Epoch [4] × Triplet type [L H] × Unpredictability | 1.000 | 0.999 – 1.002 | 0.400 | 137373.168 | .689 |
| Epoch [5] × Triplet type [L H] × Unpredictability | 1.000 | 0.998 – 1.001 | -0.406 | 137281.999 | .684 |
| Epoch [4] × Triplet type [L H] × Subj cSES | 1.001 | 0.999 – 1.002 | 0.954 | 137376.115 | .340 |
| Epoch [4] × Triplet type [L H] × Subj cSES | 1.000 | 0.999 – 1.001 | -0.700 | 137316.495 | .484 |
| Epoch [4] × Triplet type [L H] × Age | 1.000 | 1.000 – 1.000 | 0.894 | 137419.190 | .371 |
| Epoch [5] × Triplet type [L H] × Age | 1.000 | 1.000 – 1.000 | 0.649 | 137319.071 | .517 |
| **Random Effects** | | | | | |
| $\sigma^2$ | 0.028 | | | | |
| $\tau_{00}$ Participant | 0.010 | | | | |
| $\tau_{11}$ Participant.Epoch4 | < 0.001 | | | | |
| $\tau_{11}$ Participant.Epoch5 | < 0.001 | | | | |
| $\rho_{01}$ Participant.Epoch4 | 0.052 | | | | |
| $\rho_{01}$ Participant.Epoch5 | 0.075 | | | | |
| ICC | 0.259 | | | | |
| N Participant | 325 | | | | |
| Observations | 137887 | | | | |
| Marginal $R^2$ / Conditional $R^2$ | 0.017 / 0.272 | | | | |

**Supplementary Table S7. Results of the linear mixed model on log transformed RT, rewiring new knowledge.** Top table shows the Type 3 tests of fixed effects. Bottom table shows regression coefficients of fixed effects and summary information about the random

effects. Coefficients are exponentiated to obtain the multiplicative factor for each 1 unit increase in the given independent variable. E.g., the coefficient of Age being 1.005 means that each year corresponds to a 0.5% increase in RTs. The marginal R-squared considers only the variance of the fixed effects, while the conditional R-squared takes both the fixed and random effects into account (based on Nakagawa et al., 2017). Degrees of freedom are based on Satterthwaite's approximation. Statistically significant terms are highlighted in bold. Terms in brackets indicate the level of factor that is contrasted against the reference level, which is L L triplets for the Triplet type factor and Epoch 6 for the Epoch factor.
Model equation in *lmer* syntax: log(RT) ~ Epoch*Triplet type*(Harshness + Unpredictability + Subj cSES + Age) + (Epoch | Participant)

### Table S8. Rewiring New knowledge Accuracy.

| LRT test of effects | $\chi^2$ | *df* | *p* |
|---|---|---|---|
| **Epoch** | 43.67 | 2 | **< .001** |
| Triplet type | 3.09 | 1 | .079 |
| Harshness | 0.05 | 1 | .822 |
| Unpredictability | 0.45 | 1 | .504 |
| Subj cSES | 0.36 | 1 | .547 |
| Age | 1.16 | 1 | .282 |
| **Epoch x Triplet type** | 14.09 | 2 | **< .001** |
| Epoch x Harshness | 0.51 | 2 | .775 |
| Epoch x Unpredictability | 3.46 | 2 | .177 |
| Epoch x Subj cSES | 0.90 | 2 | .639 |
| Epoch x Age | 0.39 | 2 | .821 |
| Triplet type x Harshness | 1.70 | 1 | .192 |
| **Triplet type x Unpredictability** | 4.32 | 1 | **.038** |
| **Triplet type x Subj cSES** | 11.60 | 1 | **< .001** |
| **Triplet type x Age** | 4.21 | 1 | **.040** |
| Epoch x Triplet type x Harshness | 0.25 | 2 | .885 |
| Epoch x Triplet type x Unpredictability | 2.44 | 2 | .295 |
| **Epoch x Triplet type x Subj cSES** | 12.03 | 2 | **.002** |
| Epoch x Triplet type x Age | 0.23 | 2 | .892 |

| Fixed effects | *Log-odds* | *95% CI* | *z* | *df* | *p* |
|---|---|---|---|---|---|
| **(Intercept)** | 2.290 | 2.244 – 2.336 | 97.284 | Inf | **< .001** |
| Epoch [4] | -0.005 | -0.044 – 0.033 | -0.281 | Inf | .779 |
| **Epoch [5]** | 0.100 | 0.068 – 0.133 | 6.049 | Inf | **< .001** |
| Triplet type [L H] | 0.019 | -0.002 – 0.040 | 1.768 | Inf | .077 |
| Harshness | -0.006 | -0.055 – 0.044 | -0.225 | Inf | .822 |
| Unpredictability | -0.017 | -0.066 – 0.032 | -0.670 | Inf | .503 |
| Subj cSES | -0.010 | -0.044 – 0.023 | -0.604 | Inf | .546 |
| Age | 0.005 | -0.004 – 0.014 | 1.078 | Inf | .281 |
| Epoch [4] × Triplet type [L H] | -0.019 | -0.050 – 0.012 | -1.213 | Inf | .225 |
| **Epoch [5] × Triplet type [L H]** | 0.053 | 0.025 – 0.081 | 3.742 | Inf | **< .001** |
| Epoch [4] × Harshness | -0.003 | -0.042 – 0.037 | -0.127 | Inf | .899 |
| Epoch [5] × Harshness | -0.009 | -0.043 – 0.025 | -0.540 | Inf | .589 |
| Epoch [4] × Unpredictability | -0.033 | -0.073 – 0.006 | -1.673 | Inf | .094 |
| Epoch [5] × Unpredictability | 0.027 | -0.006 – 0.061 | 1.588 | Inf | .112 |
| Epoch [4] × Subj cSES | 0.011 | -0.017 – 0.038 | 0.772 | Inf | .440 |
| Epoch [5] × Subj cSES | 0.001 | -0.022 – 0.024 | 0.072 | Inf | .943 |
| Epoch [4] × Age | 0.000 | -0.007 – 0.007 | 0.034 | Inf | .973 |
| Epoch [5] × Age | -0.002 | -0.008 – 0.004 | -0.559 | Inf | .576 |

| | | | | | |
|---|---|---|---|---|---|
| Triplet type [L H] × Harshness | -0.015 | -0.038 – 0.007 | -1.310 | Inf | .190 |
| Triplet type [L H] × Unpredictability | 0.024 | 0.001 – 0.046 | 2.087 | Inf | **.037** |
| **Triplet type [L H] × Subj cSES** | 0.027 | 0.012 – 0.042 | 3.420 | Inf | **.001** |
| **Triplet type [L H] × Age** | -0.004 | -0.008 – -0.000 | -2.055 | Inf | **.040** |
| Epoch [4] × Triplet type [L H] × Harshness | 0.008 | -0.025 – 0.041 | 0.478 | Inf | .633 |
| Epoch [5] × Triplet type [L H] × Harshness | -0.005 | -0.035 – 0.025 | -0.338 | Inf | .735 |
| Epoch [4] × Triplet type [L H] × Unpredictability | -0.016 | -0.049 – 0.016 | -0.976 | Inf | .329 |
| Epoch [5] × Triplet type [L H] × Unpredictability | -0.010 | -0.040 – 0.020 | -0.649 | Inf | .516 |
| Epoch [4] × Triplet type [L H] × Subj cSES | 0.014 | -0.008 – 0.037 | 1.237 | Inf | .216 |
| **Epoch [4] × Triplet type [L H] × Subj cSES** | -0.036 | -0.056 – -0.016 | -3.456 | Inf | **.001** |
| Epoch [4] × Triplet type [L H] × Age | -0.000 | -0.006 – 0.006 | -0.087 | Inf | .931 |
| Epoch [5] × Triplet type [L H] × Age | 0.001 | -0.004 – 0.007 | 0.462 | Inf | .644 |

| **Random Effects** | |
|---|---|
| $\sigma^2$ | 3.290 |
| $\tau_{00\ \mathrm{Participant}}$ | 0.130 |
| $\tau_{11\ \mathrm{Participant.Epoch4}}$ | 0.031 |
| $\tau_{11\ \mathrm{Participant.Epoch5}}$ | 0.017 |
| $\rho_{01\ \mathrm{Participant.Epoch4}}$ | 0.129 |
| $\rho_{01\ \mathrm{Participant.Epoch5}}$ | -0.047 |
| ICC | 0.043 |
| $N_{\mathrm{Participant}}$ | 325 |
| Observations | 151710 |
| Marginal $R^2$ / Conditional $R^2$ | 0.005 / 0.047 |

**Supplementary Table S8. Results of the binomial mixed model on accuracy, rewiring new knowledge.** Top table shows the LRT tests of fixed effects. Bottom table shows regression coefficients of fixed effects and summary information about the random effects. Coefficients

are log odds, thus positive values indicate that the given independent variable is associated with an increase likelihood of correct responses, and negative values the opposite. P values for coefficients are from z tests, making the degrees of freedom infinite. The marginal R-squared considers only the variance of the fixed effects, while the conditional R-squared takes both the fixed and random effects into account (based on Nakagawa et al., 2017). Statistically significant terms are highlighted in bold. Terms in brackets indicate the level of factor that is contrasted against the reference level, which is L L triplets for the Triplet type factor and Epoch 6 for the Epoch factor.
Model equation in *lmer* syntax: Correct ~ Epoch*Triplet type*(Harshness + Unpredictability + Subj cSES + Age) + (Epoch | Participant)

**Table S9. Sensitivity model Learning RT.**

| Type III test of effects | *F* | *df* | *p* |
|---|---|---|---|
| **Epoch** | 54.01 | 2, 336.34 | **< .001** |
| **Triplet type** | 288.93 | 1, 289784.48 | **< .001** |
| Harshness | 0.33 | 1, 319.59 | .568 |
| Unpredictability | 0.92 | 1, 319.61 | .337 |
| Subj cSES | 0.05 | 1, 319.58 | .830 |
| **Age** | 9.84 | 1, 319.54 | **.002** |
| Parental education | 0.05 | 1, 319.60 | .824 |
| **Epoch x Triplet type** | 18.97 | 2, 289786.71 | **< .001** |
| Epoch x Harshness | 0.17 | 2, 336.69 | .847 |
| Epoch x Unpredictability | 0.74 | 2, 337.52 | .479 |
| **Epoch x Subj cSES** | 3.15 | 2, 336.10 | **.044** |
| **Epoch x Age** | 4.24 | 2, 334.43 | **.015** |
| Epoch x Parental education | 1.86 | 2, 336.99 | .158 |
| Triplet type x Harshness | 0.00 | 1, 289778.95 | .976 |
| Triplet type x Unpredictability | 0.91 | 1, 289767.28 | .341 |
| Triplet type x Subj cSES | 0.01 | 1, 289775.54 | .930 |
| **Triplet type x Age** | 4.90 | 1, 289785.06 | **.027** |
| Triplet type x Parental education | 1.48 | 1, 289774.66 | .224 |
| Epoch x Triplet type x Harshness | 1.48 | 2, 289780.75 | .227 |
| **Epoch x Triplet type x Unpredictability** | 3.33 | 2, 289780.14 | **.036** |
| Epoch x Triplet type x Subj cSES | 1.08 | 2, 289786.34 | .341 |
| Epoch x Triplet type x Age | 0.02 | 2, 289780.00 | .985 |
| Epoch x Triplet type x Parental education | 1.46 | 1, 289777.89 | .231 |

| Fixed effects | *exp(b)* | *95% CI* | *t* | *df* | *p* |
|---|---|---|---|---|---|
| **(Intercept)** | 386.767 | 381.991 – 391.603 | 943.308 | 319.585 | **< .001** |
| **Epoch [1]** | 1.011 | 1.008 – 1.015 | 6.394 | 330.489 | **< .001** |
| **Epoch [2]** | 1.006 | 1.003 – 1.008 | 4.722 | 343.120 | **< .001** |
| **Triplet type [High]** | 0.993 | 0.992 – 0.994 | -16.998 | 289784.476 | **< .001** |
| Harshness | 1.004 | 0.991 – 1.017 | 0.572 | 319.591 | .568 |
| Unpredictability | 0.993 | 0.980 – 1.007 | -0.961 | 319.608 | .337 |
| Subj cSES | 1.001 | 0.992 – 1.010 | 0.215 | 319.576 | .830 |
| **Age** | 1.004 | 1.001 – 1.006 | 3.137 | 319.538 | **.002** |
| Parental education | 1.001 | 0.993 – 1.009 | 0.222 | 319.598 | .824 |
| **Epoch [1] × Triplet type [High]** | 1.003 | 1.002 – 1.004 | 5.574 | 289769.230 | **< .001** |
| Epoch [2] × Triplet type [High] | 1.000 | 0.999 – 1.001 | -0.483 | 289805.041 | .629 |
| Epoch [1] × Harshness | 1.001 | 0.997 – 1.004 | 0.276 | 330.829 | .783 |
| Epoch [2] × Harshness | 1.000 | 0.998 – 1.003 | 0.342 | 343.791 | .732 |
| Epoch [1] × Unpredictability | 1.002 | 0.999 – 1.006 | 1.205 | 330.970 | .229 |
| Epoch [2] × Unpredictability | 1.000 | 0.997 – 1.002 | -0.364 | 345.563 | .716 |
| Epoch [1] × Subj cSES | 1.000 | 0.998 – 1.003 | 0.022 | 330.230 | .983 |

| | | | | | |
|---|---|---|---|---|---|
| **Epoch [2] × Subj cSES** | 0.998 | 0.996 – 1.000 | -2.285 | 343.012 | **.023** |
| Epoch [1] × Age | 1.000 | 0.999 – 1.001 | -0.433 | 329.799 | .666 |
| **Epoch [2] × Age** | 0.999 | 0.999 – 1.000 | -2.426 | 340.373 | **.016** |
| Epoch [1] × Parental education | 1.002 | 1.000 – 1.004 | 1.924 | 330.773 | .055 |
| Epoch [2] × Parental education | 0.999 | 0.998 – 1.001 | -0.910 | 344.530 | .363 |
| Triplet type [High] × Harshness | 1.000 | 0.999 – 1.001 | -0.030 | 289778.949 | .976 |
| Triplet type [High] × Unpredictability | 1.000 | 0.999 – 1.000 | -0.952 | 289767.284 | .341 |
| Triplet type [High] × Subj cSES | 1.000 | 0.999 – 1.001 | -0.088 | 289775.541 | .930 |
| **Triplet type [High] × Age** | 1.000 | 1.000 – 1.000 | -2.214 | 289785.059 | **.027** |
| Triplet type [High] × Parental education | 1.000 | 0.999 – 1.000 | -1.215 | 289774.657 | .224 |
| Epoch [1] × Triplet type [High] × Harshness | 1.001 | 1.000 – 1.002 | 1.695 | 289760.121 | .090 |
| Epoch [2] × Triplet type [High] × Harshness | 1.000 | 0.998 – 1.001 | -0.581 | 289800.459 | .561 |
| Epoch [1] × Triplet type [High] × Unpredictability | 1.000 | 0.999 – 1.002 | 0.654 | 289758.776 | .513 |
| **Epoch [2] × Triplet type [High] × Unpredictability** | 0.998 | 0.997 – 1.000 | -2.489 | 289805.579 | **.013** |
| Epoch [1] × Triplet type [High] × cSES | 1.000 | 0.999 – 1.001 | -0.239 | 289768.566 | .811 |
| Epoch [2] × Triplet type [High] × cSES | 1.001 | 1.000 – 1.001 | 1.375 | 289807.744 | .169 |
| Epoch [1] × Triplet type [High] × Age | 1.000 | 1.000 – 1.000 | -0.130 | 289756.368 | .897 |
| Epoch [2] × Triplet type [High] × Age | 1.000 | 1.000 – 1.000 | 0.168 | 289800.616 | .867 |
| Epoch [1] × Triplet type [High] × Parental education | 1.001 | 1.000 – 1.001 | 1.453 | 289761.391 | .146 |
| Epoch [2] × Triplet type [High] × Parental education | 1.000 | 0.999 – 1.001 | 0.060 | 289793.866 | .953 |
| **Random Effects** | | | | | |

| | |
|---|---|
| $\sigma^2$ | 0.035 |
| $\tau_{00\ \text{Participant}}$ | 0.012 |
| $\tau_{11\ \text{Participant.Epoch1}}$ | 0.001 |
| $\tau_{11\ \text{Participant.Epoch2}}$ | 0.000 |
| $\rho_{01\ \text{Participant.Epoch1}}$ | 0.167 |
| $\rho_{01\ \text{Participant.Epoch2}}$ | 0.073 |
| ICC | 0.265 |
| $N_{\text{Participant}}$ | 325 |
| Observations | 290682 |
| Marginal $R^2$ / Conditional $R^2$ | 0.013 / 0.275 |

**Supplementary Table S9. Results of the linear mixed model with Parental education on log transformed RT, learning.** Top table shows the Type 3 tests of fixed effects. Bottom table shows regression coefficients of fixed effects and summary information about the random effects. Coefficients are exponentiated to obtain the multiplicative factor for each 1 unit increase in the given independent variable. E.g., the coefficient of Age being 1.004 means that each year corresponds to a 0.4% increase in RTs. The marginal R-squared considers only the variance of the fixed effects, while the conditional R-squared takes both the fixed and random effects into account (based on Nakagawa et al., 2017). Degrees of freedom are based on Satterthwaite's approximation. Statistically significant terms are highlighted in bold. Terms in brackets indicate the level of factor that is contrasted against the reference level, which is Low triplets for the Triplet type factor and Epoch 3 for the Epoch factor.
Model equation in *lmer* syntax: log(RT) ~ Epoch*Triplet type*(Harshness + Unpredictability + Subj cSES + Age + Parental education) + (Epoch | Participant)

#### Table S10. Sensitivity model Learning Accuracy.

| LRT test of effects | $\chi^2$ | *df* | *p* |
|---|---|---|---|
| **Epoch** | 71.86 | 2 | **< .001** |
| **Triplet type** | 333.50 | 1 | **< .001** |
| Harshness | 0.50 | 1 | .480 |
| Unpredictability | 0.11 | 1 | .743 |
| Subj cSES | 0.77 | 1 | .382 |
| Age | 2.45 | 1 | .118 |
| Parental education | 1.20 | 1 | .273 |
| **Epoch x Triplet type** | 38.21 | 2 | **< .001** |
| Epoch x Harshness | 2.17 | 2 | .338 |
| Epoch x Unpredictability | 0.41 | 2 | .815 |
| Epoch x Subj cSES | 1.64 | 2 | .439 |
| Epoch x Age | 0.56 | 2 | .756 |
| Epoch x Parental education | 2.35 | 2 | .309 |
| Triplet type x Harshness | 0.01 | 1 | .945 |
| Triplet type x Unpredictability | 2.65 | 1 | .104 |
| Triplet type x Subj cSES | 3.13 | 1 | .077 |
| Triplet type x Age | 0.24 | 1 | .624 |
| Triplet type x Parental education | 0.48 | 1 | .488 |
| Epoch x Triplet type x Harshness | 1.68 | 2 | .432 |
| Epoch x Triplet type x Unpredictability | 2.20 | 2 | .332 |
| Epoch x Triplet type x Subj cSES | 0.00 | 2 | .999 |
| Epoch x Triplet type x Age | 1.88 | 2 | .391 |
| Epoch x Triplet type x Parental education | 0.39 | 2 | .821 |

| Fixed effects | *Log-odds* | *95% CI* | *z* | *df* | *p* |
|---|---|---|---|---|---|
| **(Intercept)** | 2.456 | 2.403 – 2.510 | 90.404 | Inf | **< .001** |
| **Epoch [1]** | 0.146 | 0.113 – 0.179 | 8.578 | Inf | **< .001** |
| **Epoch [2]** | -0.043 | -0.067 – -0.018 | -3.422 | Inf | **.001** |
| **Triplet type [High]** | 0.134 | 0.120 – 0.148 | 18.595 | Inf | **< .001** |
| Harshness | -0.021 | -0.078 – 0.036 | -0.716 | Inf | .474 |
| Unpredictability | 0.010 | -0.047 – 0.067 | 0.335 | Inf | .738 |
| cSES | -0.018 | -0.057 – 0.022 | -0.874 | Inf | .382 |
| Age | 0.008 | -0.002 – 0.018 | 1.569 | Inf | .117 |
| Parental education | 0.019 | -0.015 – 0.052 | 1.099 | Inf | .272 |
| **Epoch [1] × Triplet type [High]** | -0.048 | -0.068 – -0.027 | -4.572 | Inf | **< .001** |
| Epoch [2] × Triplet type [High] | -0.012 | -0.031 – 0.008 | -1.143 | Inf | .253 |
| Epoch [1] × Harshness | -0.005 | -0.040 – 0.031 | -0.253 | Inf | .800 |
| Epoch [2] × Harshness | 0.018 | -0.007 – 0.044 | 1.392 | Inf | .164 |
| Epoch [1] × Unpredictability | 0.011 | -0.024 – 0.046 | 0.610 | Inf | .542 |
| Epoch [2] × Unpredictability | -0.006 | -0.032 – 0.019 | -0.490 | Inf | .624 |
| Epoch [1] × cSES | 0.004 | -0.021 – 0.028 | 0.299 | Inf | .765 |

| | | | | | |
|---|---|---|---|---|---|
| Epoch [2] × cSES | -0.011 | -0.029 – 0.007 | -1.233 | Inf | .218 |
| Epoch [1] × Age | -0.002 | -0.009 – 0.004 | -0.723 | Inf | .470 |
| Epoch [2] × Age | 0.001 | -0.003 – 0.006 | 0.533 | Inf | .594 |
| Epoch [1] × Parental education | 0.016 | -0.005 – 0.037 | 1.530 | Inf | .126 |
| Epoch [2] × Parental education | -0.005 | -0.020 – 0.010 | -0.621 | Inf | .534 |
| Triplet type [High] × Harshness | -0.001 | -0.016 – 0.015 | -0.069 | Inf | .945 |
| Triplet type [High] × Unpredictability | 0.013 | -0.003 – 0.028 | 1.634 | Inf | .102 |
| Triplet type [High] × cSES | -0.010 | -0.020 – 0.001 | -1.776 | Inf | .076 |
| Triplet type [High] × Age | 0.001 | -0.002 – 0.003 | 0.492 | Inf | .622 |
| Triplet type [High] × Parental education | 0.003 | -0.006 – 0.012 | 0.695 | Inf | .487 |
| Epoch [1] × Triplet type [High] × Harshness | -0.013 | -0.035 – 0.009 | -1.128 | Inf | .259 |
| Epoch [2] × Triplet type [High] × Harshness | 0.001 | -0.021 – 0.022 | 0.053 | Inf | .958 |
| Epoch [1] × Triplet type [High] × Unpredictability | 0.008 | -0.014 – 0.031 | 0.746 | Inf | .455 |
| Epoch [2] × Triplet type [High] × Unpredictability | 0.008 | -0.014 – 0.029 | 0.687 | Inf | .492 |
| Epoch [1] × Triplet type [High] × cSES | -0.000 | -0.016 – 0.015 | -0.010 | Inf | .992 |
| Epoch [2] × Triplet type [High] × cSES | 0.000 | -0.015 – 0.015 | 0.005 | Inf | .996 |
| Epoch [1] × Triplet type [High] × Age | 0.002 | -0.002 – 0.006 | 0.776 | Inf | .438 |
| Epoch [2] × Triplet type [High] × Age | -0.003 | -0.007 – 0.001 | -1.372 | Inf | .170 |
| Epoch [1] × Triplet type [High] × Parental education | 0.004 | -0.009 – 0.017 | 0.631 | Inf | .528 |
| Epoch [2] × Triplet type [High] × Parental education | -0.002 | -0.015 – 0.010 | -0.330 | Inf | .741 |
| **Random Effects** | | | | | |
| $\sigma^2$ | 3.290 | | | | |

| | |
|---|---|
| $\tau_{00}$ Participant | 0.206 |
| $\tau_{11}$ Participant.Epoch1 | 0.048 |
| $\tau_{11}$ Participant.Epoch2 | 0.012 |
| $\rho_{01}$ Participant.Epoch1 | 0.392 |
| $\rho_{01}$ Participant.Epoch2 | -0.233 |
| ICC | 0.068 |
| N Participant | 325 |
| Observations | 316413 |
| Marginal $R^2$ / Conditional $R^2$ | 0.008 / 0.075 |

**Supplementary Table S10. Results of the binomial mixed model with Parental education on accuracy, learning.** Top table shows the LRT tests of fixed effects. Bottom table shows regression coefficients of fixed effects and summary information about the random effects. Coefficients are log odds, thus positive values indicate that the given independent variable is associated with an increase likelihood of correct responses, and negative values the opposite. P values for coefficients are from z tests, making the degrees of freedom infinite. The marginal R-squared considers only the variance of the fixed effects, while the conditional R-squared takes both the fixed and random effects into account (based on Nakagawa et al., 2017). Statistically significant terms are highlighted in bold. Terms in brackets indicate the level of factor that is contrasted against the reference level, which is Low triplets for the Triplet type factor and Epoch 3 for the Epoch factor.

Model equation in *lmer* syntax: Correct ~ Epoch*Triplet type*(Harshness + Unpredictability + cSES + Age + Parental education) + (Epoch | Participant)

### Table S11. Sensitivity model Consolidation Accuracy.

| Type III test of effects | *F* | *df* | *p* |
|---|---|---|---|
| **Epoch** | 492.92 | 1, 330.34 | **< .001** |
| **Triplet type** | 466.81 | 1, 193142.51 | **< .001** |
| Harshness | 0.54 | 1, 318.76 | .464 |
| Unpredictability | 2.00 | 1, 318.78 | .158 |
| Subj cSES | 0.12 | 1, 318.76 | .730 |
| **Age** | 17.97 | 1, 318.67 | **< .001** |
| **Delay duration** | 10.67 | 1, 318.76 | **.001** |
| Parental education | 0.00 | 1, 318.76 | .944 |
| Epoch x Triplet type | 0.33 | 1, 193146.61 | .566 |
| Epoch x Harshness | 1.08 | 1, 330.11 | .299 |
| Epoch x Unpredictability | 0.01 | 1, 330.36 | .932 |
| Epoch x Subj cSES | 3.12 | 1, 329.98 | .078 |
| Epoch x Age | 0.24 | 1, 328.55 | .622 |
| Epoch x Delay duration | 1.65 | 1, 330.03 | .201 |
| Epoch x Parental education | 0.02 | 1, 330.00 | .878 |
| Triplet type x Harshness | 0.03 | 1, 193137.03 | .865 |
| Triplet type x Unpredictability | 0.17 | 1, 193139.34 | .676 |
| Triplet type x Subj cSES | 0.50 | 1, 193146.01 | .480 |
| **Triplet type x Age** | 4.28 | 1, 193146.12 | **.039** |
| Triplet type x Delay duration | 2.95 | 1, 193138.49 | .086 |
| Triplet type x Parental education | 0.20 | 1, 193144.24 | .659 |
| Epoch x Triplet type x Harshness | 0.28 | 1, 193146.19 | .594 |
| Epoch x Triplet type x Unpredictability | 0.18 | 1, 193149.24 | .669 |
| Epoch x Triplet type x Subj cSES | 0.14 | 1, 193144.97 | .705 |
| Epoch x Triplet type x Age | 1.29 | 1, 193129.40 | .256 |
| Epoch x Triplet type x Delay duration | 0.33 | 1, 193146.30 | .568 |
| **Epoch x Triplet type x Parental education** | 3.86 | 1, 193149.20 | **.049** |

| Fixed effects | *exp(b)* | *95% CI* | *t* | *df* | *p* |
|---|---|---|---|---|---|
| **(Intercept)** | 364.906 | 360.723 – 369.137 | 1006.751 | 318.777 | **< .001** |
| **Epoch [3]** | 1.032 | 1.029 – 1.035 | 22.202 | 330.337 | **< .001** |
| **Triplet type [High]** | 0.990 | 0.989 – 0.991 | -21.606 | 193142.509 | **< .001** |
| Harshness | 1.005 | 0.992 – 1.017 | 0.734 | 318.763 | .464 |
| Unpredictability | 0.991 | 0.979 – 1.004 | -1.414 | 318.778 | .158 |
| Subj cSES | 1.001 | 0.993 – 1.010 | 0.345 | 318.757 | .730 |
| **Age** | 1.005 | 1.003 – 1.007 | 4.239 | 318.675 | **< .001** |
| **Delay duration** | 1.001 | 1.000 – 1.002 | 3.267 | 318.759 | **.001** |
| Parental education | 1.000 | 0.993 – 1.007 | -0.070 | 318.757 | .944 |
| Epoch [3] × Triplet type [High] | 1.000 | 0.999 – 1.001 | 0.574 | 193146.606 | .566 |
| Epoch [3] × Harshness | 0.998 | 0.995 – 1.001 | -1.040 | 330.111 | .299 |
| Epoch [3] × Unpredictability | 1.000 | 0.997 – 1.003 | 0.086 | 330.364 | .932 |
| Epoch [3] × Subj cSES | 1.002 | 1.000 – 1.004 | 1.768 | 329.981 | .078 |

| | | | | | |
|---|---|---|---|---|---|
| Epoch [3] × Age | 1.000 | 1.000 – 1.001 | 0.493 | 328.553 | .622 |
| Epoch [3] × Delay duration | 1.000 | 1.000 – 1.000 | -1.283 | 330.032 | .201 |
| Epoch [3] × Parental education | 1.000 | 0.998 – 1.002 | 0.154 | 330.003 | .878 |
| Triplet type [High] × Harshness | 1.000 | 0.999 – 1.001 | -0.170 | 193137.028 | .865 |
| Triplet type [High] × Unpredictability | 1.000 | 0.999 – 1.001 | 0.418 | 193139.341 | .676 |
| Triplet type [High] × Subj cSES | 1.000 | 0.999 – 1.000 | -0.707 | 193146.014 | .480 |
| **Triplet type [High] × Age** | 1.000 | 1.000 – 1.000 | -2.068 | 193146.116 | **.039** |
| Triplet type [High] × Delay duration | 1.000 | 1.000 – 1.000 | 1.717 | 193138.492 | .086 |
| Triplet type [High] × Parental education | 1.000 | 0.999 – 1.000 | -0.442 | 193144.240 | .659 |
| Epoch [3] × Triplet type [High] × Harshness | 1.000 | 0.999 – 1.001 | -0.533 | 193146.187 | .594 |
| Epoch [3] × Triplet type [High] × Unpredictability | 1.000 | 0.999 – 1.001 | 0.428 | 193149.239 | .669 |
| Epoch [3] × Triplet type [High] × Subj cSES | 1.000 | 0.999 – 1.001 | -0.379 | 193144.966 | .705 |
| Epoch [3] × Triplet type [High] × Age | 1.000 | 1.000 – 1.000 | 1.137 | 193129.396 | .256 |
| Epoch [3] × Triplet type [High] × Delay duration | 1.000 | 1.000 – 1.000 | -0.571 | 193146.300 | .568 |
| **Epoch [3] × Triplet type [High] × Parental education** | 0.999 | 0.999 – 1.000 | -1.965 | 193149.200 | **.049** |

| **Random Effects** | |
|---|---|
| $\sigma^2$ | 0.030 |
| $\tau_{00\ \text{Participant}}$ | 0.010 |
| $\tau_{11\ \text{Participant.Epoch3}}$ | 0.001 |
| $\rho_{01\ \text{Participant.Epoch3}}$ | 0.112 |
| ICC | 0.263 |
| $N_{\text{Participant}}$ | 325 |
| Observations | 193772 |
| Marginal $R^2$ / Conditional $R^2$ | 0.049 / 0.299 |

**Supplementary Table S11. Results of the linear mixed model with Parental education on log transformed RT, consolidation.** Top table shows the Type 3 tests of fixed effects. Bottom

table shows regression coefficients of fixed effects and summary information about the random effects. Coefficients are exponentiated to obtain the multiplicative factor for each 1 unit increase in the given independent variable. E.g., the coefficient of Age being 1.005 means that each year corresponds to a 0.5% increase in RTs. The marginal R-squared considers only the variance of the fixed effects, while the conditional R-squared takes both the fixed and random effects into account (based on Nakagawa et al., 2017). Degrees of freedom are based on Satterthwaite's approximation. Statistically significant terms are highlighted in bold. Terms in brackets indicate the level of factor that is contrasted against the reference level, which is Low triplets for the Triplet type factor and Epoch 4 for the Epoch factor.
Model equation in *lmer* syntax: log(RT) ~ Epoch*Triplet type*(Harshness + Unpredictability + cSES + Age + Delay duration + Parental education) + (Epoch | Participant)

### Table S12. Sensitivity model Consolidation Accuracy.

| LRT test of effects | $\chi^2$ | *df* | *p* |
|---|---|---|---|
| **Epoch** | 32.69 | 1 | **< .001** |
| **Triplet type** | 482.41 | 1 | **< .001** |
| Harshness | 0.52 | 1 | .470 |
| Unpredictability | 0.07 | 1 | .793 |
| Subj cSES | 0.17 | 1 | .678 |
| Age | 3.76 | 1 | .053 |
| Delay duration | 2.82 | 1 | .093 |
| Parental education | 0.04 | 1 | .846 |
| Epoch x Triplet type | 0.01 | 1 | .916 |
| Epoch x Harshness | 1.29 | 1 | .256 |
| Epoch x Unpredictability | 1.08 | 1 | .299 |
| Epoch x Subj cSES | 0.07 | 1 | .787 |
| Epoch x Age | 0.24 | 1 | .622 |
| Epoch x Delay duration | 0.15 | 1 | .697 |
| Epoch x Parental education | 0.32 | 1 | .575 |
| Triplet type x Harshness | 0.42 | 1 | .516 |
| Triplet type x Unpredictability | 2.08 | 1 | .149 |
| Triplet type x Subj cSES | 1.09 | 1 | .297 |
| Triplet type x Age | 1.08 | 1 | .298 |
| Triplet type x Delay duration | 0.07 | 1 | .795 |
| Triplet type x Parental education | 2.36 | 1 | .124 |
| Epoch x Triplet type x Harshness | 0.27 | 1 | .603 |
| Epoch x Triplet type x Unpredictability | 2.40 | 1 | .121 |
| Epoch x Triplet type x Subj cSES | 0.47 | 1 | .494 |
| Epoch x Triplet type x Age | 0.01 | 1 | .930 |
| Epoch x Triplet type x Delay duration | 1.10 | 1 | .294 |
| Epoch x Triplet type x Parental education | 1.50 | 1 | .220 |

| Fixed effects | *Log-odds* | *95% CI* | *z* | *df* | *p* |
|---|---|---|---|---|---|
| **(Intercept)** | 2.413 | 2.365 – 2.462 | 97.103 | Inf | **< .001** |
| **Epoch [3]** | -0.070 | -0.093 – -0.047 | -5.879 | Inf | **< .001** |
| **Triplet type [High]** | 0.195 | 0.178 – 0.212 | 22.436 | Inf | **< .001** |
| Harshness | -0.019 | -0.072 – 0.033 | -0.725 | Inf | .468 |
| Unpredictability | -0.007 | -0.059 – 0.045 | -0.259 | Inf | .796 |
| Subj cSES | -0.008 | -0.043 – 0.028 | -0.419 | Inf | .676 |
| Age | 0.009 | -0.000 – 0.019 | 1.941 | Inf | .052 |
| Delay duration | 0.003 | -0.000 – 0.006 | 1.682 | Inf | .093 |
| Parental education | 0.003 | -0.027 – 0.034 | 0.195 | Inf | .846 |
| Epoch [3] × Triplet type [High] | -0.001 | -0.018 – 0.016 | -0.106 | Inf | .916 |
| Epoch [3] × Harshness | -0.014 | -0.039 – 0.010 | -1.139 | Inf | .255 |
| Epoch [3] × Unpredictability | 0.013 | -0.011 – 0.037 | 1.040 | Inf | .298 |
| Epoch [3] × Subj cSES | -0.002 | -0.019 – 0.015 | -0.270 | Inf | .787 |
| Epoch [3] × Age | 0.001 | -0.003 – 0.006 | 0.494 | Inf | .621 |

| | | | | | |
|---|---|---|---|---|---|
| Epoch [3] × Delay duration | -0.000 | -0.002 – 0.001 | -0.390 | Inf | .697 |
| Epoch [3] × Parental education | 0.004 | -0.010 – 0.019 | 0.563 | Inf | .574 |
| Triplet type [High] × Harshness | 0.006 | -0.012 – 0.024 | 0.653 | Inf | .514 |
| Triplet type [High] × Unpredictability | 0.013 | -0.005 – 0.031 | 1.450 | Inf | .147 |
| Triplet type [High] × Subj cSES | -0.007 | -0.019 – 0.006 | -1.048 | Inf | .295 |
| Triplet type [High] × Age | 0.002 | -0.002 – 0.005 | 1.047 | Inf | .295 |
| Triplet type [High] × Delay duration | -0.000 | -0.001 – 0.001 | -0.261 | Inf | .794 |
| Triplet type [High] × Parental education | 0.008 | -0.002 – 0.019 | 1.543 | Inf | .123 |
| Epoch [3] × Triplet type [High] × Harshness | 0.005 | -0.013 – 0.023 | 0.522 | Inf | .602 |
| Epoch [3] × Triplet type [High] × Unpredictability | -0.014 | -0.032 – 0.004 | -1.555 | Inf | .120 |
| Epoch [3] × Triplet type [High] × Subj cSES | -0.004 | -0.017 – 0.008 | -0.687 | Inf | .492 |
| Epoch [3] × Triplet type [High] × Age | 0.000 | -0.003 – 0.003 | 0.088 | Inf | .930 |
| Epoch [3] × Triplet type [High] × Delay duration | 0.001 | -0.000 – 0.002 | 1.053 | Inf | .292 |
| Epoch [3] × Triplet type [High] × Parental education | -0.007 | -0.018 – 0.004 | -1.231 | Inf | .218 |
| **Random Effects** | | | | | |
| $\sigma^2$ | 3.290 | | | | |
| $\tau_{00\ Participant}$ | 0.161 | | | | |
| $\tau_{11\ Participant.Epoch3}$ | 0.017 | | | | |
| $\rho_{01\ Participant.Epoch3}$ | 0.058 | | | | |
| ICC | 0.051 | | | | |
| $N_{Participant}$ | 325 | | | | |
| Observations | 211110 | | | | |
| Marginal $R^2$ / Conditional $R^2$ | 0.012 / 0.062 | | | | |

**Supplementary Table S12. Results of the binomial mixed model with Parental education on accuracy, consolidation.** Top table shows the LRT tests of fixed effects. Bottom table shows regression coefficients of fixed effects and summary information about the random effects. Coefficients are log odds, thus positive values indicate that the given independent

variable is associated with an increase likelihood of correct responses, and negative values the opposite. P values for coefficients are from z tests, making the degrees of freedom infinite. The marginal R-squared considers only the variance of the fixed effects, while the conditional R-squared takes both the fixed and random effects into account (based on Nakagawa et al., 2017). Statistically significant terms are highlighted in bold. Terms in brackets indicate the level of factor that is contrasted against the reference level, which is Low triplets for the Triplet type factor and Epoch 4 for the Epoch factor.
Model equation in *lmer* syntax: Correct ~ Epoch*Triplet type*(Harshness + Unpredictability + Subj cSES + Age + Delay duration + Parental education) + (Epoch | Participant)

### Table S13. Sensitivity model Rewiring Old knowledge RT.

| Type III test of effects | *F* | *df* | *p* |
|---|---|---|---|
| **Epoch** | 97.94 | 2, 363.36 | **< .001** |
| **Triplet type** | 482.17 | 1, 195582.45 | **< .001** |
| Harshness | 1.09 | 1, 320.41 | .298 |
| Unpredictability | 1.73 | 1, 320.44 | .189 |
| Subj cSES | 0.03 | 1, 320.38 | .854 |
| **Age** | 16.73 | 1, 320.35 | **< .001** |
| Parental education | 0.07 | 1, 320.37 | .787 |
| Epoch x Triplet type | 1.95 | 2, 195586.05 | .142 |
| Epoch x Harshness | 0.02 | 2, 364.84 | .979 |
| Epoch x Unpredictability | 1.27 | 2, 365.85 | .281 |
| Epoch x Subj cSES | 0.77 | 2, 363.28 | .462 |
| Epoch x Age | 0.41 | 2, 362.06 | .662 |
| Epoch x Parental education | 0.35 | 2, 362.78 | .703 |
| Triplet type x Harshness | 0.63 | 1, 195579.41 | .426 |
| Triplet type x Unpredictability | 0.15 | 1, 195589.98 | .696 |
| Triplet type x Subj cSES | 0.69 | 1, 195543.58 | .405 |
| **Triplet type x Age** | 5.29 | 1, 195574.53 | **.021** |
| Triplet type x Parental education | 0.09 | 1, 195591.71 | .767 |
| Epoch x Triplet type x Harshness | 0.71 | 2, 195573.22 | .490 |
| Epoch x Triplet type x Unpredictability | 0.47 | 2, 195589.84 | .627 |
| Epoch x Triplet type x Subj cSES | 1.67 | 2, 195585.75 | .188 |
| Epoch x Triplet type x Age | 1.41 | 2, 195583.41 | .245 |
| Epoch x Triplet type x Parental education | 0.23 | 2, 195595.11 | .791 |

| Fixed effects | *exp(b)* | *95% CI* | *t* | *df* | *p* |
|---|---|---|---|---|---|
| **(Intercept)** | 360.377 | 356.355 – 364.444 | 1032.095 | 320.379 | **< .001** |
| **Epoch [4]** | 0.980 | 0.977 – 0.983 | -13.988 | 373.640 | **< .001** |
| **Epoch [5]** | 1.008 | 1.005 – 1.010 | 6.499 | 346.463 | **< .001** |
| **Triplet type [H L]** | 0.989 | 0.988 – 0.990 | -21.958 | 195582.453 | **< .001** |
| Harshness | 1.006 | 0.994 – 1.019 | 1.042 | 320.413 | .298 |
| Unpredictability | 0.992 | 0.980 – 1.004 | -1.315 | 320.437 | .189 |
| Subj cSES | 0.999 | 0.991 – 1.008 | -0.184 | 320.381 | .854 |
| **Age** | 1.004 | 1.002 – 1.007 | 4.091 | 320.347 | **< .001** |
| Parental education | 0.999 | 0.992 – 1.006 | -0.270 | 320.369 | .787 |
| Epoch [4] × Triplet type [H L] | 1.000 | 0.999 – 1.002 | 0.265 | 195554.949 | .791 |
| Epoch [5] × Triplet type [H L] | 1.001 | 1.000 – 1.003 | 1.455 | 195348.623 | .146 |
| Epoch [4] × Harshness | 1.000 | 0.997 – 1.003 | -0.058 | 373.758 | .954 |
| Epoch [5] × Harshness | 1.000 | 0.998 – 1.003 | 0.203 | 348.407 | .839 |
| Epoch [4] × Unpredictability | 0.999 | 0.996 – 1.002 | -0.428 | 374.783 | .669 |
| Epoch [5] × Unpredictability | 1.002 | 0.999 – 1.005 | 1.571 | 349.596 | .117 |
| Epoch [4] × Subj cSES | 1.000 | 0.998 – 1.002 | -0.331 | 373.477 | .741 |

| | | | | | |
|---|---|---|---|---|---|
| Epoch [5] × Subj cSES | 0.999 | 0.997 – 1.001 | -0.937 | 345.955 | .349 |
| Epoch [4] × Age | 1.000 | 0.999 – 1.000 | -0.246 | 372.403 | .806 |
| Epoch [5] × Age | 1.000 | 0.999 – 1.000 | -0.681 | 344.963 | .496 |
| Epoch [4] × Parental education | 1.000 | 0.998 – 1.002 | -0.312 | 373.158 | .756 |
| Epoch [5] × Parental education | 1.000 | 0.998 – 1.001 | -0.568 | 345.208 | .570 |
| Triplet type [H L] × Harshness | 1.000 | 0.999 – 1.002 | 0.796 | 195579.405 | .426 |
| Triplet type [H L] × Unpredictability | 1.000 | 0.999 – 1.001 | -0.390 | 195589.984 | .696 |
| Triplet type [H L] × Subj cSES | 1.000 | 1.000 – 1.001 | 0.833 | 195543.577 | .405 |
| **Triplet type [H L] × Age** | 1.000 | 1.000 – 1.000 | -2.300 | 195574.530 | **.021** |
| Triplet type [H L] × Parental education | 1.000 | 0.999 – 1.001 | 0.296 | 195591.709 | .767 |
| Epoch [4] × Triplet type [H L] × Harshness | 0.999 | 0.998 – 1.001 | -1.129 | 195531.447 | .259 |
| Epoch [5] × Triplet type [H L] × Harshness | 1.001 | 0.999 – 1.003 | 0.962 | 195485.755 | .336 |
| Epoch [4] × Triplet type [H L] × Unpredictability | 1.001 | 0.999 – 1.002 | 0.742 | 195554.903 | .458 |
| Epoch [5] × Triplet type [H L] × Unpredictability | 0.999 | 0.997 – 1.001 | -0.931 | 195473.113 | .352 |
| Epoch [4] × Triplet type [H L] × Subj cSES | 1.001 | 1.000 – 1.002 | 1.011 | 195580.231 | .312 |
| Epoch [4] × Triplet type [H L] × Subj cSES | 0.999 | 0.998 – 1.000 | -1.827 | 195164.125 | .068 |
| Epoch [4] × Triplet type [H L] × Age | 1.000 | 1.000 – 1.000 | -0.540 | 195547.653 | .589 |
| Epoch [5] × Triplet type [H L] × Age | 1.000 | 1.000 – 1.001 | 1.611 | 195322.701 | .107 |
| Epoch [4] × Triplet type [H L] × Parental education | 1.000 | 0.999 – 1.001 | 0.669 | 195569.489 | .504 |
| Epoch [5] × Triplet type [H L] × Parental education | 1.000 | 0.999 – 1.001 | -0.260 | 195378.324 | .795 |
| **Random Effects** | | | | | |

| | |
|---|---|
| $\sigma^2$ | 0.028 |
| $\tau_{00\ \text{Participant}}$ | 0.010 |
| $\tau_{11\ \text{Participant.Epoch4}}$ | < 0.001 |
| $\tau_{11\ \text{Participant.Epoch5}}$ | < 0.001 |
| $\rho_{01\ \text{Participant.Epoch4}}$ | 0.029 |
| $\rho_{01\ \text{Participant.Epoch5}}$ | -0.061 |
| ICC | 0.266 |
| $N_{\text{Participant}}$ | 325 |
| Observations | 196226 |
| Marginal $R^2$ / Conditional $R^2$ | 0.024 / 0.284 |

**Supplementary Table S13. Results of the linear mixed model with Parental education on log transformed RT, rewiring old knowledge.** Top table shows the Type 3 tests of fixed effects. Bottom table shows regression coefficients of fixed effects and summary information about the random effects. Coefficients are exponentiated to obtain the multiplicative factor for each 1 unit increase in the given independent variable. E.g., the coefficient of Age being 1.005 means that each year corresponds to a 0.5% increase in RTs. The marginal R-squared considers only the variance of the fixed effects, while the conditional R-squared takes both the fixed and random effects into account (based on Nakagawa et al., 2017). Degrees of freedom are based on Satterthwaite's approximation. Statistically significant terms are highlighted in bold. Terms in brackets indicate the level of factor that is contrasted against the reference level, which is L L triplets for the Triplet type factor and Epoch 6 for the Epoch factor.

Model equation in *lmer* syntax: log(RT) ~ Epoch*Triplet type*(Harshness + Unpredictability + Subj cSES + Age + Parental education) + (Epoch | Participant)

**Table S14. Sensitivity model Rewiring Old knowledge Accuracy.**

| LRT test of effects | $\chi^2$ | *df* | *p* |
|---|---|---|---|
| **Epoch** | 9.33 | 2 | **.009** |
| **Triplet type** | 212.13 | 1 | **< .001** |
| Harshness | 0.03 | 1 | .852 |
| Unpredictability | 0.64 | 1 | .422 |
| Subj cSES | 2.96 | 1 | .085 |
| Age | 2.91 | 1 | .088 |
| Parental education | 0.32 | 1 | .574 |
| **Epoch x Triplet type** | 21.13 | 2 | **< .001** |
| Epoch x Harshness | 0.49 | 2 | .781 |
| Epoch x Unpredictability | 0.84 | 2 | .656 |
| **Epoch x Subj cSES** | 6.96 | 2 | **.031** |
| Epoch x Age | 0.20 | 2 | .906 |
| Epoch x Parental education | 0.22 | 2 | .894 |
| Triplet type x Harshness | 0.06 | 1 | .809 |
| **Triplet type x Unpredictability** | 3.96 | 1 | **.047** |
| Triplet type x Subj cSES | 0.97 | 1 | .324 |
| Triplet type x Age | 0.35 | 1 | .554 |
| Triplet type x Parental education | 1.53 | 1 | .217 |
| Epoch x Triplet type x Harshness | 0.08 | 2 | .962 |
| Epoch x Triplet type x Unpredictability | 1.56 | 2 | .459 |
| Epoch x Triplet type x Subj cSES | 1.72 | 2 | .423 |
| Epoch x Triplet type x Age | 0.71 | 2 | .701 |
| Epoch x Triplet type x Parental education | 1.82 | 2 | .403 |

| Fixed effects | *Log-odds* | *95% CI* | *z* | *df* | *p* |
|---|---|---|---|---|---|
| **(Intercept)** | 2.427 | 2.380 – 2.474 | 101.504 | Inf | **< .001** |
| **Epoch [4]** | 0.053 | 0.019 – 0.087 | 3.071 | Inf | **.002** |
| Epoch [5] | -0.031 | -0.066 – 0.005 | -1.696 | Inf | .090 |
| **Triplet type [H L]** | 0.153 | 0.132 – 0.173 | 14.779 | Inf | **< .001** |
| Harshness | 0.005 | -0.045 – 0.055 | 0.188 | Inf | .850 |
| Unpredictability | -0.021 | -0.071 – 0.029 | -0.807 | Inf | .420 |
| Subj cSES | -0.030 | -0.064 – 0.004 | -1.728 | Inf | .084 |
| Age | 0.008 | -0.001 – 0.017 | 1.710 | Inf | .087 |
| Parental education | -0.008 | -0.037 – 0.021 | -0.564 | Inf | .573 |
| **Epoch [4] × Triplet type [H L]** | 0.043 | 0.016 – 0.070 | 3.079 | Inf | **.002** |
| **Epoch [5] × Triplet type [H L]** | -0.074 | -0.105 – -0.043 | -4.611 | Inf | **< .001** |
| Epoch [4] × Harshness | -0.010 | -0.045 – 0.025 | -0.571 | Inf | .568 |
| Epoch [5] × Harshness | -0.000 | -0.037 – 0.036 | -0.020 | Inf | .984 |
| Epoch [4] × Unpredictability | -0.005 | -0.040 – 0.030 | -0.276 | Inf | .783 |
| Epoch [5] × Unpredictability | 0.016 | -0.020 – 0.052 | 0.883 | Inf | .377 |
| Epoch [4] × Subj cSES | 0.008 | -0.016 – 0.032 | 0.625 | Inf | .532 |

| | | | | | |
|---|---|---|---|---|---|
| Epoch [5] × Subj cSES | 0.023 | -0.002 – 0.047 | 1.778 | Inf | .075 |
| Epoch [4] × Age | 0.001 | -0.005 – 0.008 | 0.325 | Inf | .745 |
| Epoch [5] × Age | -0.001 | -0.008 – 0.005 | -0.434 | Inf | .664 |
| Epoch [4] × Parental education | -0.000 | -0.021 – 0.020 | -0.045 | Inf | .964 |
| Epoch [5] × Parental education | -0.004 | -0.025 – 0.017 | -0.366 | Inf | .715 |
| Triplet type [H L] × Harshness | -0.003 | -0.024 – 0.019 | -0.242 | Inf | .809 |
| **Triplet type [H L] × Unpredictability** | 0.022 | 0.000 – 0.043 | 2.000 | Inf | **.045** |
| Triplet type [H L] × Subj cSES | 0.007 | -0.007 – 0.022 | 0.991 | Inf | .322 |
| Triplet type [H L] × Age | -0.001 | -0.005 – 0.003 | -0.594 | Inf | .553 |
| Triplet type [H L] × Parental education | 0.008 | -0.005 – 0.020 | 1.242 | Inf | .214 |
| Epoch [4] × Triplet type [H L] × Harshness | 0.000 | -0.028 – 0.029 | 0.023 | Inf | .982 |
| Epoch [5] × Triplet type [H L] × Harshness | 0.004 | -0.030 – 0.037 | 0.212 | Inf | .832 |
| Epoch [4] × Triplet type [H L] × Unpredictability | 0.011 | -0.017 – 0.040 | 0.787 | Inf | .431 |
| Epoch [5] × Triplet type [H L] × Unpredictability | -0.021 | -0.055 – 0.012 | -1.254 | Inf | .210 |
| Epoch [4] × Triplet type [H L] × Subj cSES | 0.011 | -0.009 – 0.031 | 1.105 | Inf | .269 |
| Epoch [4] × Triplet type [H L] × Subj cSES | -0.014 | -0.037 – 0.008 | -1.233 | Inf | .218 |
| Epoch [4] × Triplet type [H L] × Age | 0.000 | -0.005 – 0.006 | 0.128 | Inf | .898 |
| Epoch [5] × Triplet type [H L] × Age | 0.002 | -0.004 – 0.008 | 0.605 | Inf | .545 |
| Epoch [4] × Triplet type [H L] × Parental education | 0.011 | -0.005 – 0.028 | 1.327 | Inf | .184 |
| Epoch [5] × Triplet type [H L] × Parental education | -0.010 | -0.029 – 0.010 | -1.003 | Inf | .316 |
| **Random Effects** | | | | | |
| $\sigma^2$ | 3.290 | | | | |

| | |
|---|---|
| $\tau_{00\ Participant}$ | 0.137 |
| $\tau_{11\ Participant.Epoch4}$ | 0.026 |
| $\tau_{11\ Participant.Epoch5}$ | 0.012 |
| $\rho_{01\ Participant.Epoch4}$ | 0.086 |
| $\rho_{01\ Participant.Epoch5}$ | -0.149 |
| ICC | 0.047 |
| $N_{Participant}$ | 325 |
| Observations | 212870 |
| Marginal $R^2$ / Conditional $R^2$ | 0.008 / 0.055 |

**Supplementary Table S14. Results of the binomial mixed model with Parental education on accuracy, rewiring old knowledge.** Top table shows the LRT tests of fixed effects. Bottom table shows regression coefficients of fixed effects and summary information about the random effects. Coefficients are log odds, thus positive values indicate that the given independent variable is associated with an increase likelihood of correct responses, and negative values the opposite. P values for coefficients are from z tests, making the degrees of freedom infinite. The marginal R-squared considers only the variance of the fixed effects, while the conditional R-squared takes both the fixed and random effects into account (based on Nakagawa et al., 2017). Statistically significant terms are highlighted in bold. Terms in brackets indicate the level of factor that is contrasted against the reference level, which is L L triplets for the Triplet type factor and Epoch 6 for the Epoch factor.
Model equation in *lmer* syntax: Correct ~ Epoch*Triplet type*(Harshness + Unpredictability + Subj cSES + Age + Parental education) + (Epoch | Participant)

### Table S15. Sensitivity model Rewiring New knowledge RT.

| Type III test of effects | *F* | *df* | *p* |
|---|---|---|---|
| **Epoch** | 86.17 | 2, 352.83 | **< .001** |
| **Triplet type** | 39.10 | 1, 137391.96 | **< .001** |
| Harshness | 0.88 | 1, 319.77 | .348 |
| Unpredictability | 1.61 | 1, 319.83 | .205 |
| Subj cSES | 0.00 | 1, 319.74 | .970 |
| **Age** | 17.24 | 1, 319.71 | **< .001** |
| Parental education | 0.07 | 1, 319.74 | .791 |
| Epoch x Triplet type | 2.15 | 2, 137322.46 | .117 |
| Epoch x Harshness | 0.05 | 2, 353.85 | .955 |
| Epoch x Unpredictability | 2.50 | 2, 356.23 | .084 |
| Epoch x Subj cSES | 0.15 | 2, 352.00 | .863 |
| Epoch x Age | 1.36 | 2, 351.23 | .258 |
| Epoch x Parental education | 0.35 | 2, 352.17 | .704 |
| Triplet type x Harshness | 0.17 | 1, 137365.33 | .683 |
| Triplet type x Unpredictability | 0.01 | 1, 137418.62 | .918 |
| **Triplet type x Subj cSES** | 8.51 | 1, 137413.57 | **.004** |
| Triplet type x Age | 3.23 | 1, 137418.43 | .072 |
| Triplet type x Parental education | 0.07 | 1, 137428.86 | .795 |
| Epoch x Triplet type x Harshness | 0.05 | 2, 137299.73 | .956 |
| Epoch x Triplet type x Unpredictability | 0.11 | 2, 137323.23 | .899 |
| Epoch x Triplet type x Subj cSES | 0.50 | 2, 137328.25 | .606 |
| Epoch x Triplet type x Age | 1.07 | 2, 137335.17 | .343 |
| Epoch x Triplet type x Parental education | 0.03 | 2, 137327.83 | .968 |

| Fixed effects | *exp(b)* | *95% CI* | *t* | *df* | *p* |
|---|---|---|---|---|---|
| **(Intercept)** | 363.244 | 359.221 – 367.313 | 1041.255 | 319.751 | **< .001** |
| **Epoch [4]** | 0.981 | 0.978 – 0.984 | -13.111 | 325.487 | **< .001** |
| **Epoch [5]** | 1.007 | 1.005 – 1.009 | 6.138 | 397.899 | **< .001** |
| **Triplet type [L H]** | 0.996 | 0.995 – 0.998 | -6.253 | 137391.962 | **< .001** |
| Harshness | 1.006 | 0.994 – 1.018 | 0.939 | 319.771 | .348 |
| Unpredictability | 0.992 | 0.980 – 1.004 | -1.270 | 319.834 | .205 |
| Subj cSES | 1.000 | 0.992 – 1.008 | 0.038 | 319.737 | .970 |
| **Age** | 1.005 | 1.002 – 1.007 | 4.152 | 319.706 | **< .001** |
| Parental education | 0.999 | 0.992 – 1.006 | -0.265 | 319.739 | .791 |
| Epoch [4] × Triplet type [L H] | 1.002 | 1.000 – 1.003 | 1.813 | 137366.153 | .070 |
| Epoch [5] × Triplet type [L H] | 1.000 | 0.999 – 1.002 | 0.116 | 137287.379 | .908 |
| Epoch [4] × Harshness | 1.000 | 0.997 – 1.004 | 0.281 | 324.762 | .779 |
| Epoch [5] × Harshness | 1.000 | 0.997 – 1.002 | -0.222 | 400.464 | .824 |
| Epoch [4] × Unpredictability | 0.999 | 0.996 – 1.002 | -0.649 | 327.731 | .517 |
| **Epoch [5] × Unpredictability** | 1.003 | 1.000 – 1.005 | 2.212 | 403.537 | **.028** |
| Epoch [4] × Subj cSES | 1.000 | 0.997 – 1.002 | -0.417 | 324.917 | .677 |

| | | | | | |
|---|---|---|---|---|---|
| Epoch [5] × Subj cSES | 1.000 | 0.998 – 1.001 | -0.140 | 395.697 | .889 |
| Epoch [4] × Age | 1.000 | 1.000 – 1.001 | 0.399 | 322.467 | .690 |
| Epoch [5] × Age | 1.000 | 0.999 – 1.000 | -1.621 | 397.186 | .106 |
| Epoch [4] × Parental education | 0.999 | 0.998 – 1.001 | -0.569 | 325.253 | .570 |
| Epoch [5] × Parental education | 1.000 | 0.998 – 1.001 | -0.319 | 396.952 | .750 |
| Triplet type [L H] × Harshness | 1.000 | 0.999 – 1.001 | -0.408 | 137365.331 | .683 |
| Triplet type [L H] × Unpredictability | 1.000 | 0.999 – 1.001 | 0.103 | 137418.617 | .918 |
| **Triplet type [L H] × Subj cSES** | 1.001 | 1.000 – 1.002 | 2.917 | 137413.566 | **.004** |
| Triplet type [L H] × Age | 1.000 | 1.000 – 1.000 | -1.796 | 137418.434 | .072 |
| Triplet type [L H] × Parental education | 1.000 | 0.999 – 1.001 | 0.259 | 137428.856 | .795 |
| Epoch [4] × Triplet type [L H] × Harshness | 1.000 | 0.998 – 1.002 | -0.267 | 137339.081 | .789 |
| Epoch [5] × Triplet type [L H] × Harshness | 1.000 | 0.999 – 1.002 | 0.240 | 137272.783 | .810 |
| Epoch [4] × Triplet type [L H] × Unpredictability | 1.000 | 0.999 – 1.002 | 0.407 | 137372.422 | .684 |
| Epoch [5] × Triplet type [L H] × Unpredictability | 1.000 | 0.998 – 1.001 | -0.374 | 137289.895 | .708 |
| Epoch [4] × Triplet type [L H] × Subj cSES | 1.001 | 0.999 – 1.002 | 0.955 | 137370.064 | .339 |
| Epoch [4] × Triplet type [L H] × Subj cSES | 1.000 | 0.999 – 1.001 | -0.683 | 137313.390 | .494 |
| Epoch [4] × Triplet type [L H] × Age | 1.000 | 1.000 – 1.000 | 0.896 | 137414.216 | .370 |
| Epoch [5] × Triplet type [L H] × Age | 1.000 | 1.000 – 1.000 | 0.656 | 137313.510 | .512 |
| Epoch [4] × Triplet type [L H] × Parental education | 1.000 | 0.999 – 1.001 | 0.067 | 137369.476 | .947 |
| Epoch [5] × Triplet type [L H] × Parental education | 1.000 | 0.999 – 1.001 | 0.192 | 137319.257 | .848 |
| **Random Effects** | | | | | |
| $\sigma^2$ | 0.028 | | | | |

| | |
|---|---|
| $\tau_{00}$ Participant | 0.010 |
| $\tau_{11}$ Participant.Epoch4 | < 0.001 |
| $\tau_{11}$ Participant.Epoch5 | < 0.001 |
| $\rho_{01}$ Participant.Epoch4 | 0.051 |
| $\rho_{01}$ Participant.Epoch5 | 0.074 |
| ICC | 0.260 |
| N Participant | 325 |
| Observations | 137887 |
| Marginal $R^2$ / Conditional $R^2$ | 0.018 / 0.273 |

**Supplementary Table S15. Results of the linear mixed model with Parental education on log transformed RT, rewiring new knowledge.** Top table shows the Type 3 tests of fixed effects. Bottom table shows regression coefficients of fixed effects and summary information about the random effects. Coefficients are exponentiated to obtain the multiplicative factor for each 1 unit increase in the given independent variable. E.g., the coefficient of Age being 1.005 means that each year corresponds to a 0.5% increase in RTs. The marginal R-squared considers only the variance of the fixed effects, while the conditional R-squared takes both the fixed and random effects into account (based on Nakagawa et al., 2017). Degrees of freedom are based on Satterthwaite's approximation. Statistically significant terms are highlighted in bold. Terms in brackets indicate the level of factor that is contrasted against the reference level, which is L L triplets for the Triplet type factor and Epoch 6 for the Epoch factor.
Model equation in *lmer* syntax: log(RT) ~ Epoch*Triplet type*(Harshness + Unpredictability + Subj cSES + Age + Parental education) + (Epoch | Participant)

### Table S16. Sensitivity model Rewiring New knowledge Accuracy.

| LRT test of effects | $\chi^2$ | *df* | *p* |
|---|---|---|---|
| **Epoch** | 43.96 | 2 | **< .001** |
| Triplet type | 2.65 | 1 | .104 |
| Harshness | 0.06 | 1 | .809 |
| Unpredictability | 0.52 | 1 | .470 |
| Subj cSES | 0.39 | 1 | .532 |
| Age | 1.11 | 1 | .292 |
| Parental education | 0.17 | 1 | .678 |
| **Epoch x Triplet type** | 14.54 | 2 | **< .001** |
| Epoch x Harshness | 0.54 | 2 | .764 |
| Epoch x Unpredictability | 3.64 | 2 | .162 |
| Epoch x Subj cSES | 0.80 | 2 | .669 |
| Epoch x Age | 0.45 | 2 | .800 |
| Epoch x Parental education | 0.88 | 2 | .643 |
| Triplet type x Harshness | 1.57 | 1 | .210 |
| **Triplet type x Unpredictability** | 4.89 | 1 | **.027** |
| **Triplet type x Subj cSES** | 11.96 | 1 | **< .001** |
| **Triplet type x Age** | 3.87 | 1 | **.049** |
| Triplet type x Parental education | 1.61 | 1 | .204 |
| Epoch x Triplet type x Harshness | 0.27 | 2 | .874 |
| Epoch x Triplet type x Unpredictability | 2.56 | 2 | .278 |
| **Epoch x Triplet type x Subj cSES** | 12.26 | 2 | **.002** |
| Epoch x Triplet type x Age | 0.18 | 2 | .915 |
| Epoch x Triplet type x Parental education | 0.67 | 2 | .716 |

| Fixed effects | *Log-odds* | *95% CI* | *z* | *df* | *p* |
|---|---|---|---|---|---|
| **(Intercept)** | 2.291 | 2.244 – 2.337 | 96.980 | Inf | **< .001** |
| Epoch [4] | -0.004 | -0.043 – 0.034 | -0.222 | Inf | .824 |
| **Epoch [5]** | 0.100 | 0.068 – 0.133 | 6.039 | Inf | **< .001** |
| Triplet type [L H] | 0.018 | -0.003 – 0.039 | 1.636 | Inf | .102 |
| Harshness | -0.006 | -0.056 – 0.044 | -0.241 | Inf | .810 |
| Unpredictability | -0.018 | -0.068 – 0.031 | -0.724 | Inf | .469 |
| Subj cSES | -0.011 | -0.045 – 0.023 | -0.626 | Inf | .531 |
| Age | 0.005 | -0.004 – 0.014 | 1.056 | Inf | .291 |
| Parental education | -0.006 | -0.035 – 0.023 | -0.416 | Inf | .678 |
| Epoch [4] × Triplet type [L H] | -0.020 | -0.051 – 0.011 | -1.234 | Inf | .217 |
| **Epoch [5] × Triplet type [L H]** | 0.054 | 0.026 – 0.082 | 3.803 | Inf | **< .001** |
| Epoch [4] × Harshness | -0.003 | -0.043 – 0.037 | -0.153 | Inf | .879 |
| Epoch [5] × Harshness | -0.009 | -0.043 – 0.025 | -0.538 | Inf | .591 |
| Epoch [4] × Unpredictability | -0.035 | -0.075 – 0.004 | -1.758 | Inf | .079 |
| Epoch [5] × Unpredictability | 0.027 | -0.007 – 0.061 | 1.573 | Inf | .116 |
| Epoch [4] × Subj cSES | 0.010 | -0.017 – 0.037 | 0.732 | Inf | .464 |

| | | | | | |
|---|---|---|---|---|---|
| Epoch [5] × Subj cSES | 0.001 | -0.022 – 0.024 | 0.065 | Inf | .948 |
| Epoch [4] × Age | -0.000 | -0.007 – 0.007 | -0.021 | Inf | .984 |
| Epoch [5] × Age | -0.002 | -0.008 – 0.004 | -0.566 | Inf | .571 |
| Epoch [4] × Parental education | -0.009 | -0.032 – 0.014 | -0.770 | Inf | .442 |
| Epoch [5] × Parental education | -0.001 | -0.020 – 0.019 | -0.065 | Inf | .948 |
| Triplet type [L H] × Harshness | -0.015 | -0.037 – 0.008 | -1.260 | Inf | .208 |
| Triplet type [L H] × Unpredictability | 0.026 | 0.003 – 0.048 | 2.222 | Inf | **.026** |
| **Triplet type [L H] × Subj cSES** | 0.027 | 0.012 – 0.043 | 3.472 | Inf | **.001** |
| **Triplet type [L H] × Age** | -0.004 | -0.008 – -0.000 | -1.970 | Inf | **.049** |
| Triplet type [L H] × Parental education | 0.009 | -0.005 – 0.022 | 1.275 | Inf | .202 |
| Epoch [4] × Triplet type [L H] × Harshness | 0.008 | -0.025 – 0.041 | 0.495 | Inf | .621 |
| Epoch [5] × Triplet type [L H] × Harshness | -0.006 | -0.036 – 0.024 | -0.371 | Inf | .711 |
| Epoch [4] × Triplet type [L H] × Unpredictability | -0.016 | -0.048 – 0.017 | -0.935 | Inf | .350 |
| Epoch [5] × Triplet type [L H] × Unpredictability | -0.011 | -0.041 – 0.019 | -0.736 | Inf | .462 |
| Epoch [4] × Triplet type [L H] × Subj cSES | 0.014 | -0.008 – 0.037 | 1.245 | Inf | .213 |
| **Epoch [4] × Triplet type [L H] × Subj cSES** | -0.036 | -0.057 – -0.016 | -3.488 | Inf | **< .001** |
| Epoch [4] × Triplet type [L H] × Age | -0.000 | -0.006 – 0.006 | -0.077 | Inf | .939 |
| Epoch [5] × Triplet type [L H] × Age | 0.001 | -0.004 – 0.007 | 0.407 | Inf | .684 |
| Epoch [4] × Triplet type [L H] × Parental education | 0.003 | -0.017 – 0.022 | 0.258 | Inf | .796 |
| Epoch [5] × Triplet type [L H] × Parental education | -0.007 | -0.025 – 0.010 | -0.810 | Inf | .418 |

| **Random Effects** | |
|---|---|
| $\sigma^2$ | 3.290 |

| | |
|---|---|
| $\tau_{00}$ Participant | 0.130 |
| $\tau_{11}$ Participant.Epoch4 | 0.031 |
| $\tau_{11}$ Participant.Epoch5 | 0.017 |
| $\rho_{01}$ Participant.Epoch4 | 0.124 |
| $\rho_{01}$ Participant.Epoch5 | -0.046 |
| ICC | 0.043 |
| N Participant | 325 |
| Observations | 151710 |
| Marginal $R^2$ / Conditional $R^2$ | 0.005 / 0.047 |

**Supplementary Table S16. Results of the binomial mixed model with Parental education on accuracy, rewiring new knowledge.** Top table shows the LRT tests of fixed effects. Bottom table shows regression coefficients of fixed effects and summary information about the random effects. Coefficients are log odds, thus positive values indicate that the given independent variable is associated with an increase likelihood of correct responses, and negative values the opposite. P values for coefficients are from z tests, making the degrees of freedom infinite. The marginal R-squared considers only the variance of the fixed effects, while the conditional R-squared takes both the fixed and random effects into account (based on Nakagawa et al., 2017). Statistically significant terms are highlighted in bold. Terms in brackets indicate the level of factor that is contrasted against the reference level, which is L L triplets for the Triplet type factor and Epoch 6 for the Epoch factor.

Model equation in *lmer* syntax: Correct ~ Epoch*Triplet type*(Harshness + Unpredictability + Subj cSES + Age + Parental education) + (Epoch | Participant)
